# Plant hormone manipulation impacts salt spray tolerance, which preempts herbivory as a driver of local adaptation in the yellow monkeyflower, *Mimulus guttatus*

**DOI:** 10.1101/2024.05.23.595619

**Authors:** Katherine Toll, Megan Blanchard, Anna Scharnagl, Liza M. Holeski, David B. Lowry

## Abstract

A major challenge in evolutionary biology is identifying the selective agents and phenotypes underlying local adaptation. Local adaptation along environmental gradients may be driven by trade-offs in allocation to reproduction, growth, and herbivore resistance. To identify environmental agents of selection and their phenotypic targets, we performed a manipulative field reciprocal transplant experiment with coastal perennial and inland annual ecotypes of the common yellow monkeyflower (*Mimulus guttatus*). We manipulated herbivory with exclosures built in the field and exogenously manipulated gibberellin and jasmonic acid to shift allocation of plant resources among growth, reproduction, and herbivore resistance. Our hormone treatments influenced the timing of allocation to reproduction and allocation to phytochemical defense, but this shift was small relative to ecotype differences in allocation. Herbivore exclosures reduced herbivory and increased fitness of plants at the coastal site. However, this reduction in herbivory did not decrease the homesite advantage of coastal perennials. Unexpectedly, we found that the application of exogenous gibberellin increased mortality due to salt spray at the coastal site for both ecotypes. Our results suggest that divergence in salt spray tolerance, potentially mediated by ecotype differences in gibberellin synthesis or bioactivity, is a strong driver of local adaptation and preempts any impacts of herbivory in coastal habitats that experience salt spray.

## Introduction

Organisms experience dramatically different environmental conditions throughout their geographic ranges. Spatial gradients in abiotic factors, such as temperature, salinity, and water availability, as well as biotic factors, such as the presence of competitors, predators, and mutualists, can generate divergent natural selection (Kawecki & Ebert, 2004; Maron *et al*., 2014; López-Goldar & Agrawal, 2021). This divergent selection can in turn lead to evolutionary responses in traits that increase fitness in local environments, and result in the evolution of local adaptation (Clausen *et al*., 1940; Kawecki & Ebert, 2004; Leimu & Fischer, 2008; Hereford, 2009; Wadgymar *et al*., 2022). Identifying the causal environmental factors contributing to adaptation is a major challenge because environmental conditions often co-vary and thus, experimental manipulations are necessary to identify the environmental agents of selection (Briscoe Runquist *et al*., 2020; Hargreaves *et al*., 2020). Likewise, the phenotypic targets of selection are challenging to identify because traits are often highly correlated, so approaches that minimize trait correlations or manipulate trait variation independently of other traits are necessary to identify adaptive traits (Wadgymar *et al*., 2017, 2022). Despite their importance, experiments that simultaneously manipulate putative environmental selective agents and their phenotypic targets are uncommon (Wadgymar *et al*., 2017, 2022). These field experiments are critical because the interactions of experimental manipulations and environmental factors are complex and can have unexpected outcomes. In this study, we isolate the effect of a putative selective agent, herbivory, at sites that vary in two abiotic factors, salt spray and soil moisture, and manipulate trait variation using hormone applications to identify the environmental and biotic drivers of local adaptation.

Traits that increase fitness on one end of an environmental gradient can reduce fitness on the opposite end of that gradient, resulting in fitness trade-offs (Kawecki & Ebert, 2004; Wadgymar *et al*., 2022). Trade-offs are often directly caused by evolutionary changes in the allocation of limited resources to critical biological functions, including growth, reproduction, and defense (Bazzaz *et al*., 1987; Herms & Mattson, 1992), but may be indirectly caused by interactions with other species (Strauss *et al*., 2002). Theory predicts that resource allocation to herbivore defense should depend on the risk and consequences of herbivory on fitness, and models of the evolution of plant defense assume a cost to the production of herbivore defenses (Rhoades, 1979; Stamp, 2003). Within species, herbivore resistance frequently trades-off with reproduction (Agren & Schemske, 1993; Heil & Baldwin, 2002; Strauss *et al*., 2002; Stowe & Marquis, 2011; Cipollini *et al*., 2017), and increased herbivore resistance is associated with longer growing seasons (Kooyers *et al*., 2017).

Recent studies have shown that shifts in the allocation of resources from rapid growth to herbivore resistance are made through a set of interacting gene networks that are involved in the biosynthesis and downstream signaling of plant hormones (Kazan & Manners, 2012; Huot *et al*., 2014; Campos *et al*., 2016; Havko *et al*., 2016; Aerts *et al*., 2021; Monson *et al*., 2022).

Jasmonic acid (JA) is a key regulatory hormone involved in the response of plants to herbivore attack (Zhang & Turner, 2008; Havko *et al*., 2016). While JA production increases herbivore defense, it also can inhibit rapid plant growth through interactions with other gene networks (Yan *et al*., 2007; Zhang & Turner, 2008; Kazan & Manners, 2012; Yang *et al*., 2012). Further, the antagonistic interactions of genes in the gibberellin and jasmonic acid signalling pathways are thought to contribute to the trade-off in resource allocation in plants between rapid growth/reproduction (promoted by gibberellin) and defense (promoted by jasmonic acid) (Yang *et al*., 2012; Hou *et al*., 2013; Havko *et al*., 2016). However, empirical evidence that evolutionary changes in the GA pathway lead to changes in the relative allocation of resources to reproduction, long-term growth, and herbivore resistance is lacking.

We used locally-adapted ecotypes of the yellow monkeyflower, *Mimulus guttatus* (syn. *Erythranthe guttata*; (Barker *et al*., 2012)) to investigate the mechanisms responsible for the evolution of locally adaptive trade-offs in growth, reproduction, and resistance. In prior reciprocal transplant experiments in this system, the primary environmental factor contributing to local adaptation at inland sites was the onset of summer drought (Hall & Willis, 2006; Lowry *et al*., 2008). In contrast, above-ground factors, potentially salt spray and/or herbivory, contribute to adaptation in coastal habitats (Lowry *et al*., 2009; Popovic & Lowry, 2020). Oceanic salt spray limits plant growth, injures plant tissues, and contributes to coastal plant community zonation through the differential tolerance of species to salt spray (Boyce, 1954). Inland populations of *M. guttatus* are typically small annuals that flower in early spring to avoid the onset of summer drought. Coastal populations, which occur in habitats with year-round soil moisture, are large perennials that allocate resources primarily to long-term vegetative growth (Hall & Willis, 2006; Lowry *et al*., 2008; Hall *et al*., 2010; Baker & Diggle, 2011; Baker *et al*., 2012). Further, coastal populations have higher levels of phytochemical defenses (phenylpropanoid glycosides, PPGs) and are thought to experience higher levels of herbivory than inland annual populations (Holeski *et al*., 2010, 2013; Lowry *et al*., 2019). Although these patterns are suggestive, we lack direct evidence that coastal populations are more resistant to herbivory than annuals when grown in the field. In the greenhouse, coastal populations are more responsive to exogenous applications of gibberellin (GA_3_) than annuals and respond by recapitulating the elongated growth habit of inland annual populations (Lowry *et al*., 2019). As a result, we hypothesize that natural variation in the timing of reproduction, and allocation to vegetative growth and resistance is the result of molecular changes that alter the interactions of the gibberellin (GA) and jasmonic acid (JA) pathways.

In this study, we performed a manipulative reciprocal transplant experiment with coastal perennial and inland annual ecotypes of *M. guttatus* to test whether resource allocation trade-offs contribute to local adaptation at opposite ends of an environmental gradient. We predicted that GA treatment would cause earlier reproduction, that in turn would increase perennial fitness at the inland site, where earlier flowering would rescue fitness for individuals that typically perish before the onset of summer drought. We also predicted that JA and paclobutrazol (a GA inhibitor) treatment would cause increased allocation to vegetative growth (via delayed flowering) and herbivore resistance, that in turn would increase annual fitness at the coast, where we expected herbivore pressure to be higher. Finally, we predicted that reduction of herbivory via exclosures would rescue inland annual fitness on the coast, and thus decrease perennial home site advantage. While our study was designed to focus on the role of hormone manipulation on herbivore resistance, we instead discovered that our hormone manipulations had a much larger role in causing susceptibility to stress imposed by oceanic salt spray. To assess the generality of our observations in the manipulative reciprocal transplant experiment, we performed an additional experiment focusing on the role GA plays in regulating plant height and salt spray sensitivity across multiple populations spanning the coast to inland gradient.

## Materials and Methods

### Study locations and plants used in the reciprocal transplant experiment

In 2020, we performed a reciprocal transplant experiment at a coastal seep at the Bodega Marine Reserve in Bodega Bay, CA (Latitude: 38.3157, longitude: −123.0686), and an inland seep at the Pepperwood Preserve near Santa Rosa, CA (latitude: 38.5755, longitude: −122.7009). We used outbred maternal families from a coastal perennial population that experiences salt spray from Bodega Bay, CA (*n* = 5 families, BHW population coordinates: 38.303783, -123.064483) and an inland annual population north of Sonoma (*n* = 4 families, CAV population coordinates: 38.342817, -122.4854). Since we were primarily interested in ecotype differences, we generated outbred maternal families by intercrossing a small number of field-collected maternal families (*n* = 6 and *n* = 4, for BHW and CAV respectively) in the greenhouses at Michigan State University. Seeds from these outbred families were planted and placed in a 4°C cold room on January 27, 2020 at UC Berkeley. We staggered perennial and annual germination to synchronize their development, following (Popovic & Lowry, 2020). One and two weeks after initiating stratification, perennial and annual seeds were moved into a 16-hr day length growth chamber, respectively. Seedlings were transported from UC Berkeley to the Bodega Marine Reserve (BMR) greenhouse on February 20, 2020, where they experienced ambient light and temperature conditions. In the greenhouse, seedlings were transplanted into 288 cell plug trays filled with Sunshine Mix #4 Professional Growing Mix (Sun Gro Horticulture, Agawam, MA, USA) over a week (February 21- February 28, 2020).

### Hormone treatments applied in the reciprocal transplant experiment

We altered allocation phenotypes of each ecotype by manipulating hormone levels of plants with exogenous applications of gibberellin (GA_3_), paclobutrazol (a gibberellin inhibitor), and methyl jasmonate (MeJA, the bioactive form of Jasmonic Acid) to test the role of those hormone pathways in adaptive trade-offs. Following a week of transplanting in the BMR greenhouse, on February 28, 2020 we randomly assigned trays to one of three hormone treatments or a no-hormone control. We sprayed plants with a 100 µM solution of GA_3_ (*n* = 252 seedlings per ecotype, Consolidated Chemical Solvents LLC, following Lowry *et al*., 2019), 10 mM methyl jasmonate (*n* = 250 seedlings per ecotype; TCI America, Portland, Oregon, USA), or 14.3 mg/L solution of paclobutrazol (*n* = 250 seedlings per ecotype, General Hydroponics, Santa Rosa, California, USA). Concentrations of methyl jasmonate and paclobutrazol were chosen after conducting dose response experiments at the MSU greenhouses in winter 2019, based on the minimum concentration needed to elicit a phenotypic change relative to no-hormone controls without detrimental effects (e.g., leaf damage, stunting, death). Using a spray bottle, we sprayed individual plants 5 times, corresponding to 3.5 mL of solution. The no-hormone control consisted of spraying plants (*n* = 248 seedlings per ecotype) with 3.5mL of a 0.25% ethanol solution since a dilute ethanol solution was needed to dissolve methyl jasmonate and all hormones were dissolved in a 0.25% ethanol solution. Hormones were applied once on a single day in the greenhouse prior to field transplanting. All trays were covered with clear plastic domes for 24 hours (February 28 - February 29, 2020) and moved to different greenhouse benches to prevent cross contamination.

### Field planting of the reciprocal transplant experiment

Prior to transplanting, we removed vegetation from ten (1.08 x 0.84 m) plots at each site. We dug trenches along the edge of each plot to bury the bottom of our open control and exclosure structures. At the Pepperwood Preserve, we transplanted 800 seedlings on March 9, 2020, and 200 seedlings on March 12, 2020. At the Bodega Marine Reserve, we transplanted 800 seedlings on March 10, 2020, and 200 seedlings on March 13, 2020. All seedlings had two expanded leaf pairs at transplanting. Plants were fully randomized within each block (*n* = 100 seedlings/block).

### Herbivore exclosures used in the reciprocal transplant experiment

To lessen the effect of herbivory, we deployed herbivory exclosures on four out of ten plots at each site (Figure 1). A previous reciprocal transplant experiment at our study sites used agrofabric exclosures that blocked all above-ground factors (Popovic & Lowry, 2020), and could not separate the effects of salt spray and herbivory on plant fitness. Thus, we designed exclosures that excluded many herbivores but allowed salt spray to pass through. The exclosures were 1.08 m long x 0.84 m deep x 0.87 m tall and constructed of 2.67 cm diameter PVC pipe covered with fiberglass window-screen (1.4 x 1.6 mm mesh) that was affixed with fishing line and marine epoxy. Each exclosure had screen doors along both long sides that were attached with velcro to allow access to the plots. The screen extended 10 cm down into the soil around the plots. To control for shading or moisture-collection due to the screen, the remaining six plots (open structures) at each site were covered with window screen only on the top of each structure and 30 cm down each side.

**Figure 1.**
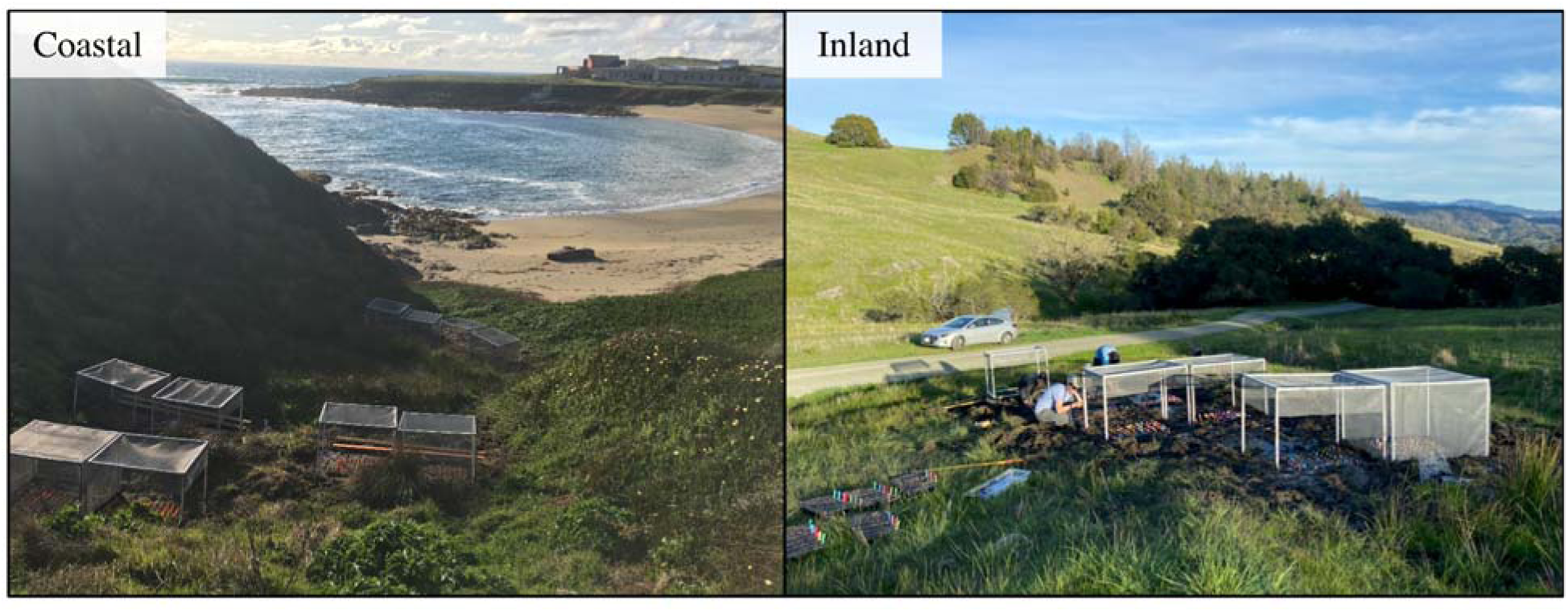
Photographs depicting transplant sites and structures used in the reciprocal transplant experiment. For our reciprocal transplant experiment, each site had ten plots, each with 100 plants. Each site had six control plots (with mesh tops and partially-mesh sides) and four exclosure plots (fully enclosed with mesh on all sides). A control and exclosure plot are shown side by side in the foreground of the inland site.

### Field Censuses of the reciprocal transplant experiment

After transplanting seedlings at the two-leaf stage, we performed regular censuses of our transplant sites, recording survival, the presence of herbivore damage, the presence of salt-spray damage, and the presence and number of reproductive structures (buds, flowers, and fruits). In our census, we distinguished damage and death caused by salt spray from herbivory: salt- damaged leaves appeared necrotic and brown and exhibited no sign of herbivore damage (i.e., no missing tissue). When salt damage spread to the entire plant and no green tissue remained, we considered plants to be killed by salt spray. When possible, we distinguished between sources of herbivory. Most notably, we observed damage to plants consistent with mammalian damage at the coastal site (removal of flowering stalks leaving plant stumps) that we could visually distinguish from insect chewing damage and leaf miners. We were prevented from accessing our transplant sites for two weeks at Bodega Marine Reserve and seven weeks at Pepperwood Preserve after transplanting due to the 2020 COVID-19 pandemic lockdowns. Due to site differences in growing season length, and restricted access due to the 2020 COVID-19 pandemic, we censused each site at different intervals and for different lengths of time (Pepperwood Preserve (inland site): 12 censuses over 139 days, April 3 through July 29, 2020; Bodega Marine Reserve (coastal site): 22 censuses over 194 days, March 24 through September 22, 2020). Since we were unable to access our sites for weeks because of the pandemic, we missed observing the first flower opening for many annual plants at the inland transplant site. For these plants, we estimated the onset of flowering as the date we first observed any reproductive structures. Our censuses occurred on a roughly weekly basis after we were able to re-access our sites.

### Plant chemistry measured in the reciprocal transplant experiment

We sampled leaves for chemical analysis 55 to 57 (May 7-9, 2020) and 59 to 64 (May 11-16, 2020) days after transplanting at the coastal and inland site, respectively. To minimize the potential effect of diurnal fluctuation in PPGs (phenylpropanoid glycosides), we sampled from 9am until 1pm, and to minimize the effect of leaf position, we sampled two leaves from the 3rd node when possible, using leaves from the 4th and 5th nodes if leaves at the 3rd node were damaged. After sampling, leaves were flash-frozen with liquid nitrogen and then freeze-dried. For samples that did not meet the minimum dry mass (3mg), we either pooled them with other low-mass samples (by grouping within all fixed and random factors as discussed below) or excluded them. Our final sample size was 216 from Bodega (perennials only due to high annual mortality) and 599 from Pepperwood. To determine the PPG concentrations in sampled leaves, we ground, extracted, and prepped extract aliquots as described in Holeski *et al*. (2013). We then used high-performance liquid chromatography (HPLC) to quantify PPGs. The HPLC method is described in (Kooyers *et al*., 2017) and was run on an Agilent 1260 HPLC with a diode array detector and Poroshell 120 EC-C18 analytical column [4.6 × 250 mm, 2.7 μm particle size]; Agilent Technologies). We calculated concentrations of individual PPGs as verbascoside equivalents, using a standard verbascoside solution (Santa Cruz Biotechnology, Dallas, Texas), as described in (Holeski *et al*., 2013, 2014).

### GA_3_-salt spray exposure experiment (2023)

Since we observed a large effect of GA_3_ treatment on plant height (although we did not measure it), necrosis and mortality at our coastal transplant site in 2020, we wanted to confirm the combined impact of GA_3_ and salt spray and assess how common this effect was across multiple populations. In 2023, we grew one outbred accession from four coastal perennial (HEC, OPB, SWB, BMR) and four inland annual (LMC, PPW, CAV, MOR), one inland perennial (SAL), one near-coastal annual (AJEN, 1.5km from the coast), and two near-coastal perennial populations (JEN and GUL, 3 and 4.7 km from the coast respectively) in a greenhouse in the Bodega Marine Reserve (Table S17). Near-coastal populations experience the same maritime climate as coastal populations but are not directly exposed to salt spray. Seeds for perennials were planted a week earlier than annuals to synchronize germination (March 2 vs March 9, 2023). Seedlings were transplanted into individual 8.9 cm square pots (T.O. Plastics, Clearwater, MN, USA) as they germinated and randomly assigned to a GA_3_ treatment or no-hormone control. We used the exact same protocol as in 2020 for the GA_3_ treatment and no-hormone control, except that we did not add ethanol to the spray bottle which was used as a solvent for methyl jasmonate. A total of 22 and 23 seedlings per population were sprayed with GA_3_ and water, respectively, on March 24, 2023. GA_3_-treated and no-hormone control plants were sprayed on separate greenhouse benches and covered with clear plastic domes for 24 hours to prevent cross contamination. Two weeks after the GA_3_ or water treatments, we randomly assigned seedlings into blocks (six 1020 trays [T.O. Plastics, Clearwater, MN, USA] per block). We measured height from the base of each plant to the apical meristem using a ruler in the greenhouse on April 19, 2023 and then immediately placed plants (*n* = 540) outdoors on a coastal bluff for one week (latitude: 38.31841, longitude: -123.07329; Figure S1). On the bluff, we either protected the plants from salt spray using two agrofabric covered exclosures (*n* = 218 plants) or exposed them to salt spray using three open structures (*n* = 324 plants) using the same structure dimensions and agrofabric as (Popovic & Lowry, 2020). The two agrofabric exclosures contained ten GA_3_-treated and eight water-sprayed plants per population, and the three open structures contained 12 GA_3_-treated and 15 water-sprayed plants per population. After one week of exposure, we transported plants back to the greenhouse and assessed levels of leaf damage on each leaf pair by eye. Standard methods for estimating whole plant damage typically break up whole plants into units (Herbivory Variability Network*† *et al*., 2023). Leaf pair was chosen as the unit to visually estimate damage because monkeyflowers produce alternate leaves at each node. Whole plant damage was estimated by averaging visual estimates of leaf damage for each leaf pair within each plant.

Because plants were maintained in pots during the entirety of the 2023 experiment, we could be assured that damage was caused by oceanic salt spray applied to experimental plants by wind coming off of the Pacific Ocean. We did not observe any herbivore damage to plants exposed or protected from the ocean in 2023.

### Statistical analyses

The overarching goal of our statistical analyses was to understand how ecotype, hormone treatment, field site, and herbivore exclosure treatment contribute to within plant resource allocation and fitness. To establish how plants with different ecotype and treatment combinations allocate resources to reproduction, we compared flowering time among plants. To compare allocation to herbivore resistance, we compared the presence or absence of herbivore damage to each plant and the concentration and composition of the defensive compounds in leaves. To compare fitness, we quantified survival across the season, the presence of flowers, and seasonal fruit production. For annuals, these measures indicate lifetime fitness, whereas perennials that survived the season have the potential to reproduce in subsequent years. We performed all statistical analysis in R version 4.3.1 (R core team 2023).

#### Univariate analysis

To test whether ecotypes differed in resource allocation and whether hormone treatments altered these differences at each transplant site, we fit mixed models for each analysis that included ecotype, hormone treatment, and exclosure type as interactive fixed factors. The response variables were flowering time, the presence or absence of herbivory, survival, total PPG concentration (summed concentration for all PPG compounds), whether an individual produced a reproductive structure (buds, flowers, or fruit), or the number of fruits produced by plants that flowered at the end of the season. When possible we categorized herbivore damage by the putative source, based on patterns of plant tissue damage. Methods and results of analyses by herbivore type are detailed in the supplement. All models also included two random effects for maternal family and experimental plot. We fit all mixed models except for the survival model with the R package *glmmTMB* (Brooks *et al*., 2017), and modeled survival using the R package *coxme* (Therneau, 2022). We identified the best fitting error distributions by evaluating model diagnostics with the R package *DHARMa* (Hartig, 2022). We fit mixed models for flowering time with Gaussian error distributions, mixed models for herbivory and flowering probability with binomial error distributions, and mixed models for log-transformed total PPGs with gamma distributions. We modeled survival using a mixed-effects Cox Proportional Hazards model, and modeled fruit number with a zero-inflated negative binomial error distribution at the coastal site and a negative binomial error distribution at the inland site.

To prevent model overfitting, we used an analysis of deviance (Wald ^2^ test) to assess the significance of model terms and sequentially removed unsupported model terms (R package *car*, (Fox & Weisberg, 2018). We compared fits of complex versus reduced models using likelihood ratio tests (LRT) to find the minimum adequate model for each response variable in each site (Tables S1 and S2). We compared treatment groups using post-hoc tests on the minimum adequate model with the R package *emmeans* (Lenth *et al*., 2020). For one of the binary response variables (flowering probability), there was no variance in one of the treatment levels (quasi-complete separation) and a binomial model was not able to accurately estimate the parameter for that treatment group. For this reason, we used a randomization test to estimate the probability that this result was due to chance, by reassigning treatments to all outcomes 10,000 times and calculating the proportion of these reshuffles that resulted in a difference as extreme as the observed (Gotelli & Ellison, 2004). We predicted the mean and 95% confidence intervals for each response variable from our models using the R package *ggeffects* (Lüdecke, 2018). For all non-binary response variables, we predicted confidence intervals via bootstrapping (*n*=500 iterations). We plotted raw data and predictions in *ggplot2* (Wickham, 2016) and combined plots with *patchwork* in R (Pedersen, 2019).

#### Multivariate PPG analysis

Within each transplant site, we modeled the concentration of all nine different PPGs (the PPG arsenal) using mixed models for each analysis that included ecotype (at Pepperwood only), hormone treatment, and exclosure type as interactive fixed factors and block as a random factor. We fit all models with PERMANOVA with Bray-Curtis distance using the adonis2 function from the R package *vegan* (Oksanen, 2016). We dropped all non-significant factors for the minimum adequate model (Table S3). To test for homogeneity of variance among treatment groups, which can influence inference, we used the betadisper function from the *vegan* package. The only factor that had heterogeneity of variance among levels was ecotype. Due to the strength of the signal for ecotype, and confirmation from other studies that annuals and perennials have different PPG arsenals (Holeski *et al*., 2013), we are confident that differences due to ecotype are not attributable only to heterogeneity of variance. We compared treatment groups using post-hoc tests on the minimum adequate model with the function pairwise.adonis2 from the R package *pairwiseAdonis* (Arbizu, 2019). To visualize how multivariate PPG arsenals are influenced by our factors, we used non-metric multidimensional scaling (NMDS) (*MetaMDS* function in *vegan* package with Bray-Curtis distance to determine dissimilarity) and added standard-error ellipses at 95% confidence around the centroid of each cluster (function *ordiellipse* from package vegan).

#### Analysis of GA_3_-salt spray exposure experiment (2023)

To analyze the effect of GA_3_ treatment on plant height in the greenhouse, we fit a linear mixed model with height as the response variable and population (*n* = 12), GA_3_ treatment (*n* = 2), and their interaction as the predictor variables and flat (*n* = 30) as a random effect. Since only 10% (21/216) of seedlings in the exclosures showed any evidence of necrosis after one week of being outdoors, compared to 78% (254/324) of seedlings in the open structures, we only analyzed the effect of hormone treatment and population on plants exposed to salt spray in the three open structures. We first logit transformed proportional leaf necrosis (Warton & Hui, 2011), and then fit a linear mixed model with logit-transformed leaf necrosis as the response variable, population, hormone treatment, and their interaction as fixed effects, and flat (*n* = 18) nested within block (*n* = 3) as random effects. Within each population, we performed pairwise comparisons of no-hormone control vs GA_3_-treated plants (function *pairs* from package emmeans). We used a multivariate *t*-adjustment for multiple comparisons (*n* = 12 comparisons). We plotted predicted leaf necrosis by transforming logit-transformed proportions back to proportions.

## Results

### Annuals flowered earlier, investing less time in vegetative growth than perennials, and hormone treatments slightly delayed flowering time

At the coastal site, flowering time was significantly associated with ecotype, hormone treatment, and an ecotype by hormone treatment interaction (Table S2: all *p* ≤ 0.03). At the inland site, flowering time was significantly associated with ecotype, hormone treatment, an ecotype by hormone treatment interaction, and a three-way ecotype by hormone treatment by exclosure treatment interaction (Table S2: all *p* ≤ 0.01). Annuals flowered 37-62 and 47-63 days earlier than perennials at the inland and coastal sites, respectively (within each hormone and exclosure treatment), respectively (Tables S4 & S5: all annual vs perennial comparison *p*-values < 0.001, Figure 2 A&B). Annuals treated with GA_3_ flowered 9 and 16 days later than annuals not treated with hormones at the inland site in open structures and at the coastal site, respectively (Tables S4 and S5, coastal site: GA_3_ vs control annuals *p* = 0.043; inland site: GA_3_ vs no-hormone control annuals in open structures *p* < 0.009, in exclosures *p* = 1). Annuals treated with paclobutrazol at the coastal site flowered 6 days later than annuals not treated with hormones (Tables S4 and S5, coastal site: paclobutrazol vs no-hormone control annuals *p* = 0.027). Mesh exclosure treatment did not influence flowering time (Table S1: *p* = 0.66; Table S5: all comparisons between exclosure treatments within ecotype and hormone treatment at the inland site *p* ≥ 0.463).

**Figure 2.**
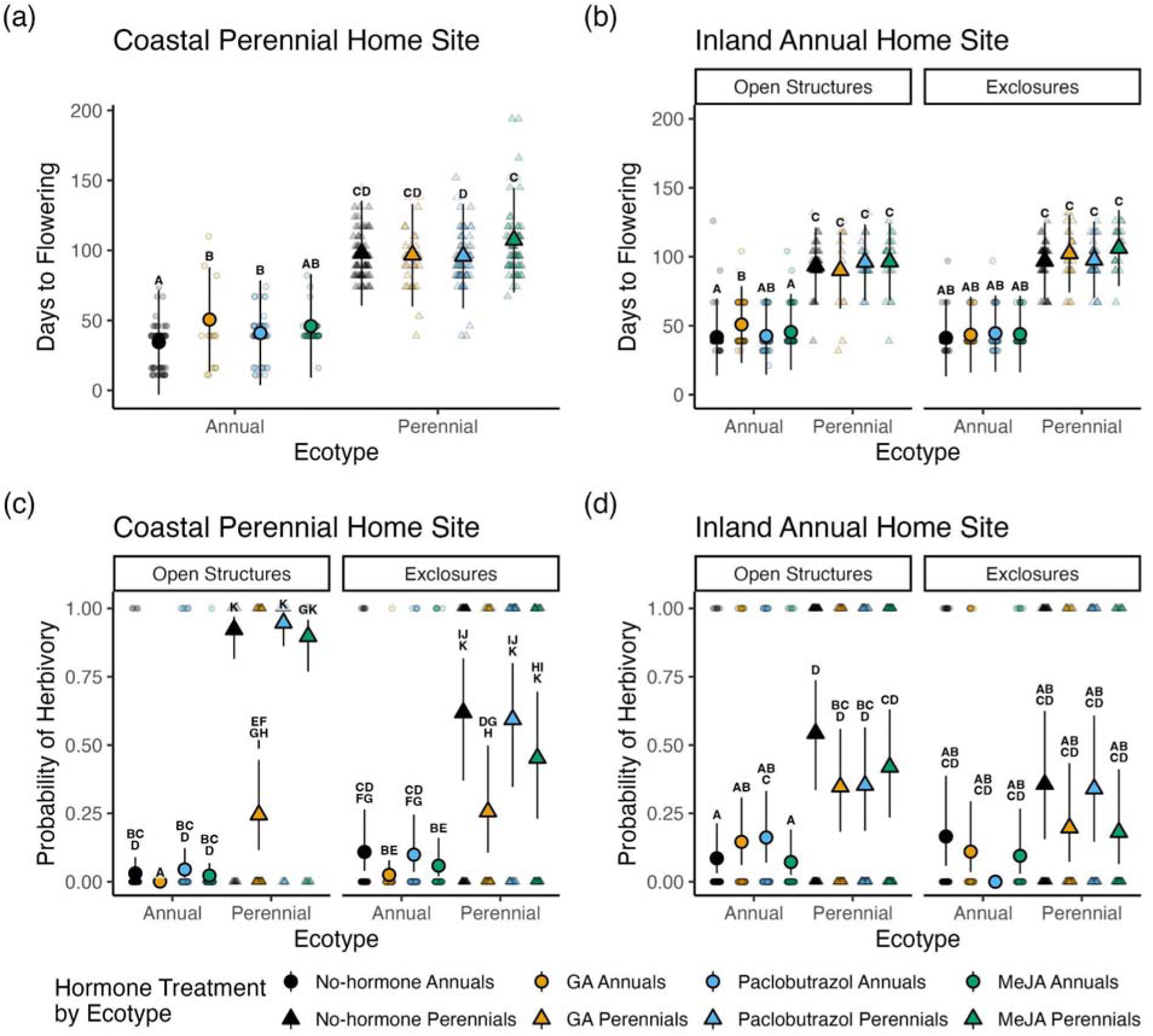
Flowering time and probability of herbivory of annuals (circles) and perennials (triangles) treated with gibberellic acid (GA, yellow), paclobutrazol (blue), and methyl jasmonate (MeJA, green), and the no-hormone controls (black) in open structures and herbivore exclosures at the coastal site, Bodega Marine Reserve (A & C), and the inland site, Pepperwood Preserve (B & D). Larger symbols in the foreground are the mean predictions and 95% confidence intervals from the minimum adequate mixed models, smaller and lighter symbols in the background are the raw data. Since no annuals treated with paclobutrazol in exclosures at the inland site were damaged by herbivores (d), the binomial model did not accurately estimate that parameter and thus we did not plot those confidence intervals (0-100%). Results of Tukey post-hoc contrasts within each site are indicated above each prediction; shared letters indicate that groups do not significantly differ, while non-overlapping letters indicate that groups significantly differ within each site. Exclosure type was not plotted for flowering time on the coast (A) because the minimum adequate model did not include exclosure type as a fixed effect.

Detailed results can be found in the supplemental materials.

### Perennials were more likely to be attacked by herbivores at both sites and GA_3_ reduced the probability of herbivory at the coastal site

At the coastal site, the probability of herbivory was significantly associated with ecotype, mesh exclosure treatment, an ecotype by exclosure treatment interaction, and a hormone treatment by exclosure treatment interaction (all *p* < 0.001, Table S2). At the inland site, the probability of herbivory was significantly associated with an ecotype by hormone treatment interaction, and an ecotype by exclosure treatment interaction (all *p* ≤ 0.03, Table S2). Contrary to expectations, perennials were 21-287 and 6-10 times more likely to experience herbivory than annuals at the coastal site in open structures and exclosures, respectively (all annual vs perennial comparisons within hormone and exclosure treatment *p* < 0.001, Table S6, Figure 2 C). Perennials were 6 times more likely to experience herbivory than annuals in open structures when treated with MeJA or no hormone at the inland site (no-hormone and MeJA-treated annual vs perennial comparisons in open structures *p* < 0.001, all other within hormone and exclosure treatment contrasts *p* ≥ 0.28; Table S7, Figure 2D). However, this result is somewhat misleading at the coastal site, since annuals perished quickly and had less time to encounter herbivores and accrue herbivory. GA_3_ was the only hormone to influence the probability of herbivory, and only at the coastal site, where, again contrary to expectations, it reduced the probability of herbivory for both ecotypes. Annuals and perennials were 77-97% and 59-73% less likely to experience herbivory when treated with GA_3_ relative to no-hormone controls (all no-hormone control vs GA_3_-treatment contrasts within each ecotype and exclosure treatment *p* < 0.001, Table S6, Figure 2C). This is potentially due to an interaction with salt-spray resistance rather than allocation to herbivore resistance.

The mesh-size of screen used in our exclosures, while necessary to allow salt spray to enter, did allow some small insects, including leaf miners and weevils, to enter the exclosures (or they were present when the exclosures were erected) and damage plants mildly. As a result, our herbivore exclosures did not significantly reduce the probability of herbivory for perennials at either site (Table S6 & S7: all open vs exclosure structure contrasts within each hormone treatment for perennials *p* ≥ 0.06, Figure 2C). However, we observed the removal of flowering stalks from perennial transplants consistent with mammalian damage (likely deer, based on the nature of the cuts, hoof prints, and observations of the resident deer population, per obs.) only in open structures at the coastal site. Exclosures significantly reduced deer herbivory by 100%, but did not significantly reduce other sources of herbivory for coastal perennials at the coastal site (detailed in the supplemental materials, Figure S2). At the coastal site, GA_3_-treated annuals were 29% more likely to experience herbivory in exclosures compared to open structures, although this is likely due to an interaction with salt-spray (Table S6: *p* = 0.02, all other open vs exclosure structure contrasts within hormone treatments for annuals *p* = 1, Figure 2C). At the inland site, hormone-treated annuals and perennials did not significantly differ from their respective no-hormone controls and exclosure treatment did not influence the probability of herbivory for either ecotype (Table S7: all open vs exclosure structure contrasts within ecotype and hormone treatment *p* ≥ 0.96, Figure 3D).

**Figure 3.**
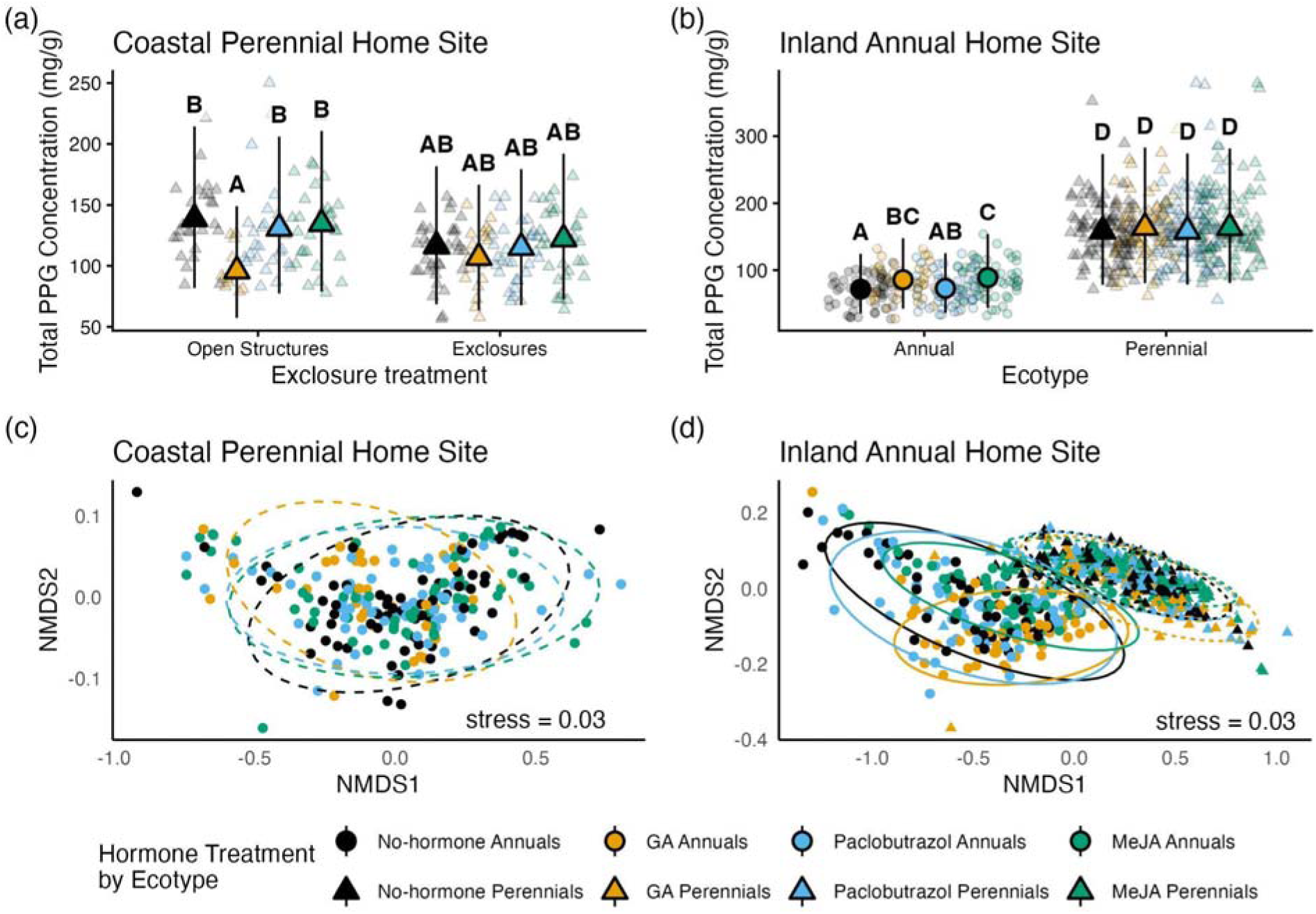
Allocation to chemical defense: total concentration of all PPGs and differences in multivariate PPG arsenals of annuals (circles) and perennials (triangles) treated with gibberellic acid (GA, yellow), paclobutrazol (blue), and methyl jasmonate (MeJA, green), and the no- hormone controls (black) in open structures and herbivore exclosures at the coastal site, Bodega Marine Reserve (A & C), and the inland site, Pepperwood Preserve (B & D). In the Total PPG figures (A & B), larger symbols in the foreground are the mean predictions and 95% confidence intervals from the minimum adequate mixed models, smaller and lighter symbols in the background are the raw data. Exclosure type was not plotted for Total PPG at the inland site (B) because the minimum adequate model did not include exclosure type as a fixed effect. PPG arsenal figures (C & D) use non-metric multidimensional scaling (NMDS) with Bray-curtis distance (and 95% confidence interval ellipses) to visualize multivariate differences among plants. Exclosure type was either not a significant factor in the multivariate model (C) or did not interact with other fixed effects (D) and was therefore not included in these plots.

### Perennials produced more PPGs, and methyl jasmonate and GA_3_ increased PPGs at the inland site while GA_3_ decreased PPGs at the coastal site

At the coastal site, total PPG concentrations were significantly associated with hormone treatment, and a hormone treatment by mesh exclosure treatment interaction (Table S2: all *p* ≤ 0.008). At the inland site, total PPG concentrations were significantly associated with ecotype, hormone treatment, and an ecotype by hormone treatment interaction (Table S2: all *p* ≤ 0.01). Perennials produced 75-87 mg/g more PPGs than annuals at the inland site (Table S9: all annual vs perennial comparisons within hormone treatments *p* < 0.001, Figure 3B), while annual mortality at the coast prevented this comparison (Figure 3A). Exclosure treatment did not significantly influence total PPG concentrations at either transplant site (Table S8: all open vs exclosure structure comparisons within hormone treatment at the coastal transplant site *p* ≥ 0.76, Table S2: *p* = 0.67). At the coastal site in the open structures, perennials treated with GA_3_ produced 43 mg/g fewer PPGs than perennials not treated with hormones (Table S8: no-hormone control vs GA_3_ in open structures: *p* < 0.001, in exclosures: *p* = 0.76). At the inland site, annuals and perennials treated with GA_3_ produced 14 mg/g and 5 mg/g more PPGs than annuals and perennials not treated with hormones, respectively (Table S9: no-hormone control vs GA_3_- treatment comparisons for each ecotype: *p* < 0.001). Annuals treated with methyl jasmonate produced 17 mg/g more PPGs than annuals not treated with hormones at the inland field site (Table S9: no-hormone control vs MeJA-treatment contrast *p* = 0.001 for annuals, *p* = 0.99 for perennials).

At the coastal site, multivariate PPG arsenals were significantly associated with hormone treatment; GA_3_-treated perennials differed from all other perennials (Table S3: all *p* ≤ 0.02, Figure 3C). At the inland site, multivariate PPG arsenals were significantly associated with ecotype, hormone treatment, mesh exclosure treatment, and an ecotype by hormone interaction (Table S3: all *p* < 0.003, Figure 3D). Detailed results can be found in the supplemental materials.

### Perennials survived longer at both sites, and GA_3_ reduced survival at the coastal site

At the coastal transplant site, survival was significantly associated with ecotype, hormone treatment, mesh exclosure treatment, an ecotype by hormone treatment interaction, and a three-way ecotype by hormone treatment by exclosure treatment interaction (Table S2: all *p* ≤ 0.04). At the inland transplant site, survival was only significantly associated with ecotype (Table S2: *p* < 0.001). At both transplant sites, perennials survived significantly longer than annuals (Tables S12 & S13: all annual vs perennial comparisons [within hormone and exclosure treatment at the coastal site] *p* < 0.001, Figure 3A&B). At the inland site, the median death date for annuals (97 days from transplanting) was 21 days earlier than perennials (118 days). At the coastal site, the median death date for annuals was 46 days, but since fewer than 50% of perennials before the end of our experiment, we could not estimate their median death date. At the coastal site, we observed tissue necrosis prior to the death of all annuals and 93% (109/117) of the perennials that died. For the remaining 8 perennial plants, we did not observe tissue necrosis but did observe herbivory prior to death. The majority of plants at the inland site died as the site began to dry out in summer.

At the coastal site, GA_3_ treatment significantly increased the risk of death for both ecotypes relative to their respective no-hormone controls (GA_3_:no-hormone cumulative hazard ratios - annuals: 2.65 [open structure], 3.34 [exclosure], perennials: 28.6 [open structure], 9.87 [exclosure]; Table S12: no-hormone control vs GA_3_-treatment comparisons for both ecotypes, *p* < 0.001). GA_3_-treated plants senesced and died faster in the open structures at the coastal site; the median death date in the open structures was 53 days compared to 96 days in the exclosures (Figure 4A). We observed tissue necrosis on GA_3_-treated plants prior to death at the coastal site. GA_3_-treated perennials were more upright compared to prostrate no-hormone controls, and elongated their stems early in development like annuals which may have increased exposure to salt spray (Figure 5). Mesh herbivore exclosures decreased the risk of death in GA_3_-treated perennials at the coastal site (open structure:exclosure cumulative hazard ratio: 2.53, Table S12: open vs exclosure structure comparison *p* < 0.001, Figure 4A). Coastal fog sometimes condensed on the screens that we used to exclude herbivores, which may have slightly decreased the transmission of salt spray into exclosures.

**Figure 4.**
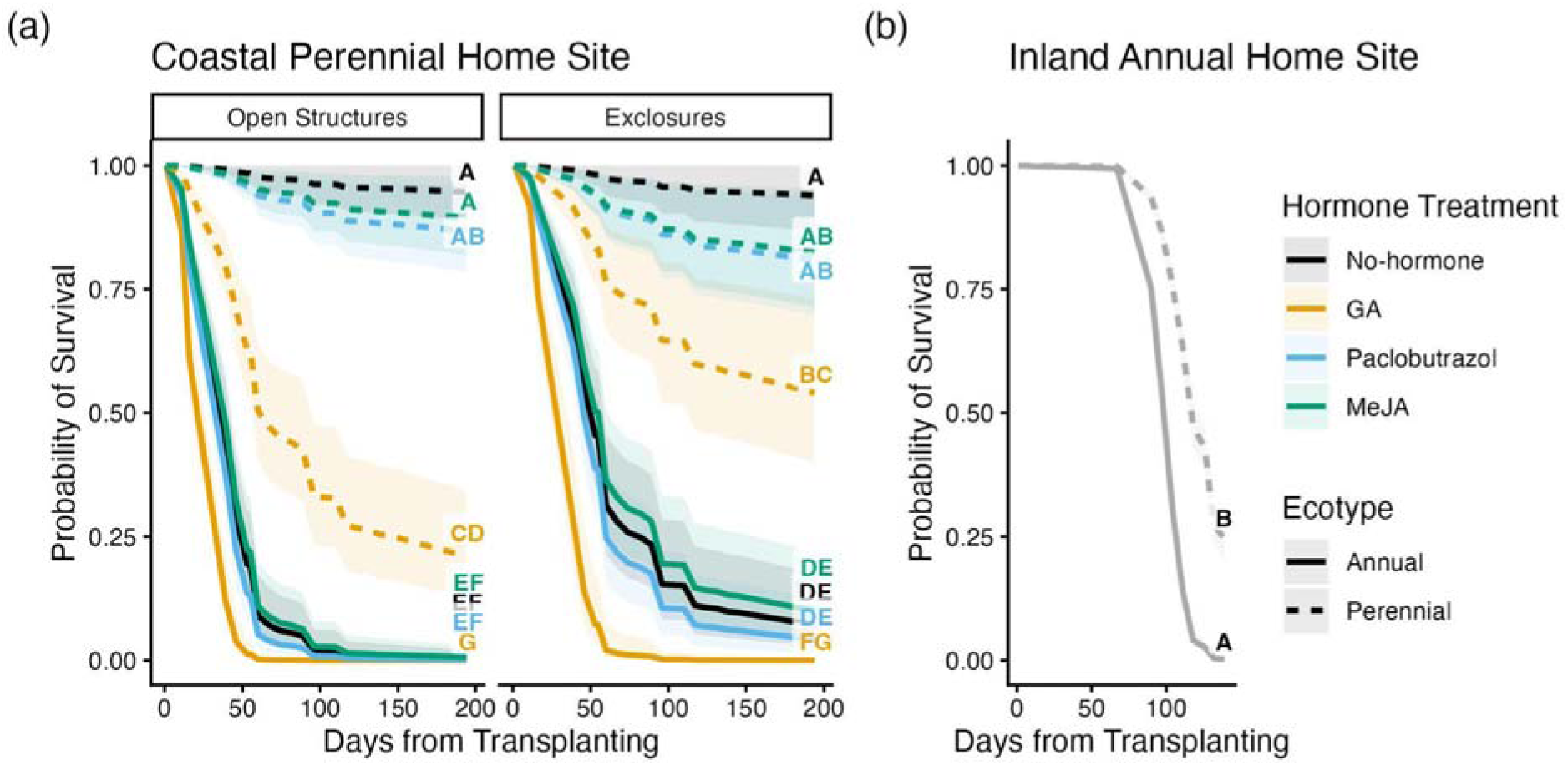
GA_3_ application decreased survival at the coastal site. Survival probabilities for annual (solid line) and perennial (dashed line) transplants at the coastal site, Bodega Marine Reserve (A), and the inland site, Pepperwood Preserve (B). Survival probabilities and 95% confidence intervals for control (black), GA (yellow), methyl jasmonate (green) and paclobutrazol (blue) treatments were predicted from Cox Proportional Hazards models. At the inland transplant site (B), survival probabilities and 95% confidence intervals were only plotted for ecotypes (grey) because the minimum adequate model did not include exclosure or hormone treatment as a fixed effect. Results of Tukey post-hoc contrasts within each site are indicated above each final predicted survival; shared letters indicate that groups do not significantly differ, while non- overlapping letters indicate that groups significantly differ within each site.

**Figure 5.**
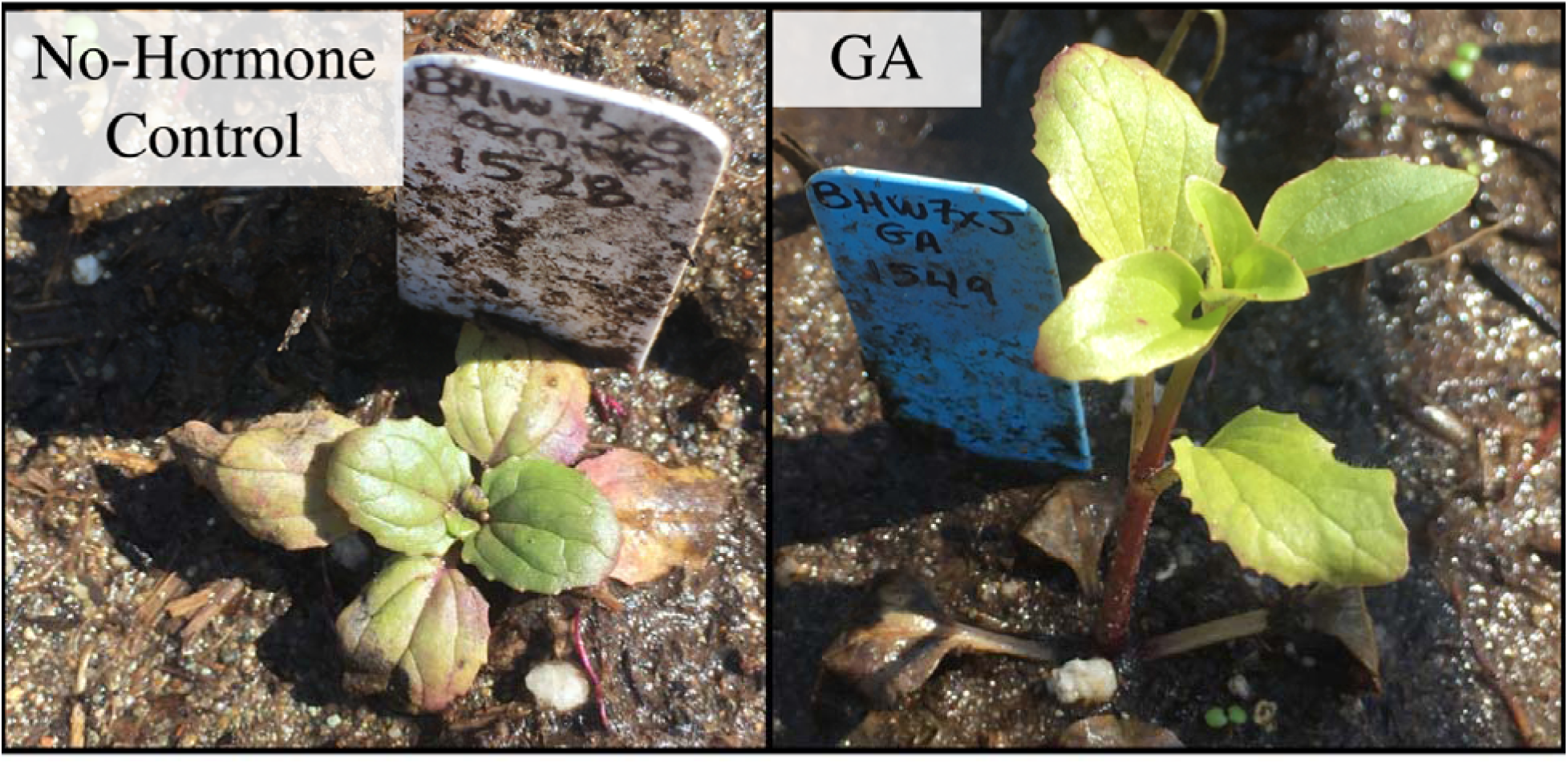
GA_3_ application on perennials resulted in stem-elongation relative to controls. The plants pictured (on day 16 after transplantation) are from the same family and were grown in the same plot at the coastal site.

### Annuals were more likely to flower at the inland site, and GA_3_ reduced flowering probability at the coastal site

At the coastal site, the probability of flowering was significantly associated with hormone treatment, mesh exclosure treatment, and an ecotype by exclosure treatment interaction (Table S2: all *p* ≤ 0.007). Despite high early season annual mortality at the coastal site, annuals and perennials did not significantly differ in the probability of flowering (Table S14: all annual vs perennial comparisons within exclosure treatment *p* ≥ 0.18, Figure 6A). Due to their rapid phenology, 40% (198/500) of annual transplants flowered prior to dying, although death occurred quickly after flowering so very few annuals produced fruit (detailed below). Perennials were 3-11 more likely to flower in herbivore exclosures than open structures (Table S14: *p* < 0.014 for open vs exclosure structure contrasts, Figure 6A), which may be due to the reduction of large mammalian herbivory and/or minor buffering of salt spray from condensation collecting on mesh screens. In addition, transplants that were not treated with hormones were more likely to flower than transplants treated with GA_3_ (8x in open structures, 3-4x in exclosures), potentially due to the effect of GA_3_ on sensitivity to salt spray (Table S14: all no-hormone control vs GA_3_- treatment contrasts *p* < 0.001, Figure 6A).

**Figure 6.**
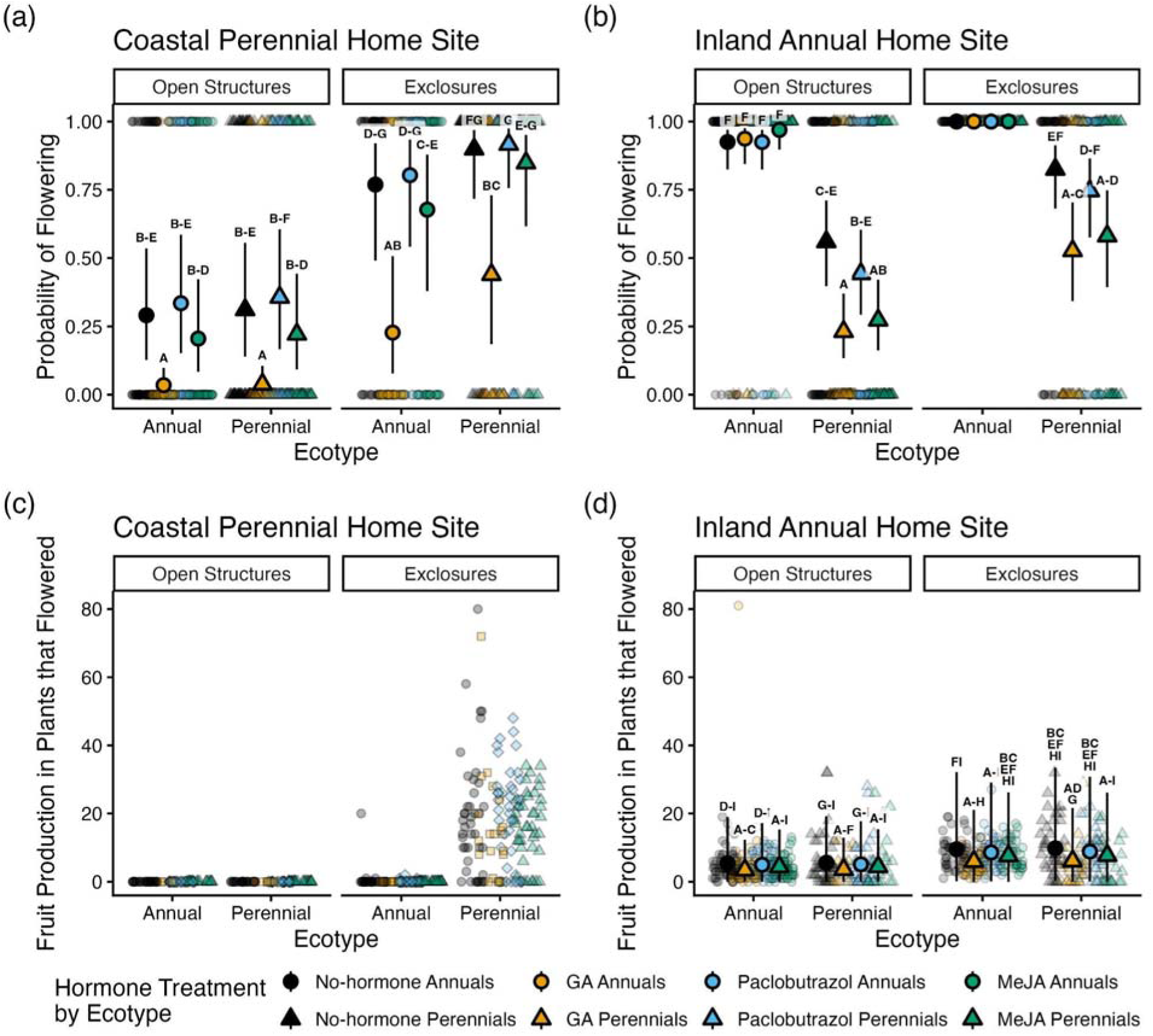
Herbivore exclosures tended to increase, while GA_3_ applications tended to decrease components of fecundity: probability of flowering and fruit number of annuals (circles) and perennials (triangles) that flowered treated with gibberellic acid (GA, yellow), paclobutrazol (blue), and methyl jasmonate (MeJA, green), and the no-hormone controls (black circles) in open structures and herbivore exclosures at the coastal site, Bodega Marine Reserve (A & C), and the inland site, Pepperwood Preserve (B & D). Larger symbols in the foreground are the mean predictions and 95% confidence intervals from mixed models, smaller and lighter symbols in the background are the raw data. Since all annuals flowered in exclosures at the inland site (b), the binomial model did not accurately estimate that parameter and thus we did not plot those confidence intervals (0-100%). Results of Tukey post-hoc contrasts within each site are indicated above each prediction; shared letters indicate that groups do not significantly differ, while non- overlapping letters indicate that groups significantly differ within each site. Predictions are not plotted for fruit production at the coastal site because no fixed factors were in the minimum adequate model. To improve visualization, one outlier that produced 164 fruit was not plotted at the coastal site.

At the inland transplant site, the probability of flowering was significantly associated with ecotype and an ecotype by hormone treatment interaction (Table S2: both *p* ≤ 0.01). At the inland site, annuals were 65-305% and 34-90% more likely to flower than perennials in open structures and exclosures, respectively (Table S15: all annual vs perennial comparisons within each hormone treatment in open structures *p* < 0.001, in exclosures *p* = 1, Figure 6B). Since all annuals flowered in the exclosures, the binomial model could not accurately estimate flowering probabilities for annuals in exclosures. Therefore, we used a randomization test to estimate the probability that the difference between annuals and perennials in exclosures was due to chance. Randomization tests showed that 0 (all hormone treatments) or 15 (the no-hormone controls) of 10,000 treatment reshuffles resulted in the observed difference between annual and perennial flowering probabilities in the exclosures, making it unlikely that this pattern arose by chance (no-hormone controls: *p* = 0.0015, GA-, methyl jasmonate- and paclobutrazol- treated plants *p* = 0.0). Perennials treated with GA_3_ and MeJA were 30-51% and 36-59% less likely to flower than perennials not treated with hormones, but hormone treatment did not affect the probability of flowering in annuals (Table S15: *p* ≤ 0.002 for no-hormone control vs GA_3_ and no-hormone control vs MeJA-treatment contrasts in perennials within each exclosure treatment, Figure 6B).

### GA_3_ reduced fruit production at the inland site, and herbivore exclosures drastically increased fruit production at the coastal site

At the inland site, fruit number among plants that flowered was significantly associated with hormone treatment and marginally associated with mesh exclosure treatment (Table S2: *p* < 0.001 and *p* = 0.06). Annuals and perennials that flowered did not significantly differ in fruit production in either open structures or exclosures (Table S16: all annual vs perennial comparisons *p* =1, Figure 6D). Each ecotype produced 2-4 fewer fruits when treated with GA_3_ relative to no-hormone controls (Table S16: all no-hormone control vs GA_3_-treatment comparisons *p* < 0.001, Figure 6).

At the coastal site, only plants protected by mesh exclosures successfully produced fruit by the end of the season (Figure 6C) because of complete inflorescence herbivory by mule deer (*Odocoileus hemionus*). Since no plants produced fruit outside of the exclosures, and few annuals produced fruit inside the exclosures, we analyzed only the effect of hormone treatments on perennial fruit production inside the exclosures at the coastal site (Figure 6C). Fruit production in perennial plants that flowered in exclosures at the coastal site was not significantly associated with hormone treatment (Table S1: *p* = 0.48).

### GA_3_ increased plant height and salt spray susceptibility

In the subsequent experiment in 2023, greenhouse plant height was significantly associated with population (Analysis of deviance: Wald Type III χ^2^= 628.78, *df* =11, *p* < 0.001), GA_3_ treatment (χ^2^= 1318.39, *df* =1, *p* < 0.001), and a population by GA_3_ treatment interaction (χ^2^= 362.10, *df* =11, *p* < 0.001). In pairwise comparisons, GA_3_ treated plants from 12 out of 13 populations were significantly taller than their respective no-hormone controls (Figure 7; Table S18). Three inland annual populations (LMC, CAV, and MOR), the inland perennial population (SAL), and one near-coastal perennial population (JEN) were the most responsive to GA_3_ treatment (effect on height > 19 mm vs < 11 mm in the remaining populations; Figure 7).

**Figure 7.**
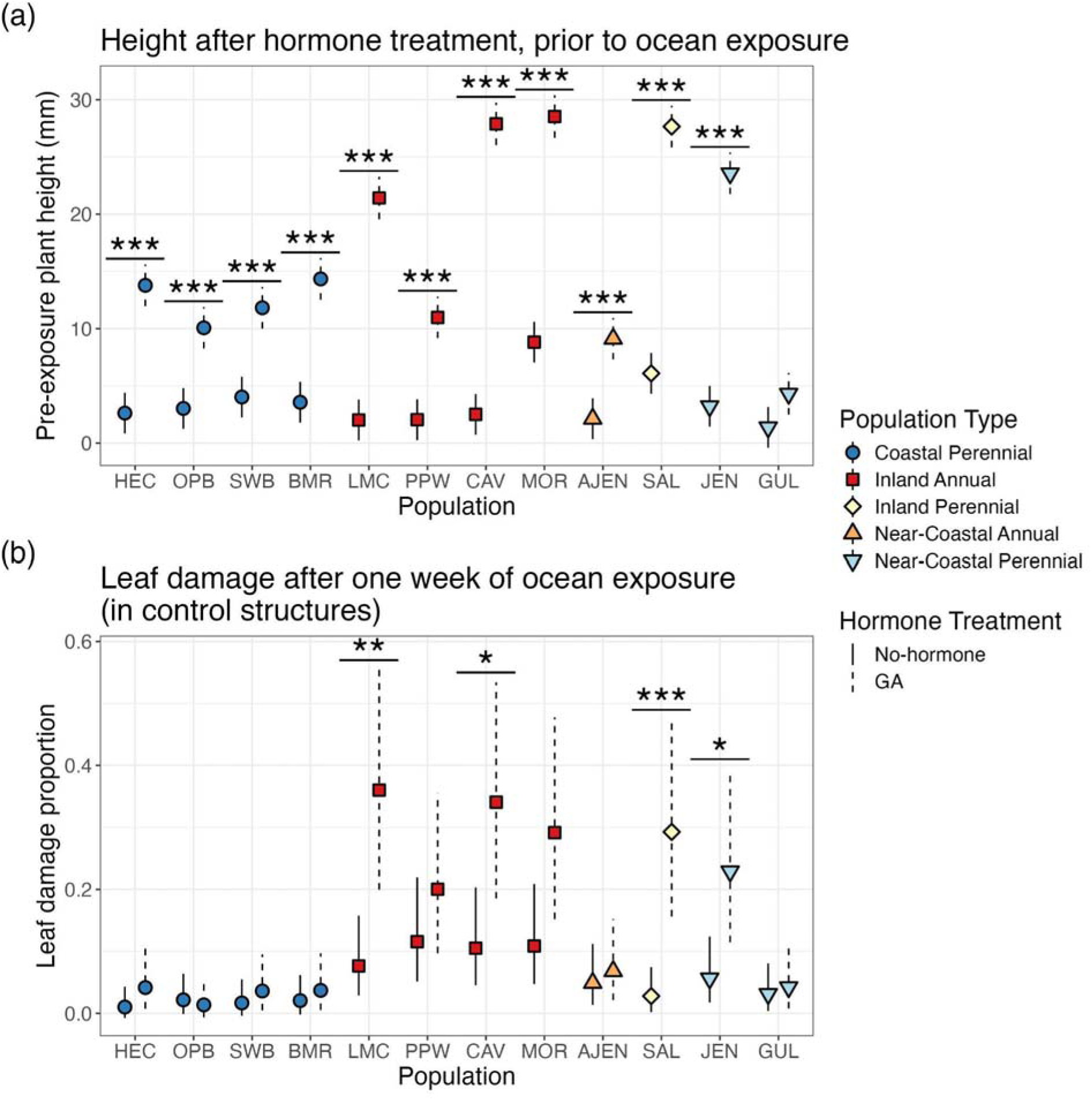
GA_3_ treatment increased plant height in a greenhouse, and increased leaf damage of plants exposed to salt spray in 2023. (a) Mean predictions and 95% confidence intervals from a linear mixed model, where the response variable was height in the greenhouse, the fixed predictor variables were population, GA_3_ treatment and their interaction, and flat was a random effect. (b) Mean predictions and 95% confidence intervals of a linear mixed model, where the response variable was logit transformed leaf damage in blocks exposed to the ocean, the fixed predictor variables were population, GA_3_ treatment, and their interaction, and flat nested within block was a random effect. Horizontal lines and asterisks (* *p* < 0.05, ** *p* < 0.01, *** *p* < 0.001) depict significant pairwise comparisons between GA_3_-treated plants and controls.

Agrofabric exclosures effectively protected seedlings from salt spray damage: 10% (21/216) of seedlings had tissue damage after a week in the exclosures compared to 78% (254/324) of seedlings in the open structures. In ocean-exposed open structures, leaf damage was significantly associated with population (Analysis of deviance: Wald Type III ^2^= 37.40, *df* =11, *p* < 0.001) and a population by GA_3_ treatment interaction (χ^2^= 28.41, *df* =11, *p* = 0.003), but was not significantly associated with hormone treatment (χ^2^= 2.34, *df* =1, *p* = 0.13). GA_3_ treatment significantly increased leaf damage by 224-952% only in four of the five accessions that increased most in height in response to GA_3_ (Table S19: p ≤ 0.03 , Figure 7, Figure S3).

## Discussion

Across environmental gradients, shifts in allocation between reproduction, growth, and defense follow predictable patterns, suggesting that these shifts underlie local adaptation (Bazzaz *et al*., 1987; Hahn & Maron, 2016; Züst & Agrawal, 2017). However, multiple abiotic and biotic factors co-vary across environmental gradients and multiple traits often differ between locally adapted populations, making the identification of selective agents and their phenotypic targets a major challenge (Wadgymar *et al*., 2017, 2022; López-Goldar & Agrawal, 2021). In this study, we used a manipulative reciprocal transplant experiment to test the hypothesis that herbivory and divergence in reproductive timing, vegetative growth, and defense against herbivores contributes to local adaptation across a coastal to inland environmental gradient.

At our coastal site, herbivore exclosures dramatically increased fecundity of local coastal perennials. However, contrary to our predictions, herbivory did not contribute to local adaptation, since excluding herbivores did not increase inland annual fitness at the coastal site. This is likely due to the abiotic effect of salt-spray pre-empting the impacts of herbivory on annuals. Our hormone treatments slightly shifted allocation between vegetative growth, reproduction and defense in each ecotype, but did not recapitulate the full effect size of differences previously observed in controlled greenhouse conditions (Lowry *et al*., 2019).

Nevertheless, we observed dramatic effects of our hormone treatments on survival and fecundity across our transplant sites. Despite delaying flowering, the GA_3_ application caused obvious earlier bolting and taller heights in the perennial transplants. A follow-up experiment confirmed that GA_3_ application both increased plant height and susceptibility to oceanic salt spray. Overall, our results suggest that oceanic salt spray is the dominant environmental factor driving local adaptation near the ocean and that coastal perennials significantly lose much of their locally adaptive salt spray tolerance advantage over inland annuals in response to GA_3_ application.

### Role of biotic interactions in local adaptation

The organisms a plant interacts with vary across the landscape, causing different selective pressures (Thompson, 2005; Urban, 2011; Friberg *et al*., 2019). Given the differences in the abiotic environment at our two sites–cool and foggy on the coast, hot and dry inland–the communities of organisms which our plants interact with differ substantially. The moist coastal environment has far more molluscan herbivores (snails, slugs), and a rare leaf-mining fly (Eiseman *et al*., 2023), which we did not see at the much drier inland site. Voles and deer also only contributed to herbivory at the coastal site. Given the differences in communities, differences in defense-levels, and prior research suggesting differences in intensity of herbivory on the coast and inland (Holeski, 2007), we predicted herbivory would be a driving factor in local adaptation. Surprisingly, we found no effects of herbivory on local adaptation at these two specific sites. At both sites, coastal perennials were more likely to be damaged by herbivores.

Although exclosures reduced deer herbivory at the coastal site, this reduction only benefited the local perennial ecotype since almost all annual transplants died of salt spray. While we did not find an effect of herbivory on local adaptation, we stress that insect-plant interactions regularly occur in a complex mosaic across the landscape and greatly vary temporally across years (Rotter *et al*., 2022).

#### High rate of herbivory at the coastal site did not contribute to local adaptation

At the coastal site, we predicted that high rates of herbivore attack would result in herbivory playing a strong role in local adaptation. A previous reciprocal transplant experiment found dramatic fitness differences between inland annuals grown in agrofabric exclosures that blocked all above ground factors, including herbivores and oceanic salt spray, and open structures at a coastal site (Popovic & Lowry, 2020). While we observed high rates of herbivory at the coastal field site, reducing deer herbivory with exclosures did not increase the success of inland annual plants because salt spray could get through the exclosures and had such a deleterious effect.

Annuals transplanted at the coastal site quickly exhibited necrosis from salt spray before dying; the window that they could have received herbivory was short, and they were likely poor quality host plants during that time. The perennials, in comparison, were larger, healthier, and had many more days in which to encounter an herbivore and receive damage. All reproductive herbivory at the coastal site (which had a very strong effect on fecundity) came after the median death date of our annual plants and thus was only experienced by perennials. These observations highlight the importance of the timing of selective events, particularly for local adaptation of ecotypes with differing life-history strategies. The importance of fecundity versus survival are likely to differ between ecotypes (DeMarche *et al*., 2016), and early-season factors (like coastal salt spray) that impact survival might disproportionately contribute to fitness differences between populations relative to late season factors that affect fecundity (such as herbivory; (Crone, 2001; Wadgymar *et al*., 2022).

While this study suggests that herbivory is preempted from playing a role in keeping annuals out of coastal environments, it does not mean it is unimportant. In the open structures on the coast, perennials completely failed to reproduce due to deer herbivory of inflorescences. By virtue of allocating growth to clonal expansion and non-reproductive tissue, perennials are likely increasing both tolerance (Stevens *et al*., 2008) and reducing the impacts of high-herbivory years on their lifetime fitness. Some populations are completely sterilized (i.e., all inflorescences are completely consumed by herbivores) in certain years (Toll, pers. obs.) and thus, herbivory may be an extremely strong selective pressure in the morphology, allocation to clonal growth, and reproductive timing of these coastal perennials. The results of this study were also clearly influenced by the close proximity of our coastal field site to the open ocean (within 50 meters of the shoreline). While it is common for coastal perennials to grow in close proximity to the ocean, where they are impacted by high levels of salt spray, it is also common for them to grow slightly further inland, where salt spray is greatly reduced (Boyce, 1954; Barbour, 1978; Du & Hesp, 2020). Coastal perennial populations of *M. guttatus* growing in close proximity to the ocean are generally shorter than coastal perennial populations that are further from the ocean, and presumably more protected from oceanic salt spray (Zambiasi & Lowry, 2024).

#### Life-history contributed to differences in herbivore attack at the inland site

Our finding that herbivory did not influence local adaptation inland is somewhat less surprising, as there is evidence that herbivore damage is generally less extensive there (Holeski, 2007). It was unexpected, however, that perennials were also more likely to be attacked by herbivores than annuals at the inland site, as we predicted that perennials would be more resistant to herbivory due to ecotype differences in phytochemical defenses (PPGs). Although early flowering can be a highly effective herbivory avoidance strategy (Krimmel & Pearse, 2016), perennials were more likely to be attacked even during periods of time when both ecotypes were alive in the same site suggesting the difference in attack rates was not due to a shorter lifespan for annuals. Higher attack rates for perennials could be due to differences in apparency caused by differences in plant size (Feeny, 1976), or herbivore preference due to nutritional differences or water content. In addition, while PPGs are feeding deterrents to generalists, some can be feeding stimulants for specialist herbivores (Holeski *et al*., 2013; Rotter *et al*., 2018), and thus perennials may be more likely to get attacked by specialists. Our presence-absence measure of herbivory also may have missed differences in degree of herbivory among plants that were attacked, which may have greater impacts on fitness.

### Oceanic salt spray sensitivity increased with gibberellin treatment

The most surprising result of our experiment was how dramatically GA_3_ application decreased survival of coastal perennial genotypes at the coastal field site in the 2020 reciprocal transplant experiment. Based on the patterns of plant tissue death, we attributed this mortality primarily to oceanic salt spray. Our 2023 experiment confirmed that GA_3_ addition made perennial and annual genotypes more susceptible to natural oceanic salt spray.

There are two non-mutually exclusive ways that GA_3_ could have decreased fitness in the coastal environment with regard to salt spray. First, the addition of GA_3_ increased plant height, as evident by increased internode elongation of plants here as well as in a prior study (Lowry *et al*., 2019). Increased plant height could put the aboveground portions of these plants more directly in the path of prevailing wind delivering the salt spray (Zambiasi & Lowry, 2024). In 2023, all seedlings treated with GA_3_ responded by increasing height, but populations that grew tallest in response to GA_3_ had the most tissue damage after a week of ocean exposure, consistent with increased height directly increasing plant exposure to salt spray.

A second hypothesis is that the GA_3_ treatment may directly increase the susceptibility of tissues to salt spray, independent of changes in plant height. The second hypothesis is particularly intriguing, as it is the opposite of what would be expected based on the soil salinity literature.

For example, previous experiments in rice (Rodríguez *et al*., 2006), wheat (Iqbal & Ashraf, 2013), apple (Wang *et al*., 2019), cucumber (Wang *et al*., 2020), and sorghum (Liu *et al*., 2023) have all found that the application of GA_3_ increases yields under saline conditions. The conflicting results of those studies and our experiment make it clear that findings from the soil salinity literature cannot be directly extrapolated to what is experienced by plants growing in coastal environments, where salt spray is a major source of stress on plant aboveground tissues (Boyce, 1954; Du & Hesp, 2020; Itoh *et al*., 2024). The exact mechanisms by which GA_3_ increases salt spray susceptibility are still not clear, but are an active focus of our current research.

While our study made the surprising discovery that GA_3_ addition makes plants more susceptible to salt spray, it is unclear the extent to which the adaptive changes in hormone pathways contributed to the evolution of the more salt spray-tolerant coastal perennial ecotype. Other recent studies have found evidence that evolutionary changes in both the gibberellin and jasmonic acid pathways contribute to the divergence between coastal perennial and inland annual ecotypes of *M. guttatus* (Olsen *et al*., 2024; Kollar *et al*., 2025). However, these evolutionary genetic changes in hormone pathways have yet to be directly linked to salt spray tolerance in this system or in other coastal ecotypes where hormone shifts have been implicated in their evolution (Wilkinson *et al*., 2021; James *et al*., 2023; Broad *et al*., 2024).

### Limitations and future directions

Although our study provided clear evidence that salt spray is a major abiotic factor contributing to local adaptation to coastal environments, our experiments had several limitations. First, we could not monitor transplants for the first few weeks of the COVID-19 lockdown. Future studies should monitor transplant fitness more frequently to capture early mortality. Second, we applied only one dose of each hormone and observed minimal trait responses. Applying multiple hormone doses in the field would allow researchers to test whether hormone effects on traits and fitness are dosage dependent. Third, mesh exclosures were chosen to reduce herbivory while also not excluding salt spray. The use of insecticides or molluscicides might further reduce different sources of herbivory, though those compounds can have unintended impacts on plants, such as through alterations of soil microbial communities. Our understanding of herbivory in this study was also limited by measuring damage as presence or absence. Attempts to quantify damage levels could be used to more precisely explore the role of herbivory in this system. Finally, we were unable to examine the role of herbivory in the absence of salt spray. Future studies should test the effect of herbivory on local adaptation in sites not directly exposed to salt spray.

## Conclusions

The evolution of local adaptation is inherently complex, involving multiple environmental factors that differ in importance among habitats and the evolution of multiple molecular and biochemical pathways. The results of our study illustrate the complexity of local adaptation and how studies to identify its underlying environmental, physiological, and genetic basis can lead to surprising yet important findings. The initial goal of our manipulative reciprocal transplant experiment was to investigate how hormone manipulation and herbivore exclosures affect the fitness of locally adaptive ecotypes in both local and foreign habitats. To our surprise, hormone manipulations ultimately had a minor impact on defensive compound production and herbivory. Instead, we accidentally discovered that the manipulation of gibberellin concentrations, through the addition of GA_3_, had a profound impact on the fitness of coastal plants in their home habitat. While it is well-established that interactions among hormone pathways mediate differences in allocation to growth, reproduction, and resistance in plants (Murphy, 2015; VanWallendael *et al*., 2019; Gasperini & Howe, 2024), few studies have investigated how evolutionary changes in hormone pathways contribute to the formation of plant ecotypes and local adaptation (Wilkinson *et al*., 2021; James *et al*., 2023; Broad *et al*., 2024), especially in the field. Future studies of local adaptation should aim to more explicitly understand how shifts in hormone pathways contribute to local adaptation and how those changes may have a broad and unexpected range of impacts on adaptations to various environmental factors. Critically, functional molecular studies should be conducted to test the hypothesis that adaptive genetic changes in hormone pathways underlie the evolution of locally adaptive shifts in resource allocation. While transgenic plants produced by those studies can readily be evaluated in the laboratory, government regulations discourage evaluation of those plants in natural habitats. The application of synthetic hormones and their antagonists, as conducted here, offer a much more acceptable way to test hypotheses about the roles of hormones in local adaptation in the field.

## Author contributions

Katherine Toll: Methodology, Investigation, Formal Analysis, Data Curation, Writing - Original Draft, Writing - Review & Editing, Visualization. Megan Blanchard: Methodology, Investigation, Formal Analysis, Data Curation, Writing - Original Draft, Writing - Review & Editing, Visualization. Anna Scharnagl: Investigation. Liza Holeski: Conceptualization, Methodology, Writing - Review & Editing, Resources, Supervision, Funding acquisition. David B. Lowry: Conceptualization, Methodology, Writing - Review & Editing, Resources, Supervision, Funding acquisition.

## Acknowledgements

We would like to thank Ben Blackman and Jack Collichio for germinating seeds at UC Berkeley. This work would not have been possible without the support of staff at the Bodega Marine Lab and Reserve, particularly Jackie Sones, Luis Morales, and Al Carranza, who facilitated field work and use of the greenhouse and lab. Michael Gillogly and Michelle Halbur of Pepperwood Preserve were instrumental to this research through their support of our work at the Preserve, including performing emergency plot-monitoring when the preserve temporarily closed to visitors due to the COVID-19 pandemic. Isabella Johnson and Billie Fraser assisted with chemical sample processing. This work was funded by National Science Foundation Division of Integrative Organismal Systems Grants to DBL and LMH (IOS-1855927) and DBL (IOS-2153100).

