## Supplementary materials for "Plant hormone manipulation impacts salt spray tolerance, which preempts herbivory as a driver of local adaptation in the yellow monkeyflower, *Mimulus guttatus*"

### Supplementary material

### Materials and Methods

#### Statistical analyses

Analysis of herbivory by putative source

At the coastal transplant site, we were able to visually distinguish between damage from leaf miners, caterpillars, slugs, deer, beetles, and voles. When we subdivided herbivory by putative source, only one annual category (treated with paclobutrazol in open structures damaged by leaf miners) had more than one observation. Therefore, we only analyzed herbivory by source in coastal perennials. To test whether hormone treatments affected damage from particular types of herbivores and whether exclosures reduced herbivory for specific herbivore types in coastal perennials, we fit mixed models for each response variable that included hormone treatment and exclosure type as interactive fixed factors. The response variables were the presence or absence of herbivory from leaf miners, caterpillars, slugs, beetles, deer, and voles. The models included transplant block and maternal family as random effects.

For deer damage, there was no variance in one of the treatment levels (quasi-complete separation) and a binomial model was not able to accurately estimate the parameter for that treatment group. For this reason, we used a randomization test to estimate the probability that this result was due to chance, by reassigning treatments to all outcomes 10,000 times and calculating the proportion of these reshuffles that resulted in a difference as extreme as the observed (Gotelli and Ellison 2004).

### Results

##### Annuals flowered earlier, investing less in vegetative growth than perennials, and hormone treatments slightly delayed flowering time

At the coastal site, gibberellic acid ($GA_3$) and paclobutrazol slightly, but significantly, delayed annual flowering time relative to no-hormone control annuals (plants sprayed with 0.25% ethanol). Paclobutrazol-treated and $GA_3$-treated annuals flowered 10-17 days later than controls (Fig. 2A, Tukey post-hoc tests: Table S3). However, these effects were small relative to ecotype differences in flowering time: all annuals flowered 46 to 64 days earlier than their corresponding hormone treated perennials, all significant differences (Table S4).

At the inland site, in the control structures only, $GA_3$ slightly, but significantly, delayed annual flowering time relative to the control. $GA_3$-treated annuals in open structures flowered 9 days later than no-hormone control annuals in open structures (Fig. 2B, Tukey post-hoc tests: Table S4). Again, this effect was small relative to ecotype differences in flowering time: all annuals flowered 39 to 62 days earlier than their perennials with the same hormone treatment in both the exclosures and control structures, all significant differences (Table S5).

Hormone treatments had no effect on perennial flowering time relative to controls at either transplant site (Tables S4 & S5). Exclosures had no effect on flowering time at either transplant site (Tables S1, S2, S4, S5).

#### Exclosures reduced deer herbivory, $GA_3$ reduced leaf miner, caterpillar, and deer herbivory at the coastal site

All results for herbivore damage by source are for coastal perennials at the coastal transplant site.

For leaf miner damage, the minimum adequate model included hormone treatment, exclosure treatment and their interaction. Leaf miner damage in coastal perennials was significantly associated with hormone treatment (Analysis of deviance: Wald Type III $\chi^2 = 53.2897$, $df = 3$, $p < 0.001$) and a hormone treatment by exclosure treatment interaction ($\chi^2 = 13.5574$, $df = 3$, $p = 0.004$), but was not significantly associated with exclosure treatment ($\chi^2$=, *df* = , *p* = ). $GA_3$ reduced the probability of leaf miner herbivory by 96% and 84% ($GA_3$ vs no-hormone contrasts in open structures, $p < 0.001$, and exclosures, $p = 0.05$). Leaf miner herbivory did not significantly differ between open structures and exclosures (open structures vs controls within each hormone treatment, $p > 0.05$).

For caterpillar damage, the minimum adequate model included hormone treatment and exclosure treatment. An interactive model with a hormone by exclosure treatment interaction did not significantly improve the model fit compared to the additive model (LRT: $\chi^2 = 0.1391$, $df = 3$, $p = 0.9868$). Caterpillar damage was significantly associated with hormone treatment (Analysis of deviance: Wald Type II $\chi^2 = 20.7520$, $df = 3$, $p < 0.001$) and exclosure treatment ($\chi^2 = 4.6693$, $df = 1$, $p = 0.031$). $GA_3$ reduced the probability of caterpillar herbivory by 79-83% ($GA_3$ vs no-hormone contrasts in open structures and exclosures, $p = 0.016$). Caterpillar herbivory did not significantly differ between open structures and exclosures (open structures vs controls within each hormone treatment, $p > 0.05$).

For slug damage, an interactive model with a hormone by exclosure treatment interaction did not significantly improve the model fit compared to the additive model (LRT: $\chi^2 = 0.7074$, $df = 3$, $p = 0.8715$). A model with hormone treatment and exclosure treatment did not improve the model fit compared to a model with hormone treatment (LRT: $\chi^2 = 0.8153$, $df = 1$, $p = 0.3666$). Finally, a model with hormone treatment did not improve the model fit compared to a model with only random effects (LRT: $\chi^2 = 5.8185$, $df = 3$, $p = 0.1208$).

For deer damage, the minimum adequate model included hormone treatment and exclosure treatment. An interactive model with a hormone by exclosure treatment interaction did not significantly improve the model fit compared to the additive model (LRT: $\chi^2 = 0$, $df = 3$, $p = 1$). Deer damage was significantly associated with hormone treatment (Analysis of deviance: Wald Type II $\chi^2 = 29.369$, $df = 3$, $p < 0.001$), but was not significantly associated with exclosure treatment ($\chi^2 = 0$, $df = 1$, $p = 0.9974$). $GA_3$ reduced the probability of deer herbivory by 85-86 % (Figure S1; GA vs no-hormone contrasts in open structures and exclosures, $p \leq 0.001$).

Since no perennials were damaged by deer in the exclosures (Figure S1), the binomial model could not accurately estimate deer herbivory for perennials in exclosures. Therefore, we used a randomization test to estimate the probability that the difference in deer herbivory between open structures and exclosures was due to chance. Randomization tests showed that 0 (no-hormone, MeJA, and paclobutrazol treatment) and 19 ($GA_3$ treatment) of 10,000 treatment reshuffles resulted in the observed difference in deer herbivory between open structures and exclosures, making it unlikely that this pattern arose by chance ($p = 0$; $p = 0.0019$).

For beetle damage, an interactive model with a hormone by exclosure treatment interaction did not significantly improve the model fit compared to the additive model (LRT: $\chi^2 = 6.7743$, $df = 3$, $p = 0.07945$). A model with hormone treatment and exclosure treatment did not improve the model fit compared to a model with exclosure treatment (LRT: $\chi^2 = 0.8137$, $df = 3$, $p = 0.8462$). Finally, a model with exclosure treatment did not improve the model fit compared to a model with only random effects (LRT: $\chi^2 = 1.8802$, $df = 1$, $p = 0.1703$).

For vole damage, an interactive model with a hormone by exclosure treatment interaction did not significantly improve the model fit compared to the additive model (LRT: $\chi^2 = 0$, $df = 3$, $p = 1$). A model with hormone treatment and exclosure treatment did not improve the model fit compared to a model with hormone treatment (LRT: $\chi^2 = 3.2342$, $df = 1$, $p = 0.07211$). Finally, a model with hormone treatment did not improve the model fit compared to a model with only random effects (LRT: $\chi^2 = 1.4998$, $df = 3$, $p = 0.6823$).

#### Perennials produced more PPGs, methyl jasmonate and $GA_3$ increased PPGs at the inland site, while GA decreased PPGs at the coastal site

At the inland site, perennials had significantly higher total PPG concentration than annuals (Tukey post-hoc tests: Table S9) and annuals and perennials differed in their multivariate PPG arsenals. The effect of ecotype was generally stronger than any hormone effects. We were unable to compare annuals and perennials at the coastal site due to high annual mortality.

At both sites, exclosures had no effect on total PPG concentration (Table S1), though exclosures did moderate the effect of hormone treatment at the coastal site (Table S2). Exclosure did not influence the multivariate PPG arsenal at the coastal site but did at the inland site (Table S3). At the coast, the only effect of hormone treatment was that $GA_3$ reduced total PPG concentration of perennials in the control plots (Figure 3a, Tukey post-hoc tests, Table S8) and caused the PPG arsenal to differ from control plants (Figure 3c, PERMANOVA pairwise, Table S10). While this impact of $GA_3$ is consistent with our predictions that $GA_3$ downregulates defense-allocation, it is also possible that the decrease in total PPG is due to increased salt-stress experienced by $GA_3$-treated plants. Inland, hormone treatments did not influence PPGs in perennials (Figure 3b,c, Table S9). In annuals at the inland site, $GA_3$ and MeJA increased the total concentration of PPGs (Figure 3b, Table S10) and caused the PPG arsenal to differ (Figure 3d, PERMANOVA pairwise, Table S11). While we expected MeJA to increase allocation to defense, we expected $GA_3$ to decrease it. However, the increase in total PPG is consistent with an increase in days to flowering in $GA_3$-treated annuals at the inland site (these traits positively covary in annuals, Kooyers et al. 2020), though the mechanism for this shift is unknown.

**Table S1.** Univariate analysis: Likelihood ratio tests comparing models with and without individual factors. Each factor dropped between models was not significantly associated with the response variable in the complex model (via analysis of deviance, not reported), and did not significantly improve model fit ($p > 0.05$ in each likelihood ratio test) and were sequentially removed to identify minimum adequate models. The full model for each response variable (except for total PPG and fruit number at the coastal transplant site) included ecotype, exclosure type, hormone treatment, all two-way interactions, and the three-way interaction as fixed effects, and block and maternal family as random effects. All sequentially reduced models included block and maternal family as random effects. At the coastal transplant site, we were only able to measure and analyze PPGs for perennials due to high annual mortality. Also at the coastal site, no plants produced fruit outside of the exclosures, and few annuals produced fruit inside the exclosures, so we only analyzed the effect of hormone treatments on perennial fruit production inside the exclosures.

| Response Variable | Site | Complex model | Simpler model | Factor dropped between models | χ2 | *df* | *p* |
|---|---|---|---|---|---|---|---|
| **Flowering Time** | Coast | Ecotype + Hormone + Exclosure + Ecotype x Hormone + Ecotype x Exclosure + Hormone x Exclosure + Ecotype x Hormone x Exclosure | Ecotype + Hormone + Exclosure + Ecotype x Hormone + Ecotype x Exclosure + Hormone x Exclosure | Ecotype x Hormone x Exclosure | 3.86 | 3 | 0.28 |
| | | Ecotype + Hormone + Exclosure + Ecotype x Hormone + Ecotype x Exclosure + Hormone x Exclosure | Ecotype + Hormone + Exclosure + Ecotype x Hormone + Ecotype x Exclosure | Hormone x Exclosure | 5.13 | 3 | 0.16 |
| | | Ecotype + Hormone + Exclosure + Ecotype x Hormone + Ecotype x Exclosure | Ecotype + Hormone + Exclosure + Ecotype x Hormone | Ecotype x Exclosure | 2.07 | 1 | 0.15 |
| | | Ecotype + Hormone + Exclosure + Ecotype x Hormone | Ecotype + Hormone + Ecotype x Hormone | Exclosure | 0.19 | 1 | 0.66 |
| **Herbivory Probability** | Coast | Ecotype + Hormone + Exclosure + Ecotype x Hormone + Ecotype x | Ecotype + Hormone + Exclosure + Ecotype x | Ecotype x Hormone x Exclosure | 4.01 | 3 | 0.26 |

| | | | | | | | |
|---|---|---|---|---|---|---|---|
| | | Exclosure + Hormone x Exclosure + Ecotype x Hormone x Exclosure | Hormone + Ecotype x Exclosure + Hormone x Exclosure | | | | |
| | | Ecotype + Hormone + Exclosure + Ecotype x Hormone + Ecotype x Exclosure + Hormone x Exclosure | Ecotype + Hormone + Exclosure + Ecotype x Exclosure + Hormone x Exclosure | Ecotype x Hormone | 0.48 | 3 | 0.92 |
| **Total PPG** | Inland | Ecotype + Hormone + Exclosure + Ecotype x Hormone + Ecotype x Exclosure + Hormone x Exclosure + Ecotype x Hormone x Exclosure | Ecotype + Hormone + Exclosure + Ecotype x Hormone + Ecotype x Exclosure + Hormone x Exclosure | Ecotype x Hormone x Exclosure | 0.27 | 3 | 0.97 |
| | | Ecotype + Hormone + Exclosure + Ecotype x Hormone + Ecotype x Exclosure + Hormone x Exclosure | Ecotype + Hormone + Exclosure + Ecotype x Hormone + Hormone x Exclosure | Ecotype x Exclosure | 0.01 | 3 | 0.94 |
| | | Ecotype + Hormone + Exclosure + Ecotype x Hormone + Hormone x Exclosuree | Ecotype + Hormone + Exclosure + Ecotype x Hormone | Hormone x Exclosure | 0.43 | 3 | 0.93 |
| | | Ecotype + Hormone + Exclosure + Ecotype x Hormone | Ecotype + Hormone + Ecotype x Hormone | Exclosure | 0.18 | 1 | 0.67 |
| **Survival Probability** | Inland | Ecotype + Hormone + Exclosure + Ecotype x Hormone + Ecotype x Exclosure + Hormone x Exclosure + Ecotype | Ecotype + Hormone + Exclosure + Ecotype x Hormone + Ecotype x Exclosure + | Ecotype x Hormone x Exclosure | 2.98 | 3 | 0.39 |

| | | | | | | | |
|---|---|---|---|---|---|---|---|
| | | x Hormone x Exclosure | Hormone x Exclosure | | | | |
| | | Ecotype + Hormone + Exclosure + Ecotype x Hormone + Ecotype x Exclosure + Hormone x Exclosure | Ecotype + Hormone + Exclosure + Ecotype x Exclosure + Hormone x Exclosure | Ecotype x Hormone | 0.10 | 3 | 0.99 |
| | | Ecotype + Hormone + Exclosure + Ecotype x Exclosure + Hormone x Exclosure | Ecotype + Hormone + Exclosure + Hormone x Exclosure | Ecotype x Exclosure | 0.94 | 1 | 0.33 |
| | | Ecotype + Hormone + Exclosure + Hormone x Exclosure | Ecotype + Hormone + Exclosure | Hormone x Exclosure | 1.19 | 3 | 0.76 |
| | | Ecotype + Hormone + Exclosure | Ecotype + Exclosure | Hormone Treatment | 0.26 | 3 | 0.97 |
| | | Ecotype + Exclosure | Ecotype | Exclosure Treatment | 1.06 | 1 | 0.30 |
| **Flowering Probability** | Coast | Ecotype + Hormone + Exclosure + Ecotype x Hormone + Ecotype x Exclosure + Hormone x Exclosure + Ecotype x Hormone x Exclosure | Ecotype + Hormone + Exclosure + Ecotype x Hormone + Ecotype x Exclosure + Hormone x Exclosure | Ecotype x Hormone x Exclosure | 3.13 | 3 | 0.37 |
| | | Ecotype + Hormone + Exclosure + Ecotype x Hormone + Ecotype x Exclosure + Hormone x Exclosure | Ecotype + Hormone + Exclosure + Ecotype x Hormone + Ecotype x Exclosure | Hormone x Exclosure | 3.78 | 3 | 0.29 |
| | | Ecotype + Hormone + Exclosure + Ecotype x Hormone + Ecotype x Exclosure | Ecotype + Hormone + Exclosure + Ecotype x Exclosure | Ecotype x Hormone | 6.90 | 3 | 0.08 |

| | | | | | | | |
|---|---|---|---|---|---|---|---|
| | Inland | Ecotype + Hormone + Exclosure + Ecotype x Hormone + Ecotype x Exclosure + Hormone x Exclosure + Ecotype x Hormone x Exclosure | Ecotype + Hormone + Exclosure + Ecotype x Hormone + Ecotype x Exclosure + Hormone x Exclosure | Ecotype x Hormone x Exclosure | 0.00 | 3 | 1.00 |
| | | Ecotype + Hormone + Exclosure + Ecotype x Hormone + Ecotype x Exclosure + Hormone x Exclosure | Ecotype + Hormone + Exclosure + Ecotype x Hormone + Ecotype x Exclosure | Hormone x Exclosure | 1.21 | 3 | 0.75 |
| **Fruit number** | Coast | Hormone | no fixed effects (only random effects) | Hormone Treatment | 2.48 | 3 | 0.48 |
| | Inland | Ecotype + Hormone + Exclosure + Ecotype x Hormone + Ecotype x Exclosure + Hormone x Exclosure + Ecotype x Hormone x Exclosure | Ecotype + Hormone + Exclosure + Ecotype x Hormone + Ecotype x Exclosure + Hormone x Exclosure | Ecotype x Hormone x Exclosure | 3.19 | 3 | 0.36 |
| | | Ecotype + Hormone + Exclosure + Ecotype x Hormone + Ecotype x Exclosure + Hormone x Exclosure | Ecotype + Hormone + Exclosure + Ecotype x Hormone + Ecotype x Exclosure | Hormone x Exclosure | 4.13 | 3 | 0.25 |
| | | Ecotype + Hormone + Exclosure + Ecotype x Hormone + Ecotype x Exclosure | Ecotype + Hormone + Exclosure + Ecotype x Exclosure | Ecotype x Hormone | 4.69 | 3 | 0.20 |
| | | Ecotype + Hormone + Exclosure + Ecotype x Exclosure | Ecotype + Hormone + Exclosure | Ecotype x Exclosure | 2.74 | 1 | 0.10 |

**Table S2.** Univariate analysis: Analysis of deviance (Wald Type III tests for models with interactions and Wald Type II tests for fully additive models) table for factors in minimum adequate models for each response variable.

| Response Variable | Site | Factor | χ2 | *df* | *p*-value |
|---|---|---|---|---|---|
| **Flowering Time** | Coast | Ecotype | 272.08 | 1 | **< 0.001** |
| | | Hormone Treatment | 15.7 | 3 | **0.001** |
| | | Ecotype x Hormone | 9.36 | 3 | **0.025** |
| | Inland | Ecotype | 294.07 | 1 | **< 0.001** |
| | | Hormone Treatment | 19.21 | 3 | **< 0.001** |
| | | Exclosure Treatment | 0.04 | 1 | 0.84 |
| | | Ecotype x Hormone | 11.53 | 3 | **0.009** |
| | | Ecotype x Exclosure | 0.92 | 1 | 0.34 |
| | | Hormone x Exclosure | 7.50 | 3 | 0.06 |
| | | Ecotype x Hormone x Exclosure | 11.40 | 3 | **0.01** |
| **Herbivory Probability** | Coast | Ecotype | 152.95 | 1 | **< 0.001** |
| | | Hormone Treatment | 2.87 | 1 | 0.09 |
| | | Exclosure Treatment | 85.36 | 3 | **< 0.001** |
| | | Ecotype x Exclosure | 34.52 | 1 | **< 0.001** |
| | | Hormone x Exclosure | 20.79 | 3 | **< 0.001** |
| | Inland | Ecotype | 28.71 | 1 | **< 0.001** |
| | | Hormone Treatment | 4.33 | 3 | 0.23 |
| | | Exclosure Treatment | 0.87 | 1 | 0.35 |
| | | Ecotype x Hormone | 8.95 | 3 | **0.030** |
| | | Ecotype x Exclosure | 4.98 | 1 | **0.026** |
| | | Hormone x Exclosure | 2.09 | 3 | 0.56 |
| | | Ecotype x Hormone x Exclosure | 1.59 | 3 | 0.66 |
| **Total PPG** | Coast | Hormone Treatment | 28.11 | 3 | **<0.001** |
| | | Exclosure Treatment | 0.95 | 1 | 0.330 |
| | | Hormone x Exclosure | 11.78 | 3 | **0.008** |
| | Inland | Ecotype | 229.42 | 1 | **< 0.001** |
| | | Hormone Treatment | 15.00 | 3 | **0.002** |
| | | Ecotype x Hormone | 11.28 | 3 | **0.010** |
| **Survival Probability** | Coast | Ecotype | 86.75 | 1 | **< 0.001** |
| | | Hormone Treatment | 82.75 | 3 | **< 0.001** |
| | | Exclosure Treatment | 4.05 | 1 | **0.04** |
| | | Ecotype x Hormone | 32.42 | 3 | **< 0.001** |
| | | Ecotype x Exclosure | 1.26 | 1 | 0.26 |
| | | Hormone x Exclosure | 1.67 | 3 | 0.64 |

| | | | | | |
|---|---|---|---|---|---|
| | | Ecotype x Hormone x Exclosure | 12.39 | 3 | **< 0.006** |
| | Inland | Ecotype | 499.27 | 1 | **< 0.001** |
| **Flowering Probability** | Coast | Ecotype | 0.12 | 1 | 0.73 |
| | | Hormone Treatment | 99.69 | 3 | **< 0.001** |
| | | Exclosure Treatment | 7.28 | 1 | **0.007** |
| | | Ecotype x Exclosure | 6.58 | 1 | **< 0.001** |
| | Inland | Ecotype | 22.93 | 1 | **< 0.001** |
| | | Hormone Treatment | 2.03 | 3 | 0.57 |
| | | Exclosure Treatment | 0.00 | 1 | 1.00 |
| | | Ecotype x Hormone | 11.03 | 3 | **0.01** |
| | | Ecotype x Exclosure | 6.90 | 3 | 0.08 |
| **Fruit Number** | Inland | Ecotype | 0.10 | 1 | 0.75 |
| | | Exclosure Treatment | 3.59 | 1 | 0.06 |
| | | Hormone Treatment | 34.467 | 3 | **< 0.001** |

**Table S3.** Multivariate analysis: Permutational multivariate analysis of variance (PERMANOVA) table for factors in simplified models.

| **Response Variable** | **Site** | **Factor** | ***df*** | **SS** | $R^2$ | **Pseudo *F*** | ***p* (Perm)** |
|---|---|---|---|---|---|---|---|
| **PPG Arsenal** | Coast | Hormone Treatment | 3 | 0.134 | 0.024 | 1.770 | **0.02** |
| | Inland | Ecotype | 1 | 16.194 | 0.470 | 555.673 | **0.001** |
| | | Hormone Treatment | 3 | 0.365 | 0.011 | 4.171 | **0.003** |
| | | Exclosure Treatment | 1 | 0.383 | 0.011 | 13.139 | **0.001** |
| | | Ecotype x Hormone | 3 | 0.319 | 0.009 | 3.650 | **0.002** |

**Table S4**. Tukey post-hoc contrasts for days to flowering at the coastal site, Bodega Marine Reserve. Minimum adequate model: Days to flowering = Ecotype + Hormone Treatment + Ecotype x Hormone treatment + (1|maternal family) + (1|Plot). *P*-values < 0.05 in bold. Contrasts are structured: Ecotype, Hormone treatment vs Ecotype, Hormone treatment. For hormone treatment: control = no-hormone, GA = $GA_3$ treatment, MeJA = methyl jasmonate treatment, and Paclo = paclobutrazol.

| **contrast** | **estimate** | **SE** | ***df*** | ***z*-ratio** | ***p*-value** |
|---|---|---|---|---|---|
| Annual Control - Perennial Control | -63.422 | 3.845 | 414 | -16.495 | **< 0.001** |
| Annual Control - Annual GA | -16.045 | 5.180 | 414 | -3.098 | **0.043** |
| Annual Control - Perennial GA | -62.163 | 4.526 | 414 | -13.734 | **< 0.001** |
| Annual Control - Annual Paclo | -11.271 | 3.465 | 414 | -3.253 | **0.027** |
| Annual Control - Perennial Paclo | -72.823 | 3.874 | 414 | -18.797 | **< 0.001** |

| | | | | | |
|---|---|---|---|---|---|
| Annual Control - Annual MeJA | -6.356 | 3.115 | 414 | -2.041 | 0.456 |
| Annual Control - Perennial MeJA | -61.265 | 3.850 | 414 | -15.914 | **< 0.001** |
| Perennial Control - Annual GA | 47.377 | 5.591 | 414 | 8.474 | **< 0.001** |
| Perennial Control - Perennial GA | 1.259 | 3.938 | 414 | 0.320 | 1.000 |
| Perennial Control - Annual Paclo | 52.152 | 4.045 | 414 | 12.894 | **< 0.001** |
| Perennial Control - Perennial Paclo | -9.401 | 3.145 | 414 | -2.989 | 0.059 |
| Perennial Control - Annual MeJA | 57.066 | 3.746 | 414 | 15.236 | **< 0.001** |
| Perennial Control - Perennial MeJA | 2.157 | 3.112 | 414 | 0.693 | 0.997 |
| Annual GA - Perennial GA | -46.118 | 6.043 | 414 | -7.632 | **< 0.001** |
| Annual GA - Annual Paclo | 4.774 | 5.344 | 414 | 0.893 | 0.987 |
| Annual GA - Perennial Paclo | -56.778 | 5.612 | 414 | -10.117 | **< 0.001** |
| Annual GA - Annual MeJA | 9.689 | 5.135 | 414 | 1.887 | 0.561 |
| Annual GA - Perennial MeJA | -45.220 | 5.594 | 414 | -8.084 | **< 0.001** |
| Perennial GA - Annual Paclo | 50.892 | 4.665 | 414 | 10.909 | **< 0.001** |
| Perennial GA - Perennial Paclo | -10.660 | 3.952 | 414 | -2.697 | 0.126 |
| Perennial GA - Annual MeJA | 55.807 | 4.455 | 414 | 12.528 | **< 0.001** |
| Perennial GA - Perennial MeJA | 0.898 | 3.933 | 414 | 0.228 | 1.000 |
| Annual Paclo - Perennial Paclo | -61.552 | 4.077 | 414 | -15.098 | **< 0.001** |
| Annual Paclo - Annual MeJA | 4.915 | 3.390 | 414 | 1.450 | 0.833 |
| Annual Paclo - Perennial MeJA | -49.995 | 4.052 | 414 | -12.340 | **< 0.001** |
| Perennial Paclo - Annual MeJA | 66.467 | 3.782 | 414 | 17.576 | **< 0.001** |
| Perennial Paclo - Perennial MeJA | 11.558 | 3.135 | 414 | 3.686 | **0.006** |
| Annual MeJA - Perennial MeJA | -54.909 | 3.758 | 414 | -14.612 | **< 0.001** |

**Table S5**. Tukey post-hoc contrasts for days to flowering at the inland site, Pepperwood Preserve. Minimum adequate model: Days to flowering = Ecotype + Hormone Treatment + Exclosure + Ecotype x Hormone + Ecotype x Exclosure + Hormone x Exclosure + (1|maternal family) + (1|Plot). *P*-values < 0.05 in bold. Contrasts are structured: Ecotype, Hormone treatment, Exclosure treatment vs Ecotype, Hormone treatment, Exclosure treatment. For hormone treatment: control = no-hormone, GA = $GA_3$ treatment, MeJA = methyl jasmonate treatment, and Paclo = paclobutrazol. For exclosure treatment: control = open structures, exclosures = mesh exclosures.

| **contrast** | **estimate** | **SE** | ***df*** | ***z*-ratio** | ***p*-value** |
|---|---|---|---|---|---|
| Annual Control Control - Perennial Control Control | -51.665 | 3.014 | 704 | -17.139 | **< 0.001** |
| Annual Control Control - Annual GA Control | -9.167 | 2.327 | 704 | -3.939 | **0.009** |
| Annual Control Control - Perennial GA Control | -47.964 | 3.869 | 704 | -12.397 | **< 0.001** |
| Annual Control Control - Annual Paclo Control | -3.538 | 2.296 | 704 | -1.541 | 0.977 |
| Annual Control Control - Perennial Paclo Control | -54.636 | 3.462 | 704 | -15.780 | **< 0.001** |
| Annual Control Control - Annual MeJA Control | -0.779 | 2.345 | 704 | -0.332 | 1.000 |

| | | | | | |
|---|---|---|---|---|---|
| Annual Control Control - Perennial MeJA Control | -54.233 | 3.188 | 704 | -17.009 | **< 0.001** |
| Annual Control Control - Annual Control Exclosure | 0.734 | 3.570 | 704 | 0.206 | 1.000 |
| Annual Control Control - Perennial Control Exclosure | -54.715 | 3.924 | 704 | -13.944 | **< 0.001** |
| Annual Control Control - Annual GA Exclosure | -1.459 | 3.549 | 704 | -0.411 | 1.000 |
| Annual Control Control - Perennial GA Exclosure | -60.144 | 4.205 | 704 | -14.304 | **< 0.001** |
| Annual Control Control - Annual Paclo Exclosure | -1.950 | 3.570 | 704 | -0.546 | 1.000 |
| Annual Control Control - Perennial Paclo Exclosure | -63.922 | 4.224 | 704 | -15.134 | **< 0.001** |
| Annual Control Control - Annual MeJA Exclosure | -2.398 | 3.559 | 704 | -0.674 | 1.000 |
| Annual Control Control - Perennial MeJA Exclosure | -55.677 | 3.952 | 704 | -14.087 | **< 0.001** |
| Perennial Control Control - Annual GA Control | 42.498 | 3.006 | 704 | 14.138 | **< 0.001** |
| Perennial Control Control - Perennial GA Control | 3.700 | 3.874 | 704 | 0.955 | 1.000 |
| Perennial Control Control - Annual Paclo Control | 48.126 | 2.982 | 704 | 16.137 | **< 0.001** |
| Perennial Control Control - Perennial Paclo Control | -2.971 | 3.473 | 704 | -0.855 | 1.000 |
| Perennial Control Control - Annual MeJA Control | 50.886 | 3.021 | 704 | 16.841 | **< 0.001** |
| Perennial Control Control - Perennial MeJA Control | -2.568 | 3.184 | 704 | -0.807 | 1.000 |
| Perennial Control Control - Annual Control Exclosure | 52.399 | 4.047 | 704 | 12.948 | **< 0.001** |
| Perennial Control Control - Perennial Control Exclosure | -3.050 | 3.929 | 704 | -0.776 | 1.000 |
| Perennial Control Control - Annual GA Exclosure | 50.206 | 4.029 | 704 | 12.461 | **< 0.001** |
| Perennial Control Control - Perennial GA Exclosure | -8.479 | 4.201 | 704 | -2.018 | 0.814 |
| Perennial Control Control - Annual Paclo Exclosure | 49.715 | 4.046 | 704 | 12.287 | **< 0.001** |
| Perennial Control Control - Perennial Paclo Exclosure | -12.257 | 4.239 | 704 | -2.891 | 0.222 |
| Perennial Control Control - Annual MeJA Exclosure | 49.267 | 4.037 | 704 | 12.203 | **< 0.001** |
| Perennial Control Control - Perennial MeJA Exclosure | -4.012 | 3.967 | 704 | -1.011 | 1.000 |
| Annual GA Control - Perennial GA Control | -38.798 | 3.856 | 704 | -10.061 | **< 0.001** |
| Annual GA Control - Annual Paclo Control | 5.628 | 2.279 | 704 | 2.470 | 0.495 |
| Annual GA Control - Perennial Paclo Control | -45.469 | 3.451 | 704 | -13.174 | **< 0.001** |
| Annual GA Control - Annual MeJA Control | 8.388 | 2.328 | 704 | 3.604 | 0.029 |
| Annual GA Control - Perennial MeJA Control | -45.066 | 3.178 | 704 | -14.179 | **< 0.001** |
| Annual GA Control - Annual Control Exclosure | 9.901 | 3.558 | 704 | 2.783 | 0.281 |
| Annual GA Control - Perennial Control Exclosure | -45.548 | 3.914 | 704 | -11.636 | **< 0.001** |
| Annual GA Control - Annual GA Exclosure | 7.708 | 3.537 | 704 | 2.179 | 0.712 |
| Annual GA Control - Perennial GA Exclosure | -50.977 | 4.196 | 704 | -12.149 | **< 0.001** |
| Annual GA Control - Annual Paclo Exclosure | 7.217 | 3.558 | 704 | 2.028 | 0.809 |
| Annual GA Control - Perennial Paclo Exclosure | -54.755 | 4.215 | 704 | -12.991 | **< 0.001** |
| Annual GA Control - Annual MeJA Exclosure | 6.769 | 3.547 | 704 | 1.908 | 0.872 |
| Annual GA Control - Perennial MeJA Exclosure | -46.510 | 3.943 | 704 | -11.797 | **< 0.001** |
| Perennial GA Control - Annual Paclo Control | 44.426 | 3.842 | 704 | 11.564 | **< 0.001** |
| Perennial GA Control - Perennial Paclo Control | -6.671 | 4.209 | 704 | -1.585 | 0.971 |

| | | | | | |
|---|---|---|---|---|---|
| Perennial GA Control - Annual MeJA Control | 47.186 | 3.870 | 704 | 12.193 | **< 0.001** |
| Perennial GA Control - Perennial MeJA Control | -6.268 | 4.004 | 704 | -1.566 | 0.974 |
| Perennial GA Control - Annual Control Exclosure | 48.699 | 4.712 | 704 | 10.334 | **< 0.001** |
| Perennial GA Control - Perennial Control Exclosure | -6.750 | 4.612 | 704 | -1.464 | 0.986 |
| Perennial GA Control - Annual GA Exclosure | 46.506 | 4.697 | 704 | 9.902 | **< 0.001** |
| Perennial GA Control - Perennial GA Exclosure | -12.180 | 4.846 | 704 | -2.513 | 0.463 |
| Perennial GA Control - Annual Paclo Exclosure | 46.015 | 4.712 | 704 | 9.766 | **< 0.001** |
| Perennial GA Control - Perennial Paclo Exclosure | -15.957 | 4.872 | 704 | -3.275 | 0.082 |
| Perennial GA Control - Annual MeJA Exclosure | 45.567 | 4.704 | 704 | 9.687 | **< 0.001** |
| Perennial GA Control - Perennial MeJA Exclosure | -7.712 | 4.635 | 704 | -1.664 | 0.955 |
| Annual Paclo Control - Perennial Paclo Control | -51.097 | 3.431 | 704 | -14.893 | **< 0.001** |
| Annual Paclo Control - Annual MeJA Control | 2.760 | 2.297 | 704 | 1.202 | 0.998 |
| Annual Paclo Control - Perennial MeJA Control | -50.694 | 3.155 | 704 | -16.066 | **< 0.001** |
| Annual Paclo Control - Annual Control Exclosure | 4.273 | 3.536 | 704 | 1.208 | 0.998 |
| Annual Paclo Control - Perennial Control Exclosure | -51.176 | 3.895 | 704 | -13.139 | **< 0.001** |
| Annual Paclo Control - Annual GA Exclosure | 2.080 | 3.515 | 704 | 0.592 | 1.000 |
| Annual Paclo Control - Perennial GA Exclosure | -56.606 | 4.178 | 704 | -13.549 | **< 0.001** |
| Annual Paclo Control - Annual Paclo Exclosure | 1.589 | 3.536 | 704 | 0.449 | 1.000 |
| Annual Paclo Control - Perennial Paclo Exclosure | -60.383 | 4.196 | 704 | -14.389 | **< 0.001** |
| Annual Paclo Control - Annual MeJA Exclosure | 1.141 | 3.526 | 704 | 0.324 | 1.000 |
| Annual Paclo Control - Perennial MeJA Exclosure | -52.138 | 3.923 | 704 | -13.290 | **< 0.001** |
| Perennial Paclo Control - Annual MeJA Control | 53.857 | 3.465 | 704 | 15.545 | **< 0.001** |
| Perennial Paclo Control - Perennial MeJA Control | 0.403 | 3.616 | 704 | 0.111 | 1.000 |
| Perennial Paclo Control - Annual Control Exclosure | 55.370 | 4.385 | 704 | 12.627 | **< 0.001** |
| Perennial Paclo Control - Perennial Control Exclosure | -0.079 | 4.284 | 704 | -0.018 | 1.000 |
| Perennial Paclo Control - Annual GA Exclosure | 53.177 | 4.368 | 704 | 12.174 | **< 0.001** |
| Perennial Paclo Control - Perennial GA Exclosure | -5.509 | 4.542 | 704 | -1.213 | 0.998 |
| Perennial Paclo Control - Annual Paclo Exclosure | 52.686 | 4.384 | 704 | 12.017 | **< 0.001** |
| Perennial Paclo Control - Perennial Paclo Exclosure | -9.286 | 4.559 | 704 | -2.037 | 0.804 |
| Perennial Paclo Control - Annual MeJA Exclosure | 52.238 | 4.376 | 704 | 11.937 | **< 0.001** |
| Perennial Paclo Control - Perennial MeJA Exclosure | -1.041 | 4.304 | 704 | -0.242 | 1.000 |
| Annual MeJA Control - Perennial MeJA Control | -53.454 | 3.193 | 704 | -16.741 | **< 0.001** |
| Annual MeJA Control - Annual Control Exclosure | 1.513 | 3.570 | 704 | 0.424 | 1.000 |
| Annual MeJA Control - Perennial Control Exclosure | -53.936 | 3.926 | 704 | -13.740 | **< 0.001** |
| Annual MeJA Control - Annual GA Exclosure | -0.680 | 3.548 | 704 | -0.192 | 1.000 |
| Annual MeJA Control - Perennial GA Exclosure | -59.365 | 4.207 | 704 | -14.113 | **< 0.001** |
| Annual MeJA Control - Annual Paclo Exclosure | -1.171 | 3.570 | 704 | -0.328 | 1.000 |
| Annual MeJA Control - Perennial Paclo Exclosure | -63.143 | 4.225 | 704 | -14.945 | **< 0.001** |
| Annual MeJA Control - Annual MeJA Exclosure | -1.619 | 3.559 | 704 | -0.455 | 1.000 |

| | | | | | |
|---|---|---|---|---|---|
| Annual MeJA Control - Perennial MeJA Exclosure | -54.898 | 3.953 | 704 | -13.886 | < **0.001** |
| Perennial MeJA Control - Annual Control Exclosure | 54.967 | 4.173 | 704 | 13.173 | < **0.001** |
| Perennial MeJA Control - Perennial Control Exclosure | -0.482 | 4.062 | 704 | -0.119 | 1.000 |
| Perennial MeJA Control - Annual GA Exclosure | 52.774 | 4.155 | 704 | 12.700 | < **0.001** |
| Perennial MeJA Control - Perennial GA Exclosure | -5.912 | 4.327 | 704 | -1.366 | 0.993 |
| Perennial MeJA Control - Annual Paclo Exclosure | 52.283 | 4.172 | 704 | 12.532 | < **0.001** |
| Perennial MeJA Control - Perennial Paclo Exclosure | -9.689 | 4.361 | 704 | -2.222 | 0.682 |
| Perennial MeJA Control - Annual MeJA Exclosure | 51.835 | 4.163 | 704 | 12.450 | < **0.001** |
| Perennial MeJA Control - Perennial MeJA Exclosure | -1.444 | 4.095 | 704 | -0.353 | 1.000 |
| Annual Control Exclosure - Perennial Control Exclosure | -55.449 | 3.190 | 704 | -17.384 | < **0.001** |
| Annual Control Exclosure - Annual GA Exclosure | -2.193 | 2.714 | 704 | -0.808 | 1.000 |
| Annual Control Exclosure - Perennial GA Exclosure | -60.879 | 3.529 | 704 | -17.250 | < **0.001** |
| Annual Control Exclosure - Annual Paclo Exclosure | -2.684 | 2.741 | 704 | -0.979 | 1.000 |
| Annual Control Exclosure - Perennial Paclo Exclosure | -64.656 | 3.553 | 704 | -18.200 | < **0.001** |
| Annual Control Exclosure - Annual MeJA Exclosure | -3.132 | 2.728 | 704 | -1.148 | 0.999 |
| Annual Control Exclosure - Perennial MeJA Exclosure | -56.411 | 3.225 | 704 | -17.489 | < **0.001** |
| Perennial Control Exclosure - Annual GA Exclosure | 53.256 | 3.166 | 704 | 16.821 | < **0.001** |
| Perennial Control Exclosure - Perennial GA Exclosure | -5.429 | 3.384 | 704 | -1.604 | 0.967 |
| Perennial Control Exclosure - Annual Paclo Exclosure | 52.765 | 3.190 | 704 | 16.542 | < **0.001** |
| Perennial Control Exclosure - Perennial Paclo Exclosure | -9.207 | 3.432 | 704 | -2.683 | 0.344 |
| Perennial Control Exclosure - Annual MeJA Exclosure | 52.317 | 3.178 | 704 | 16.460 | < **0.001** |
| Perennial Control Exclosure - Perennial MeJA Exclosure | -0.962 | 3.082 | 704 | -0.312 | 1.000 |
| Annual GA Exclosure - Perennial GA Exclosure | -58.686 | 3.508 | 704 | -16.727 | < **0.001** |
| Annual GA Exclosure - Annual Paclo Exclosure | -0.491 | 2.714 | 704 | -0.181 | 1.000 |
| Annual GA Exclosure - Perennial Paclo Exclosure | -62.463 | 3.532 | 704 | -17.687 | < **0.001** |
| Annual GA Exclosure - Annual MeJA Exclosure | -0.939 | 2.700 | 704 | -0.348 | 1.000 |
| Annual GA Exclosure - Perennial MeJA Exclosure | -54.218 | 3.202 | 704 | -16.935 | < **0.001** |
| Perennial GA Exclosure - Annual Paclo Exclosure | 58.194 | 3.529 | 704 | 16.492 | < **0.001** |
| Perennial GA Exclosure - Perennial Paclo Exclosure | -3.778 | 3.747 | 704 | -1.008 | 1.000 |
| Perennial GA Exclosure - Annual MeJA Exclosure | 57.746 | 3.519 | 704 | 16.412 | < **0.001** |
| Perennial GA Exclosure - Perennial MeJA Exclosure | 4.468 | 3.430 | 704 | 1.303 | 0.996 |
| Annual Paclo Exclosure - Perennial Paclo Exclosure | -61.972 | 3.550 | 704 | -17.458 | < **0.001** |
| Annual Paclo Exclosure - Annual MeJA Exclosure | -0.448 | 2.727 | 704 | -0.164 | 1.000 |
| Annual Paclo Exclosure - Perennial MeJA Exclosure | -53.727 | 3.223 | 704 | -16.669 | < **0.001** |
| Perennial Paclo Exclosure - Annual MeJA Exclosure | 61.524 | 3.540 | 704 | 17.382 | < **0.001** |
| Perennial Paclo Exclosure - Perennial MeJA Exclosure | 8.245 | 3.457 | 704 | 2.385 | 0.559 |

| | | | | | |
|---|---|---|---|---|---|
| Annual MeJA Exclosure - Perennial MeJA Exclosure | -53.279 | 3.212 | 704 | -16.590 | **< 0.001** |

**Table S6**. Tukey post-hoc contrasts for the probability of herbivory at the coastal site, Bodega Marine Reserve. Minimum adequate model: Days to flowering = Ecotype + Hormone Treatment + Exclosure Treatment + Ecotype x Exclosure + Ecotype x Hormone treatment + (1|maternal family) + (1|Plot). *P*-values < 0.05 in bold. Contrasts are structured: Ecotype, Hormone treatment, Exclosure treatment vs Ecotype, Hormone treatment, Exclosure treatment. For hormone treatment: control = no-hormone, GA = $GA_3$ treatment, MeJA = methyl jasmonate treatment, and Paclo = paclobutrazol. For exclosure treatment: control = open structures, exclosures = mesh exclosures.

| **contrast** | **estimate** | **SE** | ***df*** | ***z*-ratio** | ***p*-value** |
|---|---|---|---|---|---|
| Annual Control Control - Perennial Control Control | -5.939 | 0.480 | Inf | -12.367 | **<0.001** |
| Annual Control Control - Annual GA Control | 3.611 | 0.471 | Inf | 7.666 | **<0.001** |
| Annual Control Control - Perennial GA Control | -2.328 | 0.527 | Inf | -4.420 | **0.001** |
| Annual Control Control - Annual Paclo Control | -0.397 | 0.462 | Inf | -0.858 | 1.000 |
| Annual Control Control - Perennial Paclo Control | -6.336 | 0.703 | Inf | -9.018 | **<0.001** |
| Annual Control Control - Annual MeJA Control | 0.313 | 0.447 | Inf | 0.701 | 1.000 |
| Annual Control Control - Perennial MeJA Control | -5.626 | 0.632 | Inf | -8.902 | **<0.001** |
| Annual Control Control - Annual Control Exclosure | -1.352 | 0.799 | Inf | -1.693 | 0.949 |
| Annual Control Control - Perennial Control Exclosure | -3.939 | 0.778 | Inf | -5.063 | **<0.001** |
| Annual Control Control - Annual GA Exclosure | 0.199 | 0.838 | Inf | 0.238 | 1.000 |
| Annual Control Control - Perennial GA Exclosure | -2.388 | 0.792 | Inf | -3.014 | 0.163 |
| Annual Control Control - Annual Paclo Exclosure | -1.246 | 0.800 | Inf | -1.557 | 0.976 |
| Annual Control Control - Perennial Paclo Exclosure | -3.833 | 0.777 | Inf | -4.934 | **<0.001** |
| Annual Control Control - Annual MeJA Exclosure | -0.677 | 0.812 | Inf | -0.833 | 1.000 |
| Annual Control Control - Perennial MeJA Exclosure | -3.263 | 0.777 | Inf | -4.202 | **0.003** |
| Perennial Control Control - Annual GA Control | 9.550 | 0.792 | Inf | 12.055 | **<0.001** |
| Perennial Control Control - Perennial GA Control | 3.611 | 0.471 | Inf | 7.666 | **<0.001** |
| Perennial Control Control - Annual Paclo Control | 5.542 | 0.628 | Inf | 8.823 | **<0.001** |
| Perennial Control Control - Perennial Paclo Control | -0.397 | 0.462 | Inf | -0.858 | 1.000 |
| Perennial Control Control - Annual MeJA Control | 6.252 | 0.679 | Inf | 9.203 | **<0.001** |
| Perennial Control Control - Perennial MeJA Control | 0.313 | 0.447 | Inf | 0.701 | 1.000 |
| Perennial Control Control - Annual Control Exclosure | 4.587 | 0.752 | Inf | 6.103 | **<0.001** |
| Perennial Control Control - Perennial Control Exclosure | 2.000 | 0.726 | Inf | 2.753 | 0.296 |
| Perennial Control Control - Annual GA Exclosure | 6.138 | 0.795 | Inf | 7.723 | **<0.001** |
| Perennial Control Control - Perennial GA Exclosure | 3.551 | 0.743 | Inf | 4.778 | **<0.001** |
| Perennial Control Control - Annual Paclo Exclosure | 4.693 | 0.753 | Inf | 6.231 | **<0.001** |
| Perennial Control Control - Perennial Paclo Exclosure | 2.106 | 0.725 | Inf | 2.904 | 0.213 |
| Perennial Control Control - Annual MeJA Exclosure | 5.262 | 0.766 | Inf | 6.866 | **<0.001** |

| | | | | | |
|---|---|---|---|---|---|
| Perennial Control Control - Perennial MeJA Exclosure | 2.675 | 0.726 | Inf | 3.685 | **0.021** |
| Annual GA Control - Perennial GA Control | -5.939 | 0.480 | Inf | -12.367 | **<0.001** |
| Annual GA Control - Annual Paclo Control | -4.008 | 0.498 | Inf | -8.053 | **<0.001** |
| Annual GA Control - Perennial Paclo Control | -9.947 | 0.838 | Inf | -11.864 | **<0.001** |
| Annual GA Control - Annual MeJA Control | -3.298 | 0.455 | Inf | -7.253 | **<0.001** |
| Annual GA Control - Perennial MeJA Control | -9.237 | 0.763 | Inf | -12.113 | **<0.001** |
| Annual GA Control - Annual Control Exclosure | -4.964 | 0.876 | Inf | -5.668 | **<0.001** |
| Annual GA Control - Perennial Control Exclosure | -7.550 | 0.858 | Inf | -8.797 | **<0.001** |
| Annual GA Control - Annual GA Exclosure | -3.412 | 0.911 | Inf | -3.746 | **0.017** |
| Annual GA Control - Perennial GA Exclosure | -5.999 | 0.870 | Inf | -6.893 | **<0.001** |
| Annual GA Control - Annual Paclo Exclosure | -4.857 | 0.877 | Inf | -5.538 | **<0.001** |
| Annual GA Control - Perennial Paclo Exclosure | -7.444 | 0.857 | Inf | -8.684 | **<0.001** |
| Annual GA Control - Annual MeJA Exclosure | -4.288 | 0.887 | Inf | -4.832 | **<0.001** |
| Annual GA Control - Perennial MeJA Exclosure | -6.875 | 0.857 | Inf | -8.024 | **<0.001** |
| Perennial GA Control - Annual Paclo Control | 1.931 | 0.504 | Inf | 3.834 | **0.012** |
| Perennial GA Control - Perennial Paclo Control | -4.008 | 0.498 | Inf | -8.053 | **<0.001** |
| Perennial GA Control - Annual MeJA Control | 2.641 | 0.542 | Inf | 4.876 | **<0.001** |
| Perennial GA Control - Perennial MeJA Control | -3.298 | 0.455 | Inf | -7.253 | **<0.001** |
| Perennial GA Control - Annual Control Exclosure | 0.975 | 0.720 | Inf | 1.354 | 0.994 |
| Perennial GA Control - Perennial Control Exclosure | -1.612 | 0.696 | Inf | -2.316 | 0.611 |
| Perennial GA Control - Annual GA Exclosure | 2.527 | 0.764 | Inf | 3.306 | 0.073 |
| Perennial GA Control - Perennial GA Exclosure | -0.060 | 0.712 | Inf | -0.084 | 1.000 |
| Perennial GA Control - Annual Paclo Exclosure | 1.082 | 0.722 | Inf | 1.498 | 0.983 |
| Perennial GA Control - Perennial Paclo Exclosure | -1.505 | 0.694 | Inf | -2.168 | 0.720 |
| Perennial GA Control - Annual MeJA Exclosure | 1.651 | 0.735 | Inf | 2.245 | 0.665 |
| Perennial GA Control - Perennial MeJA Exclosure | -0.936 | 0.695 | Inf | -1.347 | 0.994 |
| Annual Paclo Control - Perennial Paclo Control | -5.939 | 0.480 | Inf | -12.367 | **<0.001** |
| Annual Paclo Control - Annual MeJA Control | 0.710 | 0.462 | Inf | 1.537 | 0.978 |
| Annual Paclo Control - Perennial MeJA Control | -5.229 | 0.603 | Inf | -8.676 | **<0.001** |
| Annual Paclo Control - Annual Control Exclosure | -0.955 | 0.784 | Inf | -1.219 | 0.998 |
| Annual Paclo Control - Perennial Control Exclosure | -3.542 | 0.762 | Inf | -4.646 | **<0.001** |
| Annual Paclo Control - Annual GA Exclosure | 0.596 | 0.824 | Inf | 0.724 | 1.000 |
| Annual Paclo Control - Perennial GA Exclosure | -1.991 | 0.777 | Inf | -2.563 | 0.425 |
| Annual Paclo Control - Annual Paclo Exclosure | -0.849 | 0.785 | Inf | -1.082 | 1.000 |
| Annual Paclo Control - Perennial Paclo Exclosure | -3.436 | 0.761 | Inf | -4.514 | **<0.001** |
| Annual Paclo Control - Annual MeJA Exclosure | -0.280 | 0.797 | Inf | -0.351 | 1.000 |
| Annual Paclo Control - Perennial MeJA Exclosure | -2.867 | 0.761 | Inf | -3.766 | 0.016 |
| Perennial Paclo Control - Annual MeJA Control | 6.649 | 0.724 | Inf | 9.182 | **<0.001** |
| Perennial Paclo Control - Perennial MeJA Control | 0.710 | 0.462 | Inf | 1.537 | 0.978 |

| | | | | | |
|---|---|---|---|---|---|
| Perennial Paclo Control - Annual Control Exclosure | 4.983 | 0.768 | Inf | 6.485 | **<0.001** |
| Perennial Paclo Control - Perennial Control Exclosure | 2.397 | 0.744 | Inf | 3.222 | 0.093 |
| Perennial Paclo Control - Annual GA Exclosure | 6.535 | 0.811 | Inf | 8.059 | **<0.001** |
| Perennial Paclo Control - Perennial GA Exclosure | 3.948 | 0.760 | Inf | 5.192 | **<0.001** |
| Perennial Paclo Control - Annual Paclo Exclosure | 5.090 | 0.770 | Inf | 6.610 | **<0.001** |
| Perennial Paclo Control - Perennial Paclo Exclosure | 2.503 | 0.743 | Inf | 3.370 | 0.060 |
| Perennial Paclo Control - Annual MeJA Exclosure | 5.659 | 0.783 | Inf | 7.227 | **<0.001** |
| Perennial Paclo Control - Perennial MeJA Exclosure | 3.072 | 0.743 | Inf | 4.132 | **0.004** |
| Annual MeJA Control - Perennial MeJA Control | -5.939 | 0.480 | Inf | -12.367 | **<0.001** |
| Annual MeJA Control - Annual Control Exclosure | -1.665 | 0.809 | Inf | -2.059 | 0.791 |
| Annual MeJA Control - Perennial Control Exclosure | -4.252 | 0.788 | Inf | -5.395 | **<0.001** |
| Annual MeJA Control - Annual GA Exclosure | -0.114 | 0.847 | Inf | -0.134 | 1.000 |
| Annual MeJA Control - Perennial GA Exclosure | -2.701 | 0.802 | Inf | -3.367 | 0.060 |
| Annual MeJA Control - Annual Paclo Exclosure | -1.559 | 0.810 | Inf | -1.925 | 0.865 |
| Annual MeJA Control - Perennial Paclo Exclosure | -4.146 | 0.787 | Inf | -5.268 | **<0.001** |
| Annual MeJA Control - Annual MeJA Exclosure | -0.990 | 0.822 | Inf | -1.205 | 0.998 |
| Annual MeJA Control - Perennial MeJA Exclosure | -3.577 | 0.787 | Inf | -4.545 | **<0.001** |
| Perennial MeJA Control - Annual Control Exclosure | 4.274 | 0.741 | Inf | 5.764 | **<0.001** |
| Perennial MeJA Control - Perennial Control Exclosure | 1.687 | 0.716 | Inf | 2.356 | 0.581 |
| Perennial MeJA Control - Annual GA Exclosure | 5.825 | 0.785 | Inf | 7.418 | **<0.001** |
| Perennial MeJA Control - Perennial GA Exclosure | 3.238 | 0.733 | Inf | 4.417 | **0.001** |
| Perennial MeJA Control - Annual Paclo Exclosure | 4.380 | 0.743 | Inf | 5.894 | **<0.001** |
| Perennial MeJA Control - Perennial Paclo Exclosure | 1.793 | 0.715 | Inf | 2.508 | 0.465 |
| Perennial MeJA Control - Annual MeJA Exclosure | 4.949 | 0.757 | Inf | 6.542 | **<0.001** |
| Perennial MeJA Control - Perennial MeJA Exclosure | 2.362 | 0.716 | Inf | 3.301 | 0.074 |
| Annual Control Exclosure - Perennial Control Exclosure | -2.587 | 0.315 | Inf | -8.200 | **<0.001** |
| Annual Control Exclosure - Annual GA Exclosure | 1.551 | 0.405 | Inf | 3.833 | **0.012** |
| Annual Control Exclosure - Perennial GA Exclosure | -1.035 | 0.471 | Inf | -2.200 | 0.697 |
| Annual Control Exclosure - Annual Paclo Exclosure | 0.106 | 0.356 | Inf | 0.299 | 1.000 |
| Annual Control Exclosure - Perennial Paclo Exclosure | -2.481 | 0.471 | Inf | -5.266 | <0.001 |
| Annual Control Exclosure - Annual MeJA Exclosure | 0.676 | 0.365 | Inf | 1.853 | 0.897 |
| Annual Control Exclosure - Perennial MeJA Exclosure | -1.911 | 0.458 | Inf | -4.176 | **0.003** |
| Perennial Control Exclosure - Annual GA Exclosure | 4.138 | 0.553 | Inf | 7.490 | **<0.001** |
| Perennial Control Exclosure - Perennial GA Exclosure | 1.551 | 0.405 | Inf | 3.833 | **0.012** |
| Perennial Control Exclosure - Annual Paclo Exclosure | 2.693 | 0.480 | Inf | 5.615 | **<0.001** |
| Perennial Control Exclosure - Perennial Paclo Exclosure | 0.106 | 0.356 | Inf | 0.299 | 1.000 |
| Perennial Control Exclosure - Annual MeJA Exclosure | 3.262 | 0.506 | Inf | 6.454 | **<0.001** |

| | | | | | |
|---|---|---|---|---|---|
| Perennial Control Exclosure - Perennial MeJA Exclosure | 0.676 | 0.365 | Inf | 1.853 | 0.897 |
| Annual GA Exclosure - Perennial GA Exclosure | -2.587 | 0.315 | Inf | -8.200 | **<0.001** |
| Annual GA Exclosure - Annual Paclo Exclosure | -1.445 | 0.403 | Inf | -3.585 | **0.030** |
| Annual GA Exclosure - Perennial Paclo Exclosure | -4.032 | 0.548 | Inf | -7.363 | **<0.001** |
| Annual GA Exclosure - Annual MeJA Exclosure | -0.876 | 0.405 | Inf | -2.160 | 0.725 |
| Annual GA Exclosure - Perennial MeJA Exclosure | -3.463 | 0.532 | Inf | -6.512 | **<0.001** |
| Perennial GA Exclosure - Annual Paclo Exclosure | 1.142 | 0.473 | Inf | 2.411 | 0.539 |
| Perennial GA Exclosure - Perennial Paclo Exclosure | -1.445 | 0.403 | Inf | -3.585 | **0.030** |
| Perennial GA Exclosure - Annual MeJA Exclosure | 1.711 | 0.495 | Inf | 3.457 | **0.045** |
| Perennial GA Exclosure - Perennial MeJA Exclosure | -0.876 | 0.405 | Inf | -2.160 | 0.725 |
| Annual Paclo Exclosure - Perennial Paclo Exclosure | -2.587 | 0.315 | Inf | -8.200 | **<0.001** |
| Annual Paclo Exclosure - Annual MeJA Exclosure | 0.569 | 0.364 | Inf | 1.566 | 0.974 |
| Annual Paclo Exclosure - Perennial MeJA Exclosure | -2.018 | 0.461 | Inf | -4.374 | **0.001** |
| Perennial Paclo Exclosure - Annual MeJA Exclosure | 3.156 | 0.501 | Inf | 6.304 | **<0.001** |
| Perennial Paclo Exclosure - Perennial MeJA Exclosure | 0.569 | 0.364 | Inf | 1.566 | 0.974 |
| Annual MeJA Exclosure - Perennial MeJA Exclosure | -2.587 | 0.315 | Inf | -8.200 | **<0.001** |

**Table S7**. Tukey post-hoc contrasts for the probability of herbivory at the inland site, Pepperwood Preserve. Minimum adequate model: Herbivory (presence/absence) = Ecotype + Hormone Treatment + Exclosure Treatment + Ecotype x Exclosure + Ecotype x Hormone + Hormone x Exclosure + Ecotype x Hormone x Exclosure + (1|maternal family) + (1|Plot). *P*-values < 0.05 in bold. Contrasts are structured: Ecotype, Hormone treatment, Exclosure treatment vs Ecotype, Hormone treatment, Exclosure treatment. For hormone treatment: control = no-hormone, GA = $GA_3$ treatment, MeJA = methyl jasmonate treatment, and Paclo = paclobutrazol. For exclosure treatment: control = open structures, exclosures = mesh exclosures.

| **contrast** | **estimate** | **SE** | ***df*** | ***z*-ratio** | ***p*-value** |
|---|---|---|---|---|---|
| Annual Control Control - Perennial Control Control | -2.542 | 0.474 | Inf | -5.358 | **< 0.001** |
| Annual Control Control - Annual GA Control | -0.602 | 0.514 | Inf | -1.171 | 0.999 |
| Annual Control Control - Perennial GA Control | -1.739 | 0.476 | Inf | -3.651 | **0.024** |
| Annual Control Control - Annual Paclo Control | -0.723 | 0.508 | Inf | -1.424 | 0.990 |
| Annual Control Control - Perennial Paclo Control | -1.763 | 0.477 | Inf | -3.698 | **0.020** |
| Annual Control Control - Annual MeJA Control | 0.186 | 0.588 | Inf | 0.317 | 1.000 |
| Annual Control Control - Perennial MeJA Control | -2.046 | 0.473 | Inf | -4.323 | **0.002** |
| Annual Control Control - Annual Control Exclosure | -0.750 | 0.802 | Inf | -0.935 | 1.000 |
| Annual Control Control - Perennial Control Exclosure | -1.779 | 0.782 | Inf | -2.274 | 0.644 |
| Annual Control Control - Annual GA Exclosure | -0.279 | 0.822 | Inf | -0.339 | 1.000 |
| Annual Control Control - Perennial GA Exclosure | -0.966 | 0.796 | Inf | -1.214 | 0.998 |

| | | | | | |
|---|---|---|---|---|---|
| Annual Control Control - Annual Paclo Exclosure | 18.817 | 4257.700 | Inf | 0.004 | 1.000 |
| Annual Control Control - Perennial Paclo Exclosure | -1.706 | 0.783 | Inf | -2.180 | 0.712 |
| Annual Control Control - Annual MeJA Exclosure | -0.112 | 0.834 | Inf | -0.134 | 1.000 |
| Annual Control Control - Perennial MeJA Exclosure | -0.858 | 0.801 | Inf | -1.071 | 1.000 |
| Perennial Control Control - Annual GA Control | 1.940 | 0.408 | Inf | 4.760 | **< 0.001** |
| Perennial Control Control - Perennial GA Control | 0.803 | 0.346 | Inf | 2.318 | 0.610 |
| Perennial Control Control - Annual Paclo Control | 1.818 | 0.400 | Inf | 4.551 | **< 0.001** |
| Perennial Control Control - Perennial Paclo Control | 0.779 | 0.346 | Inf | 2.249 | 0.662 |
| Perennial Control Control - Annual MeJA Control | 2.728 | 0.498 | Inf | 5.478 | **< 0.001** |
| Perennial Control Control - Perennial MeJA Control | 0.496 | 0.340 | Inf | 1.459 | 0.987 |
| Perennial Control Control - Annual Control Exclosure | 1.791 | 0.738 | Inf | 2.429 | 0.526 |
| Perennial Control Control - Perennial Control Exclosure | 0.763 | 0.711 | Inf | 1.074 | 1.000 |
| Perennial Control Control - Annual GA Exclosure | 2.263 | 0.759 | Inf | 2.980 | 0.177 |
| Perennial Control Control - Perennial GA Exclosure | 1.576 | 0.725 | Inf | 2.173 | 0.717 |
| Perennial Control Control - Annual Paclo Exclosure | 21.358 | 4257.700 | Inf | 0.005 | 1.000 |
| Perennial Control Control - Perennial Paclo Exclosure | 0.836 | 0.711 | Inf | 1.176 | 0.999 |
| Perennial Control Control - Annual MeJA Exclosure | 2.430 | 0.773 | Inf | 3.145 | 0.115 |
| Perennial Control Control - Perennial MeJA Exclosure | 1.684 | 0.731 | Inf | 2.302 | 0.622 |
| Annual GA Control - Perennial GA Control | -1.137 | 0.410 | Inf | -2.774 | 0.284 |
| Annual GA Control - Annual Paclo Control | -0.122 | 0.446 | Inf | -0.272 | 1.000 |
| Annual GA Control - Perennial Paclo Control | -1.161 | 0.410 | Inf | -2.830 | 0.252 |
| Annual GA Control - Annual MeJA Control | 0.788 | 0.536 | Inf | 1.471 | 0.986 |
| Annual GA Control - Perennial MeJA Control | -1.444 | 0.406 | Inf | -3.555 | **0.033** |
| Annual GA Control - Annual Control Exclosure | -0.148 | 0.765 | Inf | -0.194 | 1.000 |
| Annual GA Control - Perennial Control Exclosure | -1.177 | 0.744 | Inf | -1.582 | 0.972 |
| Annual GA Control - Annual GA Exclosure | 0.323 | 0.786 | Inf | 0.411 | 1.000 |
| Annual GA Control - Perennial GA Exclosure | -0.364 | 0.758 | Inf | -0.480 | 1.000 |
| Annual GA Control - Annual Paclo Exclosure | 19.418 | 4257.700 | Inf | 0.005 | 1.000 |
| Annual GA Control - Perennial Paclo Exclosure | -1.104 | 0.744 | Inf | -1.484 | 0.984 |
| Annual GA Control - Annual MeJA Exclosure | 0.490 | 0.798 | Inf | 0.613 | 1.000 |
| Annual GA Control - Perennial MeJA Exclosure | -0.256 | 0.764 | Inf | -0.336 | 1.000 |
| Perennial GA Control - Annual Paclo Control | 1.016 | 0.402 | Inf | 2.525 | 0.453 |
| Perennial GA Control - Perennial Paclo Control | -0.024 | 0.351 | Inf | -0.068 | 1.000 |
| Perennial GA Control - Annual MeJA Control | 1.925 | 0.500 | Inf | 3.851 | **0.011** |
| Perennial GA Control - Perennial MeJA Control | -0.307 | 0.345 | Inf | -0.889 | 1.000 |
| Perennial GA Control - Annual Control Exclosure | 0.989 | 0.739 | Inf | 1.337 | 0.995 |

| | | | | | |
|---|---|---|---|---|---|
| Perennial GA Control - Perennial Control Exclosure | -0.040 | 0.713 | Inf | -0.056 | 1.000 |
| Perennial GA Control - Annual GA Exclosure | 1.460 | 0.761 | Inf | 1.918 | 0.868 |
| Perennial GA Control - Perennial GA Exclosure | 0.773 | 0.727 | Inf | 1.063 | 1.000 |
| Perennial GA Control - Annual Paclo Exclosure | 20.556 | 4257.700 | Inf | 0.005 | 1.000 |
| Perennial GA Control - Perennial Paclo Exclosure | 0.033 | 0.713 | Inf | 0.046 | 1.000 |
| Perennial GA Control - Annual MeJA Exclosure | 1.627 | 0.774 | Inf | 2.101 | 0.765 |
| Perennial GA Control - Perennial MeJA Exclosure | 0.881 | 0.733 | Inf | 1.201 | 0.998 |
| Annual Paclo Control - Perennial Paclo Control | -1.040 | 0.403 | Inf | -2.582 | 0.411 |
| Annual Paclo Control - Annual MeJA Control | 0.910 | 0.530 | Inf | 1.716 | 0.943 |
| Annual Paclo Control - Perennial MeJA Control | -1.322 | 0.398 | Inf | -3.320 | 0.069 |
| Annual Paclo Control - Annual Control Exclosure | -0.027 | 0.761 | Inf | -0.035 | 1.000 |
| Annual Paclo Control - Perennial Control Exclosure | -1.056 | 0.740 | Inf | -1.427 | 0.989 |
| Annual Paclo Control - Annual GA Exclosure | 0.444 | 0.782 | Inf | 0.569 | 1.000 |
| Annual Paclo Control - Perennial GA Exclosure | -0.242 | 0.754 | Inf | -0.322 | 1.000 |
| Annual Paclo Control - Annual Paclo Exclosure | 19.540 | 4257.700 | Inf | 0.005 | 1.000 |
| Annual Paclo Control - Perennial Paclo Exclosure | -0.983 | 0.740 | Inf | -1.328 | 0.995 |
| Annual Paclo Control - Annual MeJA Exclosure | 0.611 | 0.794 | Inf | 0.769 | 1.000 |
| Annual Paclo Control - Perennial MeJA Exclosure | -0.135 | 0.759 | Inf | -0.177 | 1.000 |
| Perennial Paclo Control - Annual MeJA Control | 1.949 | 0.500 | Inf | 3.896 | **0.010** |
| Perennial Paclo Control - Perennial MeJA Control | -0.283 | 0.345 | Inf | -0.819 | 1.000 |
| Perennial Paclo Control - Annual Control Exclosure | 1.013 | 0.740 | Inf | 1.369 | 0.993 |
| Perennial Paclo Control - Perennial Control Exclosure | -0.016 | 0.713 | Inf | -0.022 | 1.000 |
| Perennial Paclo Control - Annual GA Exclosure | 1.484 | 0.761 | Inf | 1.950 | 0.853 |
| Perennial Paclo Control - Perennial GA Exclosure | 0.797 | 0.727 | Inf | 1.096 | 0.999 |
| Perennial Paclo Control - Annual Paclo Exclosure | 20.579 | 4257.700 | Inf | 0.005 | 1.000 |
| Perennial Paclo Control - Perennial Paclo Exclosure | 0.057 | 0.713 | Inf | 0.080 | 1.000 |
| Perennial Paclo Control - Annual MeJA Exclosure | 1.651 | 0.774 | Inf | 2.132 | 0.745 |
| Perennial Paclo Control - Perennial MeJA Exclosure | 0.905 | 0.733 | Inf | 1.234 | 0.998 |
| Annual MeJA Control - Perennial MeJA Control | -2.232 | 0.497 | Inf | -4.492 | **< 0.001** |
| Annual MeJA Control - Annual Control Exclosure | -0.937 | 0.816 | Inf | -1.147 | 0.999 |
| Annual MeJA Control - Perennial Control Exclosure | -1.965 | 0.797 | Inf | -2.466 | 0.497 |
| Annual MeJA Control - Annual GA Exclosure | -0.465 | 0.836 | Inf | -0.556 | 1.000 |
| Annual MeJA Control - Perennial GA Exclosure | -1.152 | 0.810 | Inf | -1.423 | 0.990 |
| Annual MeJA Control - Annual Paclo Exclosure | 18.630 | 4257.700 | Inf | 0.004 | 1.000 |
| Annual MeJA Control - Perennial Paclo Exclosure | -1.892 | 0.797 | Inf | -2.374 | 0.568 |
| Annual MeJA Control - Annual MeJA Exclosure | -0.298 | 0.848 | Inf | -0.352 | 1.000 |

| | | | | | |
|---|---|---|---|---|---|
| Annual MeJA Control - Perennial MeJA Exclosure | -1.044 | 0.815 | Inf | -1.281 | 0.997 |
| Perennial MeJA Control - Annual Control Exclosure | 1.295 | 0.737 | Inf | 1.757 | 0.931 |
| Perennial MeJA Control - Perennial Control Exclosure | 0.267 | 0.710 | Inf | 0.376 | 1.000 |
| Perennial MeJA Control - Annual GA Exclosure | 1.767 | 0.759 | Inf | 2.328 | 0.602 |
| Perennial MeJA Control - Perennial GA Exclosure | 1.080 | 0.725 | Inf | 1.490 | 0.984 |
| Perennial MeJA Control - Annual Paclo Exclosure | 20.862 | 4257.700 | Inf | 0.005 | 1.000 |
| Perennial MeJA Control - Perennial Paclo Exclosure | 0.340 | 0.710 | Inf | 0.478 | 1.000 |
| Perennial MeJA Control - Annual MeJA Exclosure | 1.934 | 0.772 | Inf | 2.505 | 0.468 |
| Perennial MeJA Control - Perennial MeJA Exclosure | 1.188 | 0.731 | Inf | 1.625 | 0.964 |
| Annual Control Exclosure - Perennial Control Exclosure | -1.029 | 0.493 | Inf | -2.088 | 0.773 |
| Annual Control Exclosure - Annual GA Exclosure | 0.471 | 0.544 | Inf | 0.867 | 1.000 |
| Annual Control Exclosure - Perennial GA Exclosure | -0.216 | 0.506 | Inf | -0.426 | 1.000 |
| Annual Control Exclosure - Annual Paclo Exclosure | 19.567 | 4257.700 | Inf | 0.005 | 1.000 |
| Annual Control Exclosure - Perennial Paclo Exclosure | -0.956 | 0.490 | Inf | -1.949 | 0.853 |
| Annual Control Exclosure - Annual MeJA Exclosure | 0.638 | 0.561 | Inf | 1.138 | 0.999 |
| Annual Control Exclosure - Perennial MeJA Exclosure | -0.108 | 0.513 | Inf | -0.210 | 1.000 |
| Perennial Control Exclosure - Annual GA Exclosure | 1.500 | 0.523 | Inf | 2.866 | 0.232 |
| Perennial Control Exclosure - Perennial GA Exclosure | 0.813 | 0.475 | Inf | 1.712 | 0.944 |
| Perennial Control Exclosure - Annual Paclo Exclosure | 20.595 | 4257.700 | Inf | 0.005 | 1.000 |
| Perennial Control Exclosure - Perennial Paclo Exclosure | 0.073 | 0.456 | Inf | 0.160 | 1.000 |
| Perennial Control Exclosure - Annual MeJA Exclosure | 1.667 | 0.542 | Inf | 3.076 | 0.139 |
| Perennial Control Exclosure - Perennial MeJA Exclosure | 0.921 | 0.483 | Inf | 1.906 | 0.874 |
| Annual GA Exclosure - Perennial GA Exclosure | -0.687 | 0.535 | Inf | -1.284 | 0.996 |
| Annual GA Exclosure - Annual Paclo Exclosure | 19.096 | 4257.700 | Inf | 0.004 | 1.000 |
| Annual GA Exclosure - Perennial Paclo Exclosure | -1.427 | 0.521 | Inf | -2.740 | 0.305 |
| Annual GA Exclosure - Annual MeJA Exclosure | 0.167 | 0.586 | Inf | 0.285 | 1.000 |
| Annual GA Exclosure - Perennial MeJA Exclosure | -0.579 | 0.541 | Inf | -1.070 | 1.000 |
| Perennial GA Exclosure - Annual Paclo Exclosure | 19.782 | 4257.700 | Inf | 0.005 | 1.000 |
| Perennial GA Exclosure - Perennial Paclo Exclosure | -0.740 | 0.473 | Inf | -1.566 | 0.974 |
| Perennial GA Exclosure - Annual MeJA Exclosure | 0.854 | 0.552 | Inf | 1.546 | 0.977 |
| Perennial GA Exclosure - Perennial MeJA Exclosure | 0.108 | 0.496 | Inf | 0.217 | 1.000 |
| Annual Paclo Exclosure - Perennial Paclo Exclosure | -20.523 | 4257.700 | Inf | -0.005 | 1.000 |
| Annual Paclo Exclosure - Annual MeJA Exclosure | -18.929 | 4257.700 | Inf | -0.004 | 1.000 |

| | | | | | |
|---|---|---|---|---|---|
| Annual Paclo Exclosure - Perennial MeJA Exclosure | -19.675 | 4257.700 | Inf | -0.005 | 1.000 |
| Perennial Paclo Exclosure - Annual MeJA Exclosure | 1.594 | 0.539 | Inf | 2.956 | 0.188 |
| Perennial Paclo Exclosure - Perennial MeJA Exclosure | 0.848 | 0.481 | Inf | 1.764 | 0.929 |
| Annual MeJA Exclosure - Perennial MeJA Exclosure | -0.746 | 0.558 | Inf | -1.336 | 0.995 |

**Table S8**. Tukey post-hoc contrasts for total PPG at the coastal site, Bodega Marine Reserve. Minimum adequate model: Total PPG = Hormone Treatment + Exclosure Treatment + Hormone treatment x Exclosure treatment + (1|maternal family) + (1|Plot). *P*-values < 0.05 in bold.Contrasts are structured: Ecotype, Hormone treatment vs Ecotype, Hormone treatment. For hormone treatment: control = no-hormone, GA = $GA_3$ treatment, MeJA = methyl jasmonate treatment, and Paclo = paclobutrazol.

| **contrast** | **estimate** | **SE** | ***df*** | ***z*-ratio** | ***p*-value** |
|---|---|---|---|---|---|
| Control Control - GA Control | 0.370 | 0.066 | Inf | 5.589 | **<0.001** |
| Control Control - MeJA Control | 0.033 | 0.053 | Inf | 0.618 | 0.999 |
| Control Control - Paclo Control | 0.056 | 0.053 | Inf | 1.049 | 0.967 |
| GA Control - MeJA Control | -0.338 | 0.0673 | Inf | -5.040 | **<0.001** |
| GA Control - Paclo Control | -0.314 | 0.067 | Inf | -4.674 | **0.001** |
| MeJA Control - Paclo Control | 0.023 | 0.0525 | Inf | 0.443 | 1 |
| Control Exclosure - GA Exclosure | 0.079 | 0.056 | Inf | 1.428 | 0.844 |
| Control Exclosure - MeJA Exclosure | -0.053 | 0.048 | Inf | -1.112 | 0.954 |
| Control Exclosure - Paclo Exclosure | 0.006 | 0.047 | Inf | 0.135 | 1 |
| GA Exclosure - MeJA Exclosure | -0.133 | 0.057 | Inf | -2.346 | 0.268 |
| GA Exclosure - Paclo Exclosure | -0.073 | 0.056 | Inf | -1.307 | 0.897 |
| MeJA Exclosure - Paclo Exclosure | 0.060 | 0.049 | Inf | 1.227 | 0.924 |
| Control Control - Control Exclosure | 0.178 | 0.112 | Inf | 1.586 | 0.759 |
| GA Control - GA Exclosure | -0.113 | 0.124 | Inf | -0.914 | 0.985 |
| MeJA Control - MeJA Exclosure | 0.092 | 0.113 | Inf | 0.822 | 0.992 |
| Paclo Control - Paclo Exclosure | 0.128 | 0.112 | Inf | 1.145 | 0.947 |

**Table S9**. Tukey post-hoc contrasts for total PPG at the inland site, Pepperwood Preserve. Minimum adequate model: Total PPG = Ecotype + Hormone treatment + Ecotype x Hormone treatment + (1|maternal family) + (1|Plot). *P*-values < 0.05 in bold. Contrasts are structured:

Ecotype, Hormone treatment vs Ecotype, Hormone treatment. For hormone treatment: control = no-hormone, GA = $GA_3$ treatment, MeJA = methyl jasmonate treatment, and Paclo = paclobutrazol.

| **contrast** | **estimate** | **SE** | ***df*** | ***z*-ratio** | ***p*-value** |
|---|---|---|---|---|---|
| Annual Control - Perennial Control | -0.792 | 0.062 | Inf | -12.809 | **< 0.001** |
| Annual Control - Annual GA | -0.185 | 0.058 | Inf | -3.196 | **0.030** |
| Annual Control - Perennial GA | -0.826 | 0.065 | Inf | -12.775 | **<0.001** |
| Annual Control - Annual MeJA | -0.219 | 0.054 | Inf | -4.113 | **0.001** |
| Annual Control - Perennial MeJA | -0.818 | 0.062 | Inf | -13.318 | **<0.001** |
| Annual Control - Annual Paclo | -0.024 | 0.054 | Inf | -0.450 | 1 |
| Annual Control - Perennial Paclo | -0.794 | 0.063 | Inf | -12.736 | **<0.001** |
| Perennial Control - Annual GA | 0.608 | 0.065 | Inf | 9.418 | **<0.001** |
| Perennial Control - Perennial GA | -0.034 | 0.043 | Inf | -0.797 | 0.993 |
| Perennial Control - Annual MeJA | 0.573 | 0.061 | Inf | 9.402 | **<0.001** |
| Perennial Control - Perennial MeJA | 0.234 | 0.080 | Inf | -0.830 | 0.991 |
| Perennial Control - Annual Paclo | 0.768 | 0.062 | Inf | 12.657 | **<0.001** |
| Perennial Control - Perennial Paclo | -0.001 | 0.040 | Inf | -0.072 | 1 |
| Annual GA - Perennial GA | -0.642 | 0.068 | Inf | -9.562 | **<0.001** |
| Annual GA - Annual MeJA | -0.035 | 0.057 | Inf | -0.621 | 0.999 |
| Annual GA - Perennial MeJA | -0.633 | 0.065 | Inf | -9.882 | **<0.001** |
| Annual GA - Annual Paclo | 0.160 | 0.057 | Inf | 2.816 | 0.091 |
| Annual GA - Perennial Paclo | -0.609 | 0.066 | Inf | -9.365 | **<0.001** |
| Perennial GA - Annual MeJA | 0.607 | 0.064 | Inf | 9.518 | **<0.001** |
| Perennial GA - Perennial MeJA | 0.009 | 0.044 | Inf | 0.035 | 1 |
| Perennial GA - Annual Paclo | 0.802 | 0.064 | Inf | 12.549 | **<0.001** |
| Perennial GA - Perennial Paclo | 0.033 | 0.044 | Inf | 0.710 | 0.997 |
| Annual MeJA - Perennial MeJA | -0.598 | 0.062 | Inf | -9.914 | **<0.001** |
| Annual MeJA - Annual Paclo | 0.195 | 0.053 | Inf | 3.706 | **0.005** |
| Annual MeJA - Perennial Paclo | -0.574 | 0.062 | Inf | -9.367 | **<0.001** |
| Perennial MeJA - Annual Paclo | 0.793 | 0.062 | Inf | 13.063 | **<0.001** |
| Perennial MeJA - Perennial Paclo | 0.024 | 0.040 | Inf | 0.594 | 0.999 |
| Annual Paclo - Perennial Paclo | -0.769 | 0.062 | Inf | -12.354 | **<0.001** |

**Table S10**. PERMANOVA pairwise comparisons for multivariate PPG arsenal at the coastal site, Bodega Marine Reserve. Model: PPG Arsenal = Hormone treatment + (1|Plot). *P*-values < 0.05 in bold. Contrasts are structured: Ecotype, Hormone treatment vs Ecotype, Hormone treatment. For hormone treatment: control = no-hormone, GA = $GA_3$ treatment, MeJA = methyl jasmonate treatment, and Paclo = paclobutrazol.

| contrast | *df* | SS | $R^2$ | **Pseudo *F*** | ***p* (Perm)** |
|---|---|---|---|---|---|
| Perennial Control - Perennial GA | 1 | 0.097 | 0.045 | 4.380 | **0.008** |
| Perennial Control - Perennial Paclo | 1 | 0.011 | 0.004 | 0.437 | **0.001** |
| Perennial Control - Perennial MeJA | 1 | 0.002 | 0.001 | 0.063 | 0.898 |
| Perennial GA - Perennial Paclo | 1 | 0.051 | 0.024 | 2.199 | **0.053** |
| Perennial GA - Perennial MeJA | 1 | 0.118 | 0.048 | 4.633 | **0.001** |
| Perennial Paclo - Perennial MeJA | 1 | 0.020 | 0.006 | 0.736 | 0.238 |

**Table S11**. PERMANOVA pairwise comparisons for multivariate PPG arsenal at the inland site, Pepperwood Preserve. Model: PPG Arsenal = Ecotype + Hormone treatment + Exclosure Treatment + Ecotype x Hormone + (1|Plot). *P*-values < 0.05 in bold. Contrasts are structured: Ecotype, Hormone treatment vs Ecotype, Hormone treatment. For hormone treatment: control = no-hormone, GA = $GA_3$ treatment, MeJA = methyl jasmonate treatment, and Paclo = paclobutrazol.

| contrast | *df* | SS | $R^2$ | **Pseudo *F*** | ***p* (Perm)** |
|---|---|---|---|---|---|
| Perennial Control - Perennial GA | 1 | 0.061 | 0.013 | 2.334 | 0.075 |
| Perennial Paclo - Perennial Control | 1 | 0.008 | 0.002 | 0.317 | 0.701 |
| Perennial MeJA - Perennial Control | 1 | 0.025 | 0.005 | 1.034 | 0.429 |
| Perennial Paclo - Perennial GA | 1 | 0.039 | 0.008 | 1.306 | 0.229 |
| Perennial MeJA - Perennial GA | 1 | 0.113 | 0.022 | 3.896 | 0.111 |
| Perennial MeJA - Perennial Paclo | 1 | 0.030 | 0.005 | 1.105 | 0.288 |
| Annual GA - Annual Control | 1 | 0.273 | 0.076 | 7.908 | **<0.001** |
| Annual Paclo - Annual Control | 1 | 0.011 | 0.003 | 0.301 | 0.655 |
| Annual MeJA - Annual Control | 1 | 0.261 | 0.062 | 7.359 | **<0.001** |
| Annual Paclo - Annual GA | 1 | 0.267 | 0.074 | 7.949 | **<0.001** |
| Annual MeJA - Annual GA | 1 | 0.107 | 0.033 | 3.404 | **0.017** |
| Annual MeJA - Annual Paclo | 1 | 0.276 | 0.065 | 7.970 | **<0.001** |
| Annual Control - Perennial Control | 1 | 5.325 | 0.550 | 194.100 | **<0.001** |
| Annual GA - Perennial GA | 1 | 2.917 | 0.445 | 93.102 | **<0.001** |
| Annual Paclo - Perennial Paclo | 1 | 5.298 | 0.524 | 169.840 | **<0.001** |
| Annual MeJA - Perennial MeJA | 1 | 3.090 | 0.400 | 107.400 | **<0.001** |
| Annual MeJA - Perennial Paclo | 1 | 3.226 | 0.410 | 108.300 | **<0.001** |
| Annual MeJA - Perennial Control | 1 | 3.401 | 0.447 | 132.360 | **<0.001** |
| Annual MeJA - Perennial GA | 1 | 3.122 | 0.423 | 96.162 | **<0.001** |
| Annual Paclo - Perennial MeJA | 1 | 5.207 | 0.521 | 172.900 | **<0.001** |
| Annual Paclo - Perennial Control | 1 | 5.582 | 0.561 | 206.890 | **<0.001** |
| Annual Paclo - Perennial GA | 1 | 4.936 | 0.528 | 144.370 | **<0.001** |

| | | | | | |
|---|---|---|---|---|---|
| Perennial MeJA - Annual GA | 1 | 3.065 | 0.433 | 111.520 | **<0.001** |
| Perennial MeJA - Annual Control | 1 | 4.959 | 0.509 | 161.830 | **<0.001** |
| Annual GA - Perennial Paclo | 1 | 3.299 | 0.479 | 136.750 | **<0.001** |
| Annual GA - Perennial Control | 1 | 3.299 | 0.479 | 136.750 | **<0.001** |
| Annual Control - Perennial Paclo | 1 | 5.055 | 0.513 | 159.180 | **<0.001** |
| Annual Control - Perennial GA | 1 | 4.755 | 0.519 | 136.100 | **<0.001** |

**Table S12**. Tukey post-hoc contrasts for survival at the coastal site, Bodega Marine Reserve. Minimum adequate model: Survival = Ecotype + Hormone Treatment + Exclosure Treatment + Ecotype x Exclosure + Ecotype x Hormone + Hormone x Exclosure + Ecotype x Hormone x Exclosure + (1|maternal family) + (1|Plot). *P*-values < 0.05 in bold. Contrasts are structured: Ecotype, Hormone treatment, Exclosure treatment vs Ecotype, Hormone treatment, Exclosure treatment. For hormone treatment: control = no-hormone, GA = $GA_3$ treatment, MeJA = methyl jasmonate treatment, and Paclo = paclobutrazol. For exclosure treatment: control = open structures, exclosures = mesh exclosures.

| **contrast** | **estimate** | **SE** | ***df*** | ***z*-ratio** | ***p*-value** |
|---|---|---|---|---|---|
| Annual Control Control - Perennial Control Control | 4.850 | 0.521 | Inf | 9.314 | < **0.001** |
| Annual Control Control - Annual GA Control | -1.348 | 0.173 | Inf | -7.789 | < **0.001** |
| Annual Control Control - Perennial GA Control | 1.276 | 0.196 | Inf | 6.525 | < **0.001** |
| Annual Control Control - Annual Paclo Control | 0.013 | 0.167 | Inf | 0.079 | 1.000 |
| Annual Control Control - Perennial Paclo Control | 4.130 | 0.382 | Inf | 10.800 | < **0.001** |
| Annual Control Control - Annual MeJA Control | -0.207 | 0.169 | Inf | -1.227 | 0.998 |
| Annual Control Control - Perennial MeJA Control | 3.891 | 0.348 | Inf | 11.187 | < **0.001** |
| Annual Control Control - Annual Control Exclosure | 0.741 | 0.369 | Inf | 2.011 | 0.819 |
| Annual Control Control - Perennial Control Exclosure | 4.709 | 0.675 | Inf | 6.973 | < **0.001** |
| Annual Control Control - Annual GA Exclosure | -0.698 | 0.369 | Inf | -1.893 | 0.880 |
| Annual Control Control - Perennial GA Exclosure | 2.216 | 0.410 | Inf | 5.400 | < **0.001** |
| Annual Control Control - Annual Paclo Exclosure | 0.997 | 0.369 | Inf | 2.702 | 0.329 |
| Annual Control Control - Perennial Paclo Exclosure | 3.589 | 0.483 | Inf | 7.424 | < **0.001** |
| Annual Control Control - Annual MeJA Exclosure | 0.587 | 0.367 | Inf | 1.597 | 0.969 |
| Annual Control Control - Perennial MeJA Exclosure | 3.462 | 0.471 | Inf | 7.346 | < **0.001** |
| Perennial Control Control - Annual GA Control | -6.198 | 0.525 | Inf | -11.810 | < **0.001** |
| Perennial Control Control - Perennial GA Control | -3.575 | 0.520 | Inf | -6.879 | < **0.001** |
| Perennial Control Control - Annual Paclo Control | -4.837 | 0.522 | Inf | -9.269 | < **0.001** |
| Perennial Control Control - Perennial Paclo Control | -0.721 | 0.612 | Inf | -1.177 | 0.999 |
| Perennial Control Control - Annual MeJA Control | -5.058 | 0.521 | Inf | -9.709 | < **0.001** |
| Perennial Control Control - Perennial MeJA Control | -0.960 | 0.592 | Inf | -1.622 | 0.965 |
| Perennial Control Control - Annual Control Exclosure | -4.109 | 0.614 | Inf | -6.687 | < **0.001** |
| Perennial Control Control - Perennial Control Exclosure | -0.142 | 0.828 | Inf | -0.171 | 1.000 |

| | | | | | |
|---|---|---|---|---|---|
| Perennial Control Control - Annual GA Exclosure | -5.549 | 0.618 | Inf | -8.974 | < **0.001** |
| Perennial Control Control - Perennial GA Exclosure | -2.635 | 0.632 | Inf | -4.169 | 0.003 |
| Perennial Control Control - Annual Paclo Exclosure | -3.854 | 0.614 | Inf | -6.275 | < **0.001** |
| Perennial Control Control - Perennial Paclo Exclosure | -1.262 | 0.680 | Inf | -1.854 | 0.896 |
| Perennial Control Control - Annual MeJA Exclosure | -4.264 | 0.615 | Inf | -6.934 | < **0.001** |
| Perennial Control Control - Perennial MeJA Exclosure | -1.388 | 0.672 | Inf | -2.066 | 0.787 |
| Annual GA Control - Perennial GA Control | 2.624 | 0.204 | Inf | 12.840 | < **0.001** |
| Annual GA Control - Annual Paclo Control | 1.361 | 0.171 | Inf | 7.941 | < **0.001** |
| Annual GA Control - Perennial Paclo Control | 5.478 | 0.388 | Inf | 14.134 | < **0.001** |
| Annual GA Control - Annual MeJA Control | 1.141 | 0.174 | Inf | 6.559 | < **0.001** |
| Annual GA Control - Perennial MeJA Control | 5.239 | 0.354 | Inf | 14.813 | < **0.001** |
| Annual GA Control - Annual Control Exclosure | 2.089 | 0.372 | Inf | 5.613 | < **0.001** |
| Annual GA Control - Perennial Control Exclosure | 6.057 | 0.678 | Inf | 8.939 | < **0.001** |
| Annual GA Control - Annual GA Exclosure | 0.650 | 0.369 | Inf | 1.759 | 0.931 |
| Annual GA Control - Perennial GA Exclosure | 3.564 | 0.414 | Inf | 8.617 | < **0.001** |
| Annual GA Control - Annual Paclo Exclosure | 2.345 | 0.373 | Inf | 6.283 | < **0.001** |
| Annual GA Control - Perennial Paclo Exclosure | 4.937 | 0.487 | Inf | 10.144 | < **0.001** |
| Annual GA Control - Annual MeJA Exclosure | 1.935 | 0.371 | Inf | 5.215 | < **0.001** |
| Annual GA Control - Perennial MeJA Exclosure | 4.810 | 0.475 | Inf | 10.135 | < **0.001** |
| Perennial GA Control - Annual Paclo Control | -1.263 | 0.198 | Inf | -6.383 | < **0.001** |
| Perennial GA Control - Perennial Paclo Control | 2.854 | 0.381 | Inf | 7.492 | < **0.001** |
| Perennial GA Control - Annual MeJA Control | -1.483 | 0.196 | Inf | -7.572 | < **0.001** |
| Perennial GA Control - Perennial MeJA Control | 2.615 | 0.346 | Inf | 7.547 | < **0.001** |
| Perennial GA Control - Annual Control Exclosure | -0.535 | 0.381 | Inf | -1.404 | 0.991 |
| Perennial GA Control - Perennial Control Exclosure | 3.433 | 0.674 | Inf | 5.090 | < **0.001** |
| Perennial GA Control - Annual GA Exclosure | -1.974 | 0.385 | Inf | -5.126 | < **0.001** |
| Perennial GA Control - Perennial GA Exclosure | 0.940 | 0.410 | Inf | 2.290 | 0.631 |
| Perennial GA Control - Annual Paclo Exclosure | -0.279 | 0.381 | Inf | -0.733 | 1.000 |
| Perennial GA Control - Perennial Paclo Exclosure | 2.313 | 0.482 | Inf | 4.795 | < **0.001** |
| Perennial GA Control - Annual MeJA Exclosure | -0.689 | 0.381 | Inf | -1.809 | 0.914 |
| Perennial GA Control - Perennial MeJA Exclosure | 2.187 | 0.470 | Inf | 4.649 | < **0.001** |
| Annual Paclo Control - Perennial Paclo Control | 4.117 | 0.384 | Inf | 10.732 | < **0.001** |
| Annual Paclo Control - Annual MeJA Control | -0.220 | 0.166 | Inf | -1.327 | 0.995 |
| Annual Paclo Control - Perennial MeJA Control | 3.878 | 0.349 | Inf | 11.102 | < **0.001** |
| Annual Paclo Control - Annual Control Exclosure | 0.728 | 0.369 | Inf | 1.975 | 0.839 |
| Annual Paclo Control - Perennial Control Exclosure | 4.696 | 0.676 | Inf | 6.947 | < **0.001** |
| Annual Paclo Control - Annual GA Exclosure | -0.711 | 0.368 | Inf | -1.933 | 0.861 |
| Annual Paclo Control - Perennial GA Exclosure | 2.203 | 0.411 | Inf | 5.356 | < **0.001** |
| Annual Paclo Control - Annual Paclo Exclosure | 0.984 | 0.369 | Inf | 2.664 | 0.354 |

| | | | | | |
|---|---|---|---|---|---|
| Annual Paclo Control - Perennial Paclo Exclosure | 3.576 | 0.484 | Inf | 7.381 | < **0.001** |
| Annual Paclo Control - Annual MeJA Exclosure | 0.574 | 0.367 | Inf | 1.563 | 0.975 |
| Annual Paclo Control - Perennial MeJA Exclosure | 3.449 | 0.472 | Inf | 7.302 | < **0.001** |
| Perennial Paclo Control - Annual MeJA Control | -4.337 | 0.382 | Inf | -11.348 | < **0.001** |
| Perennial Paclo Control - Perennial MeJA Control | -0.239 | 0.474 | Inf | -0.504 | 1.000 |
| Perennial Paclo Control - Annual Control Exclosure | -3.388 | 0.502 | Inf | -6.744 | < **0.001** |
| Perennial Paclo Control - Perennial Control Exclosure | 0.579 | 0.748 | Inf | 0.774 | 1.000 |
| Perennial Paclo Control - Annual GA Exclosure | -4.828 | 0.507 | Inf | -9.522 | < **0.001** |
| Perennial Paclo Control - Perennial GA Exclosure | -1.914 | 0.524 | Inf | -3.654 | 0.023 |
| Perennial Paclo Control - Annual Paclo Exclosure | -3.133 | 0.502 | Inf | -6.241 | < **0.001** |
| Perennial Paclo Control - Perennial Paclo Exclosure | -0.541 | 0.581 | Inf | -0.930 | 1.000 |
| Perennial Paclo Control - Annual MeJA Exclosure | -3.543 | 0.503 | Inf | -7.044 | < **0.001** |
| Perennial Paclo Control - Perennial MeJA Exclosure | -0.667 | 0.571 | Inf | -1.168 | 0.999 |
| Annual MeJA Control - Perennial MeJA Control | 4.098 | 0.348 | Inf | 11.781 | < **0.001** |
| Annual MeJA Control - Annual Control Exclosure | 0.949 | 0.369 | Inf | 2.570 | 0.420 |
| Annual MeJA Control - Perennial Control Exclosure | 4.916 | 0.675 | Inf | 7.279 | < **0.001** |
| Annual MeJA Control - Annual GA Exclosure | -0.491 | 0.369 | Inf | -1.331 | 0.995 |
| Annual MeJA Control - Perennial GA Exclosure | 2.423 | 0.411 | Inf | 5.902 | < **0.001** |
| Annual MeJA Control - Annual Paclo Exclosure | 1.204 | 0.370 | Inf | 3.258 | 0.084 |
| Annual MeJA Control - Perennial Paclo Exclosure | 3.796 | 0.484 | Inf | 7.849 | < **0.001** |
| Annual MeJA Control - Annual MeJA Exclosure | 0.794 | 0.368 | Inf | 2.159 | 0.726 |
| Annual MeJA Control - Perennial MeJA Exclosure | 3.670 | 0.472 | Inf | 7.782 | < **0.001** |
| Perennial MeJA Control - Annual Control Exclosure | -3.149 | 0.477 | Inf | -6.607 | < **0.001** |
| Perennial MeJA Control - Perennial Control Exclosure | 0.818 | 0.732 | Inf | 1.119 | 0.999 |
| Perennial MeJA Control - Annual GA Exclosure | -4.589 | 0.482 | Inf | -9.529 | < **0.001** |
| Perennial MeJA Control - Perennial GA Exclosure | -1.675 | 0.499 | Inf | -3.355 | 0.063 |
| Perennial MeJA Control - Annual Paclo Exclosure | -2.894 | 0.476 | Inf | -6.076 | < **0.001** |
| Perennial MeJA Control - Perennial Paclo Exclosure | -0.302 | 0.559 | Inf | -0.539 | 1.000 |
| Perennial MeJA Control - Annual MeJA Exclosure | -3.304 | 0.477 | Inf | -6.923 | < **0.001** |
| Perennial MeJA Control - Perennial MeJA Exclosure | -0.428 | 0.549 | Inf | -0.780 | 1.000 |
| Annual Control Exclosure - Perennial Control Exclosure | 3.968 | 0.600 | Inf | 6.610 | < **0.001** |
| Annual Control Exclosure - Annual GA Exclosure | -1.440 | 0.209 | Inf | -6.897 | < **0.001** |
| Annual Control Exclosure - Perennial GA Exclosure | 1.475 | 0.272 | Inf | 5.431 | < **0.001** |
| Annual Control Exclosure - Annual Paclo Exclosure | 0.256 | 0.203 | Inf | 1.261 | 0.997 |
| Annual Control Exclosure - Perennial Paclo Exclosure | 2.848 | 0.371 | Inf | 7.666 | < **0.001** |
| Annual Control Exclosure - Annual MeJA Exclosure | -0.155 | 0.201 | Inf | -0.769 | 1.000 |
| Annual Control Exclosure - Perennial MeJA Exclosure | 2.721 | 0.356 | Inf | 7.647 | < **0.001** |
| Perennial Control Exclosure - Annual GA Exclosure | -5.407 | 0.605 | Inf | -8.931 | < **0.001** |
| Perennial Control Exclosure - Perennial GA Exclosure | -2.493 | 0.619 | Inf | -4.029 | **0.006** |

| | | | | | |
|---|---|---|---|---|---|
| Perennial Control Exclosure - Annual Paclo Exclosure | -3.712 | 0.599 | Inf | -6.193 | < **0.001** |
| Perennial Control Exclosure - Perennial Paclo Exclosure | -1.120 | 0.667 | Inf | -1.680 | 0.952 |
| Perennial Control Exclosure - Annual MeJA Exclosure | -4.122 | 0.600 | Inf | -6.866 | < **0.001** |
| Perennial Control Exclosure - Perennial MeJA Exclosure | -1.246 | 0.658 | Inf | -1.893 | 0.880 |
| Annual GA Exclosure - Perennial GA Exclosure | 2.914 | 0.276 | Inf | 10.554 | < **0.001** |
| Annual GA Exclosure - Annual Paclo Exclosure | 1.695 | 0.211 | Inf | 8.015 | < **0.001** |
| Annual GA Exclosure - Perennial Paclo Exclosure | 4.287 | 0.380 | Inf | 11.283 | < **0.001** |
| Annual GA Exclosure - Annual MeJA Exclosure | 1.285 | 0.206 | Inf | 6.239 | < **0.001** |
| Annual GA Exclosure - Perennial MeJA Exclosure | 4.161 | 0.364 | Inf | 11.439 | < **0.001** |
| Perennial GA Exclosure - Annual Paclo Exclosure | -1.219 | 0.271 | Inf | -4.506 | < **0.001** |
| Perennial GA Exclosure - Perennial Paclo Exclosure | 1.373 | 0.401 | Inf | 3.424 | 0.050 |
| Perennial GA Exclosure - Annual MeJA Exclosure | -1.629 | 0.272 | Inf | -5.991 | < **0.001** |
| Perennial GA Exclosure - Perennial MeJA Exclosure | 1.246 | 0.386 | Inf | 3.230 | 0.091 |
| Annual Paclo Exclosure - Perennial Paclo Exclosure | 2.592 | 0.370 | Inf | 7.005 | < **0.001** |
| Annual Paclo Exclosure - Annual MeJA Exclosure | -0.410 | 0.201 | Inf | -2.043 | 0.801 |
| Annual Paclo Exclosure - Perennial MeJA Exclosure | 2.466 | 0.354 | Inf | 6.958 | < **0.001** |
| Perennial Paclo Exclosure - Annual MeJA Exclosure | -3.002 | 0.372 | Inf | -8.077 | < **0.001** |
| Perennial Paclo Exclosure - Perennial MeJA Exclosure | -0.127 | 0.460 | Inf | -0.275 | 1.000 |
| Annual MeJA Exclosure - Perennial MeJA Exclosure | 2.876 | 0.356 | Inf | 8.075 | < **0.001** |

**Table S13**. Tukey post-hoc contrasts for survival at the inland site, Pepperwood Preserve. Tukey post-hoc contrasts for survival at the inland site, Pepperwood Preserve. Minimum adequate model: Probability of survival = Ecotype + (1|maternal family) + (1|Plot). *P*-values < 0.05 in bold.

| **contrast** | **estimate** | **SE** | ***df*** | ***z*-ratio** | ***p*-value** |
|---|---|---|---|---|---|
| Annual - Perennial | 1.74 | 0.078 | Inf | 22.344 | **< 0.001** |

**Table S14**. Tukey post-hoc contrasts for the probability of flowering at the coastal site, Bodega Marine Reserve. Minimum adequate model: Probability of flowering = Ecotype + Hormone Treatment + Exclosure Treatment + Ecotype x Exclosure + (1|maternal family) + (1|Plot). *P*-values < 0.05 in bold. Contrasts are structured: Ecotype, Hormone treatment, Exclosure treatment vs Ecotype, Hormone treatment, Exclosure treatment. For hormone treatment: control = no-hormone, GA = $GA_3$ treatment, MeJA = methyl jasmonate treatment, and Paclo = paclobutrazol. For exclosure treatment: control = open structures, exclosures = mesh exclosures.

| **contrast** | **estimate** | **SE** | ***df*** | ***z*-ratio** | ***p*-value** |
|---|---|---|---|---|---|
| Annual Control Control - Perennial Control Control | -0.096 | 0.282 | Inf | -0.340 | 1.000 |
| Annual Control Control - Annual GA Control | 2.425 | 0.276 | Inf | 8.776 | < **0.001** |

| | | | | | |
|---|---|---|---|---|---|
| Annual Control Control - Perennial GA Control | 2.330 | 0.393 | Inf | 5.923 | < **0.001** |
| Annual Control Control - Annual Paclo Control | -0.205 | 0.222 | Inf | -0.922 | 1.000 |
| Annual Control Control - Perennial Paclo Control | -0.301 | 0.359 | Inf | -0.837 | 1.000 |
| Annual Control Control - Annual MeJA Control | 0.461 | 0.226 | Inf | 2.040 | 0.803 |
| Annual Control Control - Perennial MeJA Control | 0.365 | 0.361 | Inf | 1.012 | 1.000 |
| Annual Control Control - Annual Control Exclosure | -2.094 | 0.776 | Inf | -2.699 | 0.331 |
| Annual Control Control - Perennial Control Exclosure | -3.073 | 0.807 | Inf | -3.810 | **0.013** |
| Annual Control Control - Annual GA Exclosure | 0.332 | 0.806 | Inf | 0.412 | 1.000 |
| Annual Control Control - Perennial GA Exclosure | -0.648 | 0.825 | Inf | -0.785 | 1.000 |
| Annual Control Control - Annual Paclo Exclosure | -2.298 | 0.808 | Inf | -2.845 | 0.243 |
| Annual Control Control - Perennial Paclo Exclosure | -3.278 | 0.838 | Inf | -3.911 | **0.009** |
| Annual Control Control - Annual MeJA Exclosure | -1.632 | 0.804 | Inf | -2.029 | 0.809 |
| Annual Control Control - Perennial MeJA Exclosure | -2.612 | 0.833 | Inf | -3.134 | 0.119 |
| Perennial Control Control - Annual GA Control | 2.521 | 0.396 | Inf | 6.364 | < **0.001** |
| Perennial Control Control - Perennial GA Control | 2.425 | 0.276 | Inf | 8.776 | < **0.001** |
| Perennial Control Control - Annual Paclo Control | -0.109 | 0.359 | Inf | -0.304 | 1.000 |
| Perennial Control Control - Perennial Paclo Control | -0.205 | 0.222 | Inf | -0.922 | 1.000 |
| Perennial Control Control - Annual MeJA Control | 0.557 | 0.362 | Inf | 1.540 | 0.978 |
| Perennial Control Control - Perennial MeJA Control | 0.461 | 0.226 | Inf | 2.040 | 0.803 |
| Perennial Control Control - Annual Control Exclosure | -1.998 | 0.797 | Inf | -2.507 | 0.466 |
| Perennial Control Control - Perennial Control Exclosure | -2.977 | 0.785 | Inf | -3.795 | **0.014** |
| Perennial Control Control - Annual GA Exclosure | 0.427 | 0.827 | Inf | 0.517 | 1.000 |
| Perennial Control Control - Perennial GA Exclosure | -0.552 | 0.804 | Inf | -0.686 | 1.000 |
| Perennial Control Control - Annual Paclo Exclosure | -2.203 | 0.828 | Inf | -2.660 | 0.357 |
| Perennial Control Control - Perennial Paclo Exclosure | -3.182 | 0.817 | Inf | -3.896 | **0.010** |
| Perennial Control Control - Annual MeJA Exclosure | -1.537 | 0.825 | Inf | -1.863 | 0.893 |
| Perennial Control Control - Perennial MeJA Exclosure | -2.516 | 0.812 | Inf | -3.098 | 0.131 |
| Annual GA Control - Perennial GA Control | -0.096 | 0.282 | Inf | -0.340 | 1.000 |
| Annual GA Control - Annual Paclo Control | -2.630 | 0.278 | Inf | -9.455 | < **0.001** |
| Annual GA Control - Perennial Paclo Control | -2.726 | 0.398 | Inf | -6.856 | < **0.001** |
| Annual GA Control - Annual MeJA Control | -1.964 | 0.272 | Inf | -7.218 | < **0.001** |
| Annual GA Control - Perennial MeJA Control | -2.060 | 0.393 | Inf | -5.242 | < **0.001** |
| Annual GA Control - Annual Control Exclosure | -4.519 | 0.841 | Inf | -5.375 | < **0.001** |
| Annual GA Control - Perennial Control Exclosure | -5.498 | 0.879 | Inf | -6.253 | < **0.001** |
| Annual GA Control - Annual GA Exclosure | -2.094 | 0.776 | Inf | -2.699 | 0.331 |
| Annual GA Control - Perennial GA Exclosure | -3.073 | 0.807 | Inf | -3.810 | 0.013 |
| Annual GA Control - Annual Paclo Exclosure | -4.724 | 0.842 | Inf | -5.608 | < **0.001** |
| Annual GA Control - Perennial Paclo Exclosure | -5.703 | 0.881 | Inf | -6.472 | < **0.001** |
| Annual GA Control - Annual MeJA Exclosure | -4.058 | 0.836 | Inf | -4.854 | < **0.001** |

| | | | | | |
|---|---|---|---|---|---|
| Annual GA Control - Perennial MeJA Exclosure | -5.037 | 0.874 | Inf | -5.764 | < **0.001** |
| Perennial GA Control - Annual Paclo Control | -2.534 | 0.394 | Inf | -6.426 | < **0.001** |
| Perennial GA Control - Perennial Paclo Control | -2.630 | 0.278 | Inf | -9.455 | < **0.001** |
| Perennial GA Control - Annual MeJA Control | -1.868 | 0.391 | Inf | -4.784 | < **0.001** |
| Perennial GA Control - Perennial MeJA Control | -1.964 | 0.272 | Inf | -7.218 | < **0.001** |
| Perennial GA Control - Annual Control Exclosure | -4.423 | 0.860 | Inf | -5.145 | < **0.001** |
| Perennial GA Control - Perennial Control Exclosure | -5.403 | 0.858 | Inf | -6.294 | < **0.001** |
| Perennial GA Control - Annual GA Exclosure | -1.998 | 0.797 | Inf | -2.507 | 0.466 |
| Perennial GA Control - Perennial GA Exclosure | -2.977 | 0.785 | Inf | -3.795 | **0.014** |
| Perennial GA Control - Annual Paclo Exclosure | -4.628 | 0.861 | Inf | -5.375 | < **0.001** |
| Perennial GA Control - Perennial Paclo Exclosure | -5.608 | 0.860 | Inf | -6.518 | < **0.001** |
| Perennial GA Control - Annual MeJA Exclosure | -3.962 | 0.855 | Inf | -4.634 | < **0.001** |
| Perennial GA Control - Perennial MeJA Exclosure | -4.941 | 0.853 | Inf | -5.793 | < **0.001** |
| Annual Paclo Control - Perennial Paclo Control | -0.096 | 0.282 | Inf | -0.340 | 1.000 |
| Annual Paclo Control - Annual MeJA Control | 0.666 | 0.226 | Inf | 2.948 | 0.192 |
| Annual Paclo Control - Perennial MeJA Control | 0.570 | 0.361 | Inf | 1.581 | 0.972 |
| Annual Paclo Control - Annual Control Exclosure | -1.889 | 0.806 | Inf | -2.343 | 0.591 |
| Annual Paclo Control - Perennial Control Exclosure | -2.868 | 0.835 | Inf | -3.434 | **0.049** |
| Annual Paclo Control - Annual GA Exclosure | 0.537 | 0.806 | Inf | 0.666 | 1.000 |
| Annual Paclo Control - Perennial GA Exclosure | -0.443 | 0.824 | Inf | -0.537 | 1.000 |
| Annual Paclo Control - Annual Paclo Exclosure | -2.094 | 0.776 | Inf | -2.699 | 0.331 |
| Annual Paclo Control - Perennial Paclo Exclosure | -3.073 | 0.807 | Inf | -3.810 | **0.013** |
| Annual Paclo Control - Annual MeJA Exclosure | -1.428 | 0.803 | Inf | -1.777 | 0.925 |
| Annual Paclo Control - Perennial MeJA Exclosure | -2.407 | 0.832 | Inf | -2.893 | 0.218 |
| Perennial Paclo Control - Annual MeJA Control | 0.762 | 0.362 | Inf | 2.106 | 0.762 |
| Perennial Paclo Control - Perennial MeJA Control | 0.666 | 0.226 | Inf | 2.948 | 0.192 |
| Perennial Paclo Control - Annual Control Exclosure | -1.793 | 0.826 | Inf | -2.170 | 0.719 |
| Perennial Paclo Control - Perennial Control Exclosure | -2.772 | 0.814 | Inf | -3.406 | 0.053 |
| Perennial Paclo Control - Annual GA Exclosure | 0.632 | 0.827 | Inf | 0.765 | 1.000 |
| Perennial Paclo Control - Perennial GA Exclosure | -0.347 | 0.803 | Inf | -0.432 | 1.000 |
| Perennial Paclo Control - Annual Paclo Exclosure | -1.998 | 0.797 | Inf | -2.507 | 0.466 |
| Perennial Paclo Control - Perennial Paclo Exclosure | -2.977 | 0.785 | Inf | -3.795 | **0.014** |
| Perennial Paclo Control - Annual MeJA Exclosure | -1.332 | 0.824 | Inf | -1.616 | 0.966 |
| Perennial Paclo Control - Perennial MeJA Exclosure | -2.311 | 0.811 | Inf | -2.851 | 0.240 |
| Annual MeJA Control - Perennial MeJA Control | -0.096 | 0.282 | Inf | -0.340 | 1.000 |
| Annual MeJA Control - Annual Control Exclosure | -2.555 | 0.812 | Inf | -3.148 | 0.114 |
| Annual MeJA Control - Perennial Control Exclosure | -3.534 | 0.842 | Inf | -4.197 | **0.003** |
| Annual MeJA Control - Annual GA Exclosure | -0.129 | 0.808 | Inf | -0.160 | 1.000 |
| Annual MeJA Control - Perennial GA Exclosure | -1.109 | 0.828 | Inf | -1.339 | 0.994 |

| | | | | | |
|---|---|---|---|---|---|
| Annual MeJA Control - Annual Paclo Exclosure | -2.760 | 0.813 | Inf | -3.396 | 0.055 |
| Annual MeJA Control - Perennial Paclo Exclosure | -3.739 | 0.843 | Inf | -4.433 | **0.001** |
| Annual MeJA Control - Annual MeJA Exclosure | -2.094 | 0.776 | Inf | -2.699 | 0.331 |
| Annual MeJA Control - Perennial MeJA Exclosure | -3.073 | 0.807 | Inf | -3.810 | **0.013** |
| Perennial MeJA Control - Annual Control Exclosure | -2.459 | 0.832 | Inf | -2.957 | 0.188 |
| Perennial MeJA Control - Perennial Control Exclosure | -3.438 | 0.821 | Inf | -4.189 | **0.003** |
| Perennial MeJA Control - Annual GA Exclosure | -0.034 | 0.829 | Inf | -0.041 | 1.000 |
| Perennial MeJA Control - Perennial GA Exclosure | -1.013 | 0.807 | Inf | -1.255 | 0.997 |
| Perennial MeJA Control - Annual Paclo Exclosure | -2.664 | 0.833 | Inf | -3.200 | 0.099 |
| Perennial MeJA Control - Perennial Paclo Exclosure | -3.643 | 0.822 | Inf | -4.432 | **0.001** |
| Perennial MeJA Control - Annual MeJA Exclosure | -1.998 | 0.797 | Inf | -2.507 | 0.466 |
| Perennial MeJA Control - Perennial MeJA Exclosure | -2.977 | 0.785 | Inf | -3.795 | **0.014** |
| Annual Control Exclosure - Perennial Control Exclosure | -0.980 | 0.328 | Inf | -2.984 | 0.176 |
| Annual Control Exclosure - Annual GA Exclosure | 2.425 | 0.276 | Inf | 8.776 | < **0.001** |
| Annual Control Exclosure - Perennial GA Exclosure | 1.446 | 0.408 | Inf | 3.540 | **0.034** |
| Annual Control Exclosure - Annual Paclo Exclosure | -0.205 | 0.222 | Inf | -0.922 | 1.000 |
| Annual Control Exclosure - Perennial Paclo Exclosure | -1.184 | 0.397 | Inf | -2.980 | 0.177 |
| Annual Control Exclosure - Annual MeJA Exclosure | 0.461 | 0.226 | Inf | 2.040 | 0.803 |
| Annual Control Exclosure - Perennial MeJA Exclosure | -0.518 | 0.397 | Inf | -1.306 | 0.996 |
| Perennial Control Exclosure - Annual GA Exclosure | 3.405 | 0.449 | Inf | 7.584 | < **0.001** |
| Perennial Control Exclosure - Perennial GA Exclosure | 2.425 | 0.276 | Inf | 8.776 | < **0.001** |
| Perennial Control Exclosure - Annual Paclo Exclosure | 0.775 | 0.395 | Inf | 1.959 | 0.848 |
| Perennial Control Exclosure - Perennial Paclo Exclosure | -0.205 | 0.222 | Inf | -0.922 | 1.000 |
| Perennial Control Exclosure - Annual MeJA Exclosure | 1.441 | 0.400 | Inf | 3.599 | **0.028** |
| Perennial Control Exclosure - Perennial MeJA Exclosure | 0.461 | 0.226 | Inf | 2.040 | 0.803 |
| Annual GA Exclosure - Perennial GA Exclosure | -0.980 | 0.328 | Inf | -2.984 | 0.176 |
| Annual GA Exclosure - Annual Paclo Exclosure | -2.630 | 0.278 | Inf | -9.455 | < **0.001** |
| Annual GA Exclosure - Perennial Paclo Exclosure | -3.610 | 0.451 | Inf | -8.004 | < **0.001** |
| Annual GA Exclosure - Annual MeJA Exclosure | -1.964 | 0.272 | Inf | -7.218 | < **0.001** |
| Annual GA Exclosure - Perennial MeJA Exclosure | -2.944 | 0.445 | Inf | -6.619 | < **0.001** |
| Perennial GA Exclosure - Annual Paclo Exclosure | -1.651 | 0.409 | Inf | -4.040 | **0.005** |
| Perennial GA Exclosure - Perennial Paclo Exclosure | -2.630 | 0.278 | Inf | -9.455 | < **0.001** |
| Perennial GA Exclosure - Annual MeJA Exclosure | -0.985 | 0.407 | Inf | -2.418 | 0.534 |
| Perennial GA Exclosure - Perennial MeJA Exclosure | -1.964 | 0.272 | Inf | -7.218 | < **0.001** |
| Annual Paclo Exclosure - Perennial Paclo Exclosure | -0.980 | 0.328 | Inf | -2.984 | 0.176 |
| Annual Paclo Exclosure - Annual MeJA Exclosure | 0.666 | 0.226 | Inf | 2.948 | 0.192 |

| | | | | | |
|---|---|---|---|---|---|
| Annual Paclo Exclosure - Perennial MeJA Exclosure | -0.313 | 0.396 | Inf | -0.792 | 1.000 |
| Perennial Paclo Exclosure - Annual MeJA Exclosure | 1.646 | 0.401 | Inf | 4.100 | **0.004** |
| Perennial Paclo Exclosure - Perennial MeJA Exclosure | 0.666 | 0.226 | Inf | 2.948 | 0.192 |
| Annual MeJA Exclosure - Perennial MeJA Exclosure | -0.980 | 0.328 | Inf | -2.984 | 0.176 |

**Table S15**. Tukey post-hoc contrasts for the probability of flowering at the inland site, Pepperwood Preserve. Minimum adequate model: Probability of flowering = Ecotype + Hormone treatment +Exclosure type + Ecotype x Hormone treatment + Ecotype x Exclosure type + (1|maternal family) + (1|Plot). *P*-values < 0.05 in bold. Contrasts are structured: Ecotype, Hormone treatment, Exclosure treatment vs Ecotype, Hormone treatment, Exclosure treatment. For hormone treatment: control = no-hormone, GA = $GA_3$ treatment, MeJA = methyl jasmonate treatment, and Paclo = paclobutrazol. For exclosure treatment: control = open structures, exclosures = mesh exclosures.

| contrast | estimate | SE | *df* | *z*-ratio | *p*-value |
|---|---|---|---|---|---|
| Annual Control Control - Perennial Control Control | 2.270 | 0.474 | Inf | 4.789 | **< 0.001** |
| Annual Control Control - Annual GA Control | -0.189 | 0.601 | Inf | -0.315 | 1.000 |
| Annual Control Control - Perennial GA Control | 3.713 | 0.482 | Inf | 7.697 | **< 0.001** |
| Annual Control Control - Annual Paclo Control | 0.007 | 0.580 | Inf | 0.012 | 1.000 |
| Annual Control Control - Perennial Paclo Control | 2.742 | 0.474 | Inf | 5.789 | **< 0.001** |
| Annual Control Control - Annual MeJA Control | -0.944 | 0.726 | Inf | -1.300 | 0.996 |
| Annual Control Control - Perennial MeJA Control | 3.492 | 0.479 | Inf | 7.287 | **< 0.001** |
| Annual Control Control - Annual Control Exclosure | -19.692 | 5016.600 | Inf | -0.004 | 1.000 |
| Annual Control Control - Perennial Control Exclosure | 0.962 | 0.634 | Inf | 1.519 | 0.981 |
| Annual Control Control - Annual GA Exclosure | -19.882 | 5016.600 | Inf | -0.004 | 1.000 |
| Annual Control Control - Perennial GA Exclosure | 2.405 | 0.627 | Inf | 3.835 | **0.012** |
| Annual Control Control - Annual Paclo Exclosure | -19.685 | 5016.600 | Inf | -0.004 | 1.000 |
| Annual Control Control - Perennial Paclo Exclosure | 1.434 | 0.629 | Inf | 2.278 | 0.640 |
| Annual Control Control - Annual MeJA Exclosure | -20.636 | 5016.600 | Inf | -0.004 | 1.000 |
| Annual Control Control - Perennial MeJA Exclosure | 2.184 | 0.628 | Inf | 3.480 | **0.042** |
| Perennial Control Control - Annual GA Control | -2.459 | 0.499 | Inf | -4.931 | **< 0.001** |
| Perennial Control Control - Perennial GA Control | 1.443 | 0.290 | Inf | 4.969 | **< 0.001** |
| Perennial Control Control - Annual Paclo Control | -2.263 | 0.472 | Inf | -4.797 | **< 0.001** |
| Perennial Control Control - Perennial Paclo Control | 0.472 | 0.286 | Inf | 1.651 | 0.959 |
| Perennial Control Control - Annual MeJA Control | -3.214 | 0.645 | Inf | -4.983 | **< 0.001** |
| Perennial Control Control - Perennial MeJA Control | 1.222 | 0.287 | Inf | 4.253 | **0.002** |
| Perennial Control Control - Annual Control Exclosure | -21.962 | 5016.600 | Inf | -0.004 | 1.000 |
| Perennial Control Control - Perennial Control Exclosure | -1.308 | 0.456 | Inf | -2.865 | 0.233 |
| Perennial Control Control - Annual GA Exclosure | -22.152 | 5016.600 | Inf | -0.004 | 1.000 |
| Perennial Control Control - Perennial GA Exclosure | 0.135 | 0.526 | Inf | 0.256 | 1.000 |

| | | | | | |
|---|---|---|---|---|---|
| Perennial Control Control - Annual Paclo Exclosure | -21.955 | 5016.600 | Inf | -0.004 | 1.000 |
| Perennial Control Control - Perennial Paclo Exclosure | -0.836 | 0.534 | Inf | -1.566 | 0.974 |
| Perennial Control Control - Annual MeJA Exclosure | -22.906 | 5016.600 | Inf | -0.005 | 1.000 |
| Perennial Control Control - Perennial MeJA Exclosure | -0.086 | 0.528 | Inf | -0.163 | 1.000 |
| Annual GA Control - Perennial GA Control | 3.902 | 0.507 | Inf | 7.699 | **< 0.001** |
| Annual GA Control - Annual Paclo Control | 0.196 | 0.600 | Inf | 0.327 | 1.000 |
| Annual GA Control - Perennial Paclo Control | 2.931 | 0.498 | Inf | 5.881 | **< 0.001** |
| Annual GA Control - Annual MeJA Control | -0.754 | 0.742 | Inf | -1.017 | 1.000 |
| Annual GA Control - Perennial MeJA Control | 3.681 | 0.504 | Inf | 7.307 | **< 0.001** |
| Annual GA Control - Annual Control Exclosure | -19.503 | 5016.600 | Inf | -0.004 | 1.000 |
| Annual GA Control - Perennial Control Exclosure | 1.152 | 0.652 | Inf | 1.766 | 0.928 |
| Annual GA Control - Annual GA Exclosure | -19.692 | 5016.600 | Inf | -0.004 | 1.000 |
| Annual GA Control - Perennial GA Exclosure | 2.594 | 0.646 | Inf | 4.016 | **0.006** |
| Annual GA Control - Annual Paclo Exclosure | -19.496 | 5016.600 | Inf | -0.004 | 1.000 |
| Annual GA Control - Perennial Paclo Exclosure | 1.623 | 0.648 | Inf | 2.504 | 0.468 |
| Annual GA Control - Annual MeJA Exclosure | -20.447 | 5016.600 | Inf | -0.004 | 1.000 |
| Annual GA Control - Perennial MeJA Exclosure | 2.374 | 0.646 | Inf | 3.672 | **0.022** |
| Perennial GA Control - Annual Paclo Control | -3.706 | 0.480 | Inf | -7.723 | **< 0.001** |
| Perennial GA Control - Perennial Paclo Control | -0.971 | 0.281 | Inf | -3.459 | **0.045** |
| Perennial GA Control - Annual MeJA Control | -4.656 | 0.651 | Inf | -7.147 | **< 0.001** |
| Perennial GA Control - Perennial MeJA Control | -0.221 | 0.279 | Inf | -0.792 | 1.000 |
| Perennial GA Control - Annual Control Exclosure | -23.405 | 5016.600 | Inf | -0.005 | 1.000 |
| Perennial GA Control - Perennial Control Exclosure | -2.750 | 0.556 | Inf | -4.950 | **< 0.001** |
| Perennial GA Control - Annual GA Exclosure | -23.594 | 5016.600 | Inf | -0.005 | 1.000 |
| Perennial GA Control - Perennial GA Exclosure | -1.308 | 0.456 | Inf | -2.865 | 0.233 |
| Perennial GA Control - Annual Paclo Exclosure | -23.398 | 5016.600 | Inf | -0.005 | 1.000 |
| Perennial GA Control - Perennial Paclo Exclosure | -2.279 | 0.546 | Inf | -4.171 | **0.003** |
| Perennial GA Control - Annual MeJA Exclosure | -24.349 | 5016.600 | Inf | -0.005 | 1.000 |
| Perennial GA Control - Perennial MeJA Exclosure | -1.528 | 0.538 | Inf | -2.840 | 0.246 |
| Annual Paclo Control - Perennial Paclo Control | 2.735 | 0.471 | Inf | 5.805 | **< 0.001** |
| Annual Paclo Control - Annual MeJA Control | -0.951 | 0.725 | Inf | -1.310 | 0.996 |
| Annual Paclo Control - Perennial MeJA Control | 3.485 | 0.477 | Inf | 7.312 | **< 0.001** |
| Annual Paclo Control - Annual Control Exclosure | -19.699 | 5016.600 | Inf | -0.004 | 1.000 |
| Annual Paclo Control - Perennial Control Exclosure | 0.955 | 0.632 | Inf | 1.511 | 0.982 |
| Annual Paclo Control - Annual GA Exclosure | -19.889 | 5016.600 | Inf | -0.004 | 1.000 |
| Annual Paclo Control - Perennial GA Exclosure | 2.398 | 0.626 | Inf | 3.833 | **0.012** |
| Annual Paclo Control - Annual Paclo Exclosure | -19.692 | 5016.600 | Inf | -0.004 | 1.000 |
| Annual Paclo Control - Perennial Paclo Exclosure | 1.427 | 0.628 | Inf | 2.273 | 0.644 |
| Annual Paclo Control - Annual MeJA Exclosure | -20.643 | 5016.600 | Inf | -0.004 | 1.000 |

| | | | | | |
|---|---|---|---|---|---|
| Annual Paclo Control - Perennial MeJA Exclosure | 2.177 | 0.626 | Inf | 3.478 | 0.042 |
| Perennial Paclo Control - Annual MeJA Control | -3.685 | 0.645 | Inf | -5.716 | **< 0.001** |
| Perennial Paclo Control - Perennial MeJA Control | 0.750 | 0.278 | Inf | 2.700 | 0.331 |
| Perennial Paclo Control - Annual Control Exclosure | -22.434 | 5016.600 | Inf | -0.004 | 1.000 |
| Perennial Paclo Control - Perennial Control Exclosure | -1.779 | 0.543 | Inf | -3.278 | 0.079 |
| Perennial Paclo Control - Annual GA Exclosure | -22.624 | 5016.600 | Inf | -0.005 | 1.000 |
| Perennial Paclo Control - Perennial GA Exclosure | -0.337 | 0.525 | Inf | -0.641 | 1.000 |
| Perennial Paclo Control - Annual Paclo Exclosure | -22.427 | 5016.600 | Inf | -0.004 | 1.000 |
| Perennial Paclo Control - Perennial Paclo Exclosure | -1.308 | 0.456 | Inf | -2.865 | 0.233 |
| Perennial Paclo Control - Annual MeJA Exclosure | -23.378 | 5016.600 | Inf | -0.005 | 1.000 |
| Perennial Paclo Control - Perennial MeJA Exclosure | -0.557 | 0.527 | Inf | -1.057 | 1.000 |
| Annual MeJA Control - Perennial MeJA Control | 4.436 | 0.649 | Inf | 6.834 | **< 0.001** |
| Annual MeJA Control - Annual Control Exclosure | -18.749 | 5016.600 | Inf | -0.004 | 1.000 |
| Annual MeJA Control - Perennial Control Exclosure | 1.906 | 0.769 | Inf | 2.477 | 0.489 |
| Annual MeJA Control - Annual GA Exclosure | -18.938 | 5016.600 | Inf | -0.004 | 1.000 |
| Annual MeJA Control - Perennial GA Exclosure | 3.349 | 0.765 | Inf | 4.380 | **0.001** |
| Annual MeJA Control - Annual Paclo Exclosure | -18.742 | 5016.600 | Inf | -0.004 | 1.000 |
| Annual MeJA Control - Perennial Paclo Exclosure | 2.378 | 0.766 | Inf | 3.103 | 0.129 |
| Annual MeJA Control - Annual MeJA Exclosure | -19.692 | 5016.600 | Inf | -0.004 | 1.000 |
| Annual MeJA Control - Perennial MeJA Exclosure | 3.128 | 0.765 | Inf | 4.089 | **0.004** |
| Perennial MeJA Control - Annual Control Exclosure | -23.184 | 5016.600 | Inf | -0.005 | 1.000 |
| Perennial MeJA Control - Perennial Control Exclosure | -2.530 | 0.551 | Inf | -4.593 | **< 0.001** |
| Perennial MeJA Control - Annual GA Exclosure | -23.374 | 5016.600 | Inf | -0.005 | 1.000 |
| Perennial MeJA Control - Perennial GA Exclosure | -1.087 | 0.531 | Inf | -2.046 | 0.799 |
| Perennial MeJA Control - Annual Paclo Exclosure | -23.177 | 5016.600 | Inf | -0.005 | 1.000 |
| Perennial MeJA Control - Perennial Paclo Exclosure | -2.058 | 0.542 | Inf | -3.800 | **0.014** |
| Perennial MeJA Control - Annual MeJA Exclosure | -24.128 | 5016.600 | Inf | -0.005 | 1.000 |
| Perennial MeJA Control - Perennial MeJA Exclosure | -1.308 | 0.456 | Inf | -2.865 | 0.233 |
| Annual Control Exclosure - Perennial Control Exclosure | 20.655 | 5016.600 | Inf | 0.004 | 1.000 |
| Annual Control Exclosure - Annual GA Exclosure | -0.189 | 0.601 | Inf | -0.315 | 1.000 |
| Annual Control Exclosure - Perennial GA Exclosure | 22.097 | 5016.600 | Inf | 0.004 | 1.000 |
| Annual Control Exclosure - Annual Paclo Exclosure | 0.007 | 0.580 | Inf | 0.012 | 1.000 |
| Annual Control Exclosure - Perennial Paclo Exclosure | 21.126 | 5016.600 | Inf | 0.004 | 1.000 |
| Annual Control Exclosure - Annual MeJA Exclosure | -0.944 | 0.726 | Inf | -1.300 | 0.996 |
| Annual Control Exclosure - Perennial MeJA Exclosure | 21.877 | 5016.600 | Inf | 0.004 | 1.000 |
| Perennial Control Exclosure - Annual GA Exclosure | -20.844 | 5016.600 | Inf | -0.004 | 1.000 |
| Perennial Control Exclosure - Perennial GA Exclosure | 1.443 | 0.290 | Inf | 4.969 | **< 0.001** |
| Perennial Control Exclosure - Annual Paclo Exclosure | -20.648 | 5016.600 | Inf | -0.004 | 1.000 |
| Perennial Control Exclosure - Perennial Paclo Exclosure | 0.472 | 0.286 | Inf | 1.651 | 0.959 |

| | | | | | |
|---|---|---|---|---|---|
| Perennial Control Exclosure - Annual MeJA Exclosure | -21.598 | 5016.600 | Inf | -0.004 | 1.000 |
| Perennial Control Exclosure - Perennial MeJA Exclosure | 1.222 | 0.287 | Inf | 4.253 | **0.002** |
| Annual GA Exclosure - Perennial GA Exclosure | 22.287 | 5016.600 | Inf | 0.004 | 1.000 |
| Annual GA Exclosure - Annual Paclo Exclosure | 0.196 | 0.600 | Inf | 0.327 | 1.000 |
| Annual GA Exclosure - Perennial Paclo Exclosure | 21.316 | 5016.600 | Inf | 0.004 | 1.000 |
| Annual GA Exclosure - Annual MeJA Exclosure | -0.754 | 0.742 | Inf | -1.017 | 1.000 |
| Annual GA Exclosure - Perennial MeJA Exclosure | 22.066 | 5016.600 | Inf | 0.004 | 1.000 |
| Perennial GA Exclosure - Annual Paclo Exclosure | -22.090 | 5016.600 | Inf | -0.004 | 1.000 |
| Perennial GA Exclosure - Perennial Paclo Exclosure | -0.971 | 0.281 | Inf | -3.459 | **0.045** |
| Perennial GA Exclosure - Annual MeJA Exclosure | -23.041 | 5016.600 | Inf | -0.005 | 1.000 |
| Perennial GA Exclosure - Perennial MeJA Exclosure | -0.221 | 0.279 | Inf | -0.792 | 1.000 |
| Annual Paclo Exclosure - Perennial Paclo Exclosure | 21.119 | 5016.600 | Inf | 0.004 | 1.000 |
| Annual Paclo Exclosure - Annual MeJA Exclosure | -0.951 | 0.725 | Inf | -1.310 | 0.996 |
| Annual Paclo Exclosure - Perennial MeJA Exclosure | 21.870 | 5016.600 | Inf | 0.004 | 1.000 |
| Perennial Paclo Exclosure - Annual MeJA Exclosure | -22.070 | 5016.600 | Inf | -0.004 | 1.000 |
| Perennial Paclo Exclosure - Perennial MeJA Exclosure | 0.750 | 0.278 | Inf | 2.700 | 0.331 |
| Annual MeJA Exclosure - Perennial MeJA Exclosure | 22.820 | 5016.600 | Inf | 0.005 | 1.000 |

**Table S16**. Tukey post-hoc contrasts for fruit production at the inland site, Pepperwood Preserve. Minimum adequate model: Fruit number (among plants that flowered) = Ecotype + Hormone treatment + Exclosure type + (1|maternal family) + (1|Plot). *P*-values < 0.05 in bold. Contrasts are structured: Ecotype, Hormone treatment, Exclosure treatment vs Ecotype, Hormone treatment, Exclosure treatment. For hormone treatment: control = no-hormone, GA = $GA_3$ treatment, MeJA = methyl jasmonate treatment, and Paclo = paclobutrazol. For exclosure treatment: control = open structures, exclosures = mesh exclosures.

| **contrast** | **estimate** | **SE** | ***df*** | ***z*-ratio** | ***p*-value** |
|---|---|---|---|---|---|
| Annual Control Control - Perennial Control Control | -0.021 | 0.066 | Inf | -0.321 | 1.000 |
| Annual Control Control - Annual GA Control | 0.468 | 0.083 | Inf | 5.640 | **< 0.001** |
| Annual Control Control - Perennial GA Control | 0.447 | 0.112 | Inf | 3.978 | 0.007 |
| Annual Control Control - Annual MeJA Control | 0.234 | 0.080 | Inf | 2.913 | 0.208 |
| Annual Control Control - Perennial MeJA Control | 0.212 | 0.109 | Inf | 1.943 | 0.856 |
| Annual Control Control - Annual Paclo Control | 0.103 | 0.077 | Inf | 1.343 | 0.994 |
| Annual Control Control - Perennial Paclo Control | 0.082 | 0.104 | Inf | 0.789 | 1.000 |
| Annual Control Control - Annual Control Exclosure | -0.577 | 0.305 | Inf | -1.894 | 0.879 |
| Annual Control Control - Perennial Control Exclosure | -0.598 | 0.311 | Inf | -1.925 | 0.865 |
| Annual Control Control - Annual GA Exclosure | -0.109 | 0.316 | Inf | -0.346 | 1.000 |
| Annual Control Control - Perennial GA Exclosure | -0.130 | 0.324 | Inf | -0.403 | 1.000 |
| Annual Control Control - Annual MeJA Exclosure | -0.343 | 0.315 | Inf | -1.088 | 0.999 |

| | | | | | |
|---|---|---|---|---|---|
| Annual Control Control - Perennial MeJA Exclosure | -0.365 | 0.323 | Inf | -1.128 | 0.999 |
| Annual Control Control - Annual Paclo Exclosure | -0.474 | 0.314 | Inf | -1.509 | 0.982 |
| Annual Control Control - Perennial Paclo Exclosure | -0.495 | 0.321 | Inf | -1.544 | 0.977 |
| Perennial Control Control - Annual GA Control | 0.489 | 0.100 | Inf | 4.909 | **< 0.001** |
| Perennial Control Control - Perennial GA Control | 0.468 | 0.083 | Inf | 5.640 | **< 0.001** |
| Perennial Control Control - Annual MeJA Control | 0.255 | 0.098 | Inf | 2.590 | 0.405 |
| Perennial Control Control - Perennial MeJA Control | 0.234 | 0.080 | Inf | 2.913 | 0.208 |
| Perennial Control Control - Annual Paclo Control | 0.124 | 0.099 | Inf | 1.255 | 0.997 |
| Perennial Control Control - Perennial Paclo Control | 0.103 | 0.077 | Inf | 1.343 | 0.994 |
| Perennial Control Control - Annual Control Exclosure | -0.556 | 0.313 | Inf | -1.778 | 0.925 |
| Perennial Control Control - Perennial Control Exclosure | -0.577 | 0.305 | Inf | -1.894 | 0.879 |
| Perennial Control Control - Annual GA Exclosure | -0.088 | 0.321 | Inf | -0.273 | 1.000 |
| Perennial Control Control - Perennial GA Exclosure | -0.109 | 0.316 | Inf | -0.346 | 1.000 |
| Perennial Control Control - Annual MeJA Exclosure | -0.322 | 0.321 | Inf | -1.002 | 1.000 |
| Perennial Control Control - Perennial MeJA Exclosure | -0.343 | 0.315 | Inf | -1.088 | 0.999 |
| Perennial Control Control - Annual Paclo Exclosure | -0.453 | 0.321 | Inf | -1.410 | 0.991 |
| Perennial Control Control - Perennial Paclo Exclosure | -0.474 | 0.314 | Inf | -1.509 | 0.982 |
| Annual GA Control - Perennial GA Control | -0.021 | 0.066 | Inf | -0.321 | 1.000 |
| Annual GA Control - Annual MeJA Control | -0.234 | 0.086 | Inf | -2.708 | 0.325 |
| Annual GA Control - Perennial MeJA Control | -0.255 | 0.108 | Inf | -2.366 | 0.574 |
| Annual GA Control - Annual Paclo Control | -0.365 | 0.084 | Inf | -4.334 | **0.002** |
| Annual GA Control - Perennial Paclo Control | -0.386 | 0.103 | Inf | -3.750 | **0.016** |
| Annual GA Control - Annual Control Exclosure | -1.045 | 0.316 | Inf | -3.311 | 0.071 |
| Annual GA Control - Perennial Control Exclosure | -1.066 | 0.319 | Inf | -3.337 | 0.066 |
| Annual GA Control - Annual GA Exclosure | -0.577 | 0.305 | Inf | -1.894 | 0.879 |
| Annual GA Control - Perennial GA Exclosure | -0.598 | 0.311 | Inf | -1.925 | 0.865 |
| Annual GA Control - Annual MeJA Exclosure | -0.811 | 0.317 | Inf | -2.559 | 0.428 |
| Annual GA Control - Perennial MeJA Exclosure | -0.832 | 0.323 | Inf | -2.580 | 0.413 |
| Annual GA Control - Annual Paclo Exclosure | -0.942 | 0.316 | Inf | -2.982 | 0.177 |
| Annual GA Control - Perennial Paclo Exclosure | -0.963 | 0.321 | Inf | -3.005 | 0.167 |
| Perennial GA Control - Annual MeJA Control | -0.213 | 0.110 | Inf | -1.939 | 0.858 |
| Perennial GA Control - Perennial MeJA Control | -0.234 | 0.086 | Inf | -2.708 | 0.325 |
| Perennial GA Control - Annual Paclo Control | -0.344 | 0.111 | Inf | -3.093 | 0.133 |
| Perennial GA Control - Perennial Paclo Control | -0.365 | 0.084 | Inf | -4.334 | 0.002 |
| Perennial GA Control - Annual Control Exclosure | -1.024 | 0.325 | Inf | -3.146 | 0.115 |
| Perennial GA Control - Perennial Control Exclosure | -1.045 | 0.316 | Inf | -3.311 | 0.071 |
| Perennial GA Control - Annual GA Exclosure | -0.556 | 0.313 | Inf | -1.778 | 0.925 |
| Perennial GA Control - Perennial GA Exclosure | -0.577 | 0.305 | Inf | -1.894 | 0.879 |
| Perennial GA Control - Annual MeJA Exclosure | -0.790 | 0.325 | Inf | -2.431 | 0.524 |

| | | | | | |
|---|---|---|---|---|---|
| Perennial GA Control - Perennial MeJA Exclosure | -0.811 | 0.317 | Inf | -2.559 | 0.428 |
| Perennial GA Control - Annual Paclo Exclosure | -0.921 | 0.325 | Inf | -2.833 | 0.250 |
| Perennial GA Control - Perennial Paclo Exclosure | -0.942 | 0.316 | Inf | -2.982 | 0.177 |
| Annual MeJA Control - Perennial MeJA Control | -0.021 | 0.066 | Inf | -0.321 | 1.000 |
| Annual MeJA Control - Annual Paclo Control | -0.131 | 0.081 | Inf | -1.608 | 0.967 |
| Annual MeJA Control - Perennial Paclo Control | -0.152 | 0.102 | Inf | -1.495 | 0.983 |
| Annual MeJA Control - Annual Control Exclosure | -0.811 | 0.314 | Inf | -2.578 | 0.414 |
| Annual MeJA Control - Perennial Control Exclosure | -0.832 | 0.319 | Inf | -2.610 | 0.391 |
| Annual MeJA Control - Annual GA Exclosure | -0.343 | 0.316 | Inf | -1.084 | 0.999 |
| Annual MeJA Control - Perennial GA Exclosure | -0.364 | 0.323 | Inf | -1.128 | 0.999 |
| Annual MeJA Control - Annual MeJA Exclosure | -0.577 | 0.305 | Inf | -1.894 | 0.879 |
| Annual MeJA Control - Perennial MeJA Exclosure | -0.598 | 0.311 | Inf | -1.925 | 0.865 |
| Annual MeJA Control - Annual Paclo Exclosure | -0.708 | 0.315 | Inf | -2.248 | 0.662 |
| Annual MeJA Control - Perennial Paclo Exclosure | -0.729 | 0.320 | Inf | -2.280 | 0.639 |
| Perennial MeJA Control - Annual Paclo Control | -0.109 | 0.108 | Inf | -1.014 | 1.000 |
| Perennial MeJA Control - Perennial Paclo Control | -0.131 | 0.081 | Inf | -1.608 | 0.967 |
| Perennial MeJA Control - Annual Control Exclosure | -0.789 | 0.324 | Inf | -2.436 | 0.520 |
| Perennial MeJA Control - Perennial Control Exclosure | -0.811 | 0.314 | Inf | -2.578 | 0.414 |
| Perennial MeJA Control - Annual GA Exclosure | -0.322 | 0.324 | Inf | -0.993 | 1.000 |
| Perennial MeJA Control - Perennial GA Exclosure | -0.343 | 0.316 | Inf | -1.084 | 0.999 |
| Perennial MeJA Control - Annual MeJA Exclosure | -0.556 | 0.313 | Inf | -1.778 | 0.925 |
| Perennial MeJA Control - Perennial MeJA Exclosure | -0.577 | 0.305 | Inf | -1.894 | 0.879 |
| Perennial MeJA Control - Annual Paclo Exclosure | -0.686 | 0.324 | Inf | -2.121 | 0.751 |
| Perennial MeJA Control - Perennial Paclo Exclosure | -0.708 | 0.315 | Inf | -2.248 | 0.662 |
| Annual Paclo Control - Perennial Paclo Control | -0.021 | 0.066 | Inf | -0.321 | 1.000 |
| Annual Paclo Control - Annual Control Exclosure | -0.680 | 0.314 | Inf | -2.165 | 0.722 |
| Annual Paclo Control - Perennial Control Exclosure | -0.701 | 0.319 | Inf | -2.196 | 0.701 |
| Annual Paclo Control - Annual GA Exclosure | -0.212 | 0.316 | Inf | -0.671 | 1.000 |
| Annual Paclo Control - Perennial GA Exclosure | -0.233 | 0.323 | Inf | -0.721 | 1.000 |
| Annual Paclo Control - Annual MeJA Exclosure | -0.446 | 0.316 | Inf | -1.414 | 0.990 |
| Annual Paclo Control - Perennial MeJA Exclosure | -0.467 | 0.323 | Inf | -1.448 | 0.988 |
| Annual Paclo Control - Annual Paclo Exclosure | -0.577 | 0.305 | Inf | -1.894 | 0.879 |
| Annual Paclo Control - Perennial Paclo Exclosure | -0.598 | 0.311 | Inf | -1.925 | 0.865 |
| Perennial Paclo Control - Annual Control Exclosure | -0.659 | 0.323 | Inf | -2.042 | 0.801 |
| Perennial Paclo Control - Perennial Control Exclosure | -0.680 | 0.314 | Inf | -2.165 | 0.722 |
| Perennial Paclo Control - Annual GA Exclosure | -0.191 | 0.322 | Inf | -0.592 | 1.000 |
| Perennial Paclo Control - Perennial GA Exclosure | -0.212 | 0.316 | Inf | -0.671 | 1.000 |
| Perennial Paclo Control - Annual MeJA Exclosure | -0.425 | 0.322 | Inf | -1.318 | 0.995 |
| Perennial Paclo Control - Perennial MeJA Exclosure | -0.446 | 0.316 | Inf | -1.414 | 0.990 |

| | | | | | |
|---|---|---|---|---|---|
| Perennial Paclo Control - Annual Paclo Exclosure | -0.556 | 0.313 | Inf | -1.778 | 0.925 |
| Perennial Paclo Control - Perennial Paclo Exclosure | -0.577 | 0.305 | Inf | -1.894 | 0.879 |
| Annual Control Exclosure - Perennial Control Exclosure | -0.021 | 0.066 | Inf | -0.321 | 1.000 |
| Annual Control Exclosure - Annual GA Exclosure | 0.468 | 0.083 | Inf | 5.640 | **< 0.001** |
| Annual Control Exclosure - Perennial GA Exclosure | 0.447 | 0.112 | Inf | 3.978 | **0.007** |
| Annual Control Exclosure - Annual MeJA Exclosure | 0.234 | 0.080 | Inf | 2.913 | 0.208 |
| Annual Control Exclosure - Perennial MeJA Exclosure | 0.212 | 0.109 | Inf | 1.943 | 0.856 |
| Annual Control Exclosure - Annual Paclo Exclosure | 0.103 | 0.077 | Inf | 1.343 | 0.994 |
| Annual Control Exclosure - Perennial Paclo Exclosure | 0.082 | 0.104 | Inf | 0.789 | 1.000 |
| Perennial Control Exclosure - Annual GA Exclosure | 0.489 | 0.100 | Inf | 4.909 | **< 0.001** |
| Perennial Control Exclosure - Perennial GA Exclosure | 0.468 | 0.083 | Inf | 5.640 | **< 0.001** |
| Perennial Control Exclosure - Annual MeJA Exclosure | 0.255 | 0.098 | Inf | 2.590 | 0.405 |
| Perennial Control Exclosure - Perennial MeJA Exclosure | 0.234 | 0.080 | Inf | 2.913 | 0.208 |
| Perennial Control Exclosure - Annual Paclo Exclosure | 0.124 | 0.099 | Inf | 1.255 | 0.997 |
| Perennial Control Exclosure - Perennial Paclo Exclosure | 0.103 | 0.077 | Inf | 1.343 | 0.994 |
| Annual GA Exclosure - Perennial GA Exclosure | -0.021 | 0.066 | Inf | -0.321 | 1.000 |
| Annual GA Exclosure - Annual MeJA Exclosure | -0.234 | 0.086 | Inf | -2.708 | 0.325 |
| Annual GA Exclosure - Perennial MeJA Exclosure | -0.255 | 0.108 | Inf | -2.366 | 0.574 |
| Annual GA Exclosure - Annual Paclo Exclosure | -0.365 | 0.084 | Inf | -4.334 | **0.002** |
| Annual GA Exclosure - Perennial Paclo Exclosure | -0.386 | 0.103 | Inf | -3.750 | **0.016** |
| Perennial GA Exclosure - Annual MeJA Exclosure | -0.213 | 0.110 | Inf | -1.939 | 0.858 |
| Perennial GA Exclosure - Perennial MeJA Exclosure | -0.234 | 0.086 | Inf | -2.708 | 0.325 |
| Perennial GA Exclosure - Annual Paclo Exclosure | -0.344 | 0.111 | Inf | -3.093 | 0.133 |
| Perennial GA Exclosure - Perennial Paclo Exclosure | -0.365 | 0.084 | Inf | -4.334 | 0.002 |
| Annual MeJA Exclosure - Perennial MeJA Exclosure | -0.021 | 0.066 | Inf | -0.321 | 1.000 |
| Annual MeJA Exclosure - Annual Paclo Exclosure | -0.131 | 0.081 | Inf | -1.608 | 0.967 |
| Annual MeJA Exclosure - Perennial Paclo Exclosure | -0.152 | 0.102 | Inf | -1.495 | 0.983 |
| Perennial MeJA Exclosure - Annual Paclo Exclosure | -0.109 | 0.108 | Inf | -1.014 | 1.000 |
| Perennial MeJA Exclosure - Perennial Paclo Exclosure | -0.131 | 0.081 | Inf | -1.608 | 0.967 |
| Annual Paclo Exclosure - Perennial Paclo Exclosure | -0.021 | 0.066 | Inf | -0.321 | 1.000 |

**Table S17**. Localities of Populations used in the 2023 GA experiment.

| Population | Ecotype | Latitude | Longitude |
|---|---|---|---|
| HEC | Coastal Perennial | 44.13506 | -124.1228 |
| OPB | Coastal Perennial | 42.46401 | -124.42291 |

| | | | |
|---|---|---|---|
| SWB | Coastal Perennial | 39.03598 | -123.69046 |
| BMR | Coastal Perennial | 38.31608 | -123.06907 |
| LMC | Inland Annual | 38.86398 | -123.08391 |
| PPW | Inland Annual | 38.5755 | -122.7009 |
| CAV | Inland Annual | 38.34281 | -122.4854 |
| MOR | Inland Annual | 38.42958 | -122.94496 |
| SAL | Inland Perennial | 41.33966 | -123.38816 |
| AJEN | Near-Coastal Annual | 38.45672 | -123.11599 |
| JEN | Near-Coastal Perennial | 38.4677 | -123.12831 |
| GUL | Near-Coastal Perennial | 38.481706 | -123.13299 |

**Table S18**. Pairwise contrasts of height in the greenhouse between control water sprayed vs GA-treated seedlings. *P*-values were adjusted using a multivariate *t*-adjustment for multiple comparisons. *P*-values < 0.05 in bold.

| Population | estimate | SE | *df* | *t*-ratio | *p*-value |
|---|---|---|---|---|---|
| HEC | -11.17 | 1.27 | 511 | -8.767 | < **0.0001** |
| OPB | -7.05 | 1.27 | 511 | -5.537 | < **0.0001** |
| SWB | -7.8 | 1.27 | 511 | -6.127 | < **0.0001** |
| BMR | -10.78 | 1.27 | 511 | -8.467 | < **0.0001** |
| LMC | -19.4 | 1.29 | 511 | -15.046 | < **0.0001** |
| PPW | -8.94 | 1.27 | 511 | -7.019 | < **0.0001** |
| CAV | -25.39 | 1.29 | 511 | -19.694 | < **0.0001** |
| MOR | -19.72 | 1.29 | 511 | -15.299 | < **0.0001** |
| AJEN | -6.98 | 1.27 | 511 | -5.486 | < **0.0001** |
| SAL | -21.57 | 1.27 | 511 | -16.947 | < **0.0001** |

| | | | | | |
|---|---|---|---|---|---|
| JEN | -20.35 | 1.27 | 511 | -15.974 | **< 0.0001** |
| GUL | -2.93 | 1.27 | 511 | -2.3 | 0.2319 |

**Table S19.** Contrasts of logit transformed leaf damage in the ocean-exposed control structures between water sprayed vs GA-treated seedlings. *P*-values were adjusted using a multivariate *t*-adjustment for multiple comparisons. *P*-values < 0.05 in bold.

| Population | estimate | SE | *df* | *t*-ratio | *p*-value |
|---|---|---|---|---|---|
| HEC | -0.653 | 0.427 | 294 | -1.528 | 0.8011 |
| OPB | 0.192 | 0.427 | 294 | 0.449 | 1.0000 |
| SWB | -0.388 | 0.427 | 294 | -0.908 | 0.9954 |
| BMR | -0.317 | 0.427 | 294 | -0.743 | 0.9993 |
| LMC | -1.68 | 0.439 | 294 | -3.83 | **0.0019** |
| PPW | -0.566 | 0.427 | 294 | -1.326 | 0.9125 |
| CAV | -1.323 | 0.439 | 294 | -3.013 | **0.0331** |
| MOR | -1.08 | 0.438 | 294 | -2.468 | 0.1562 |
| AJEN | -0.243 | 0.428 | 294 | -0.568 | 1.0000 |
| SAL | -2.082 | 0.428 | 294 | -4.868 | **< 0.0001** |
| JEN | -1.326 | 0.427 | 294 | -3.103 | **0.0249** |
| GUL | -0.185 | 0.427 | 294 | -0.434 | 1.0000 |

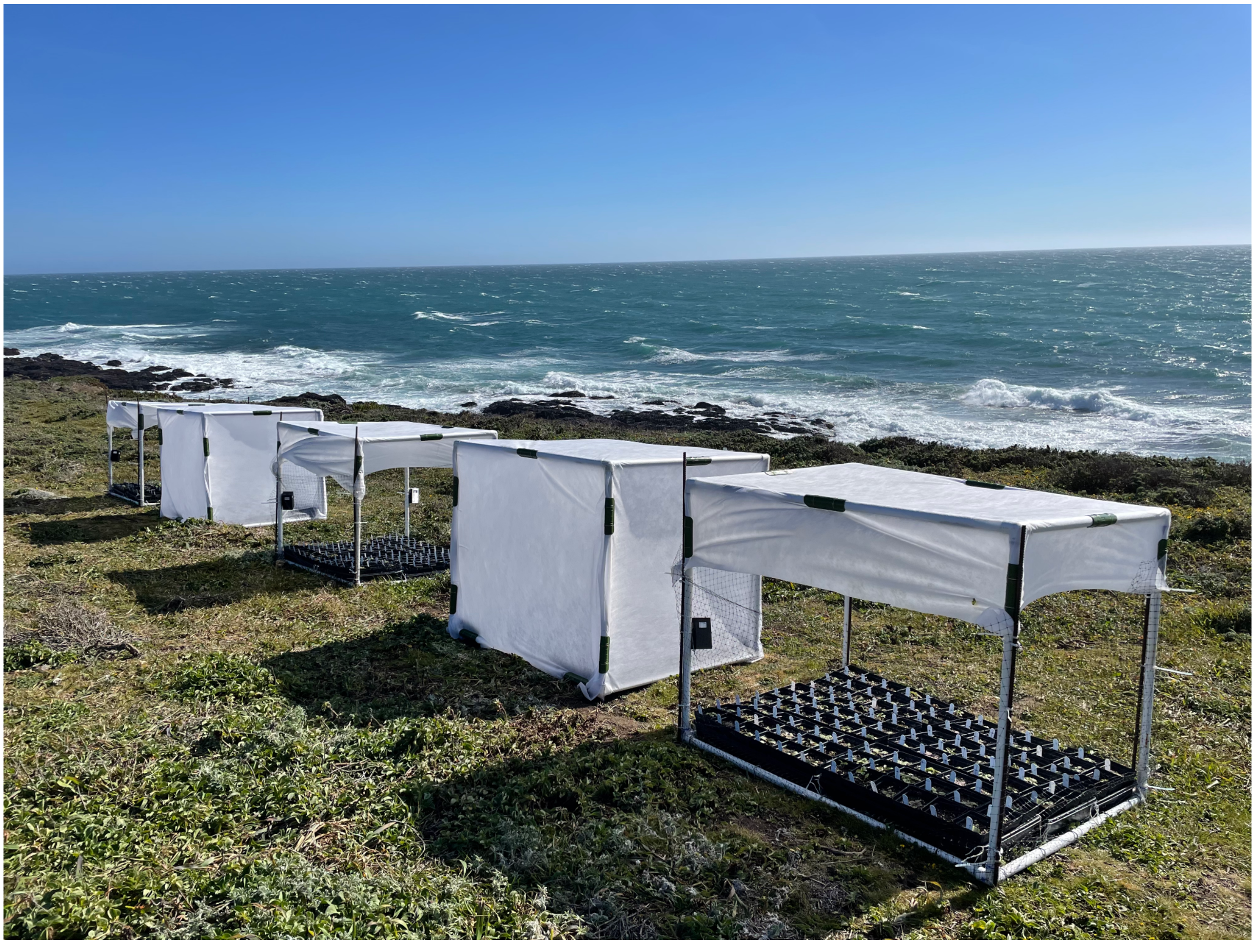

**Figure S1.** Photograph of exclosures and control structures on the coastal bluff in the 2023 GA experiment.

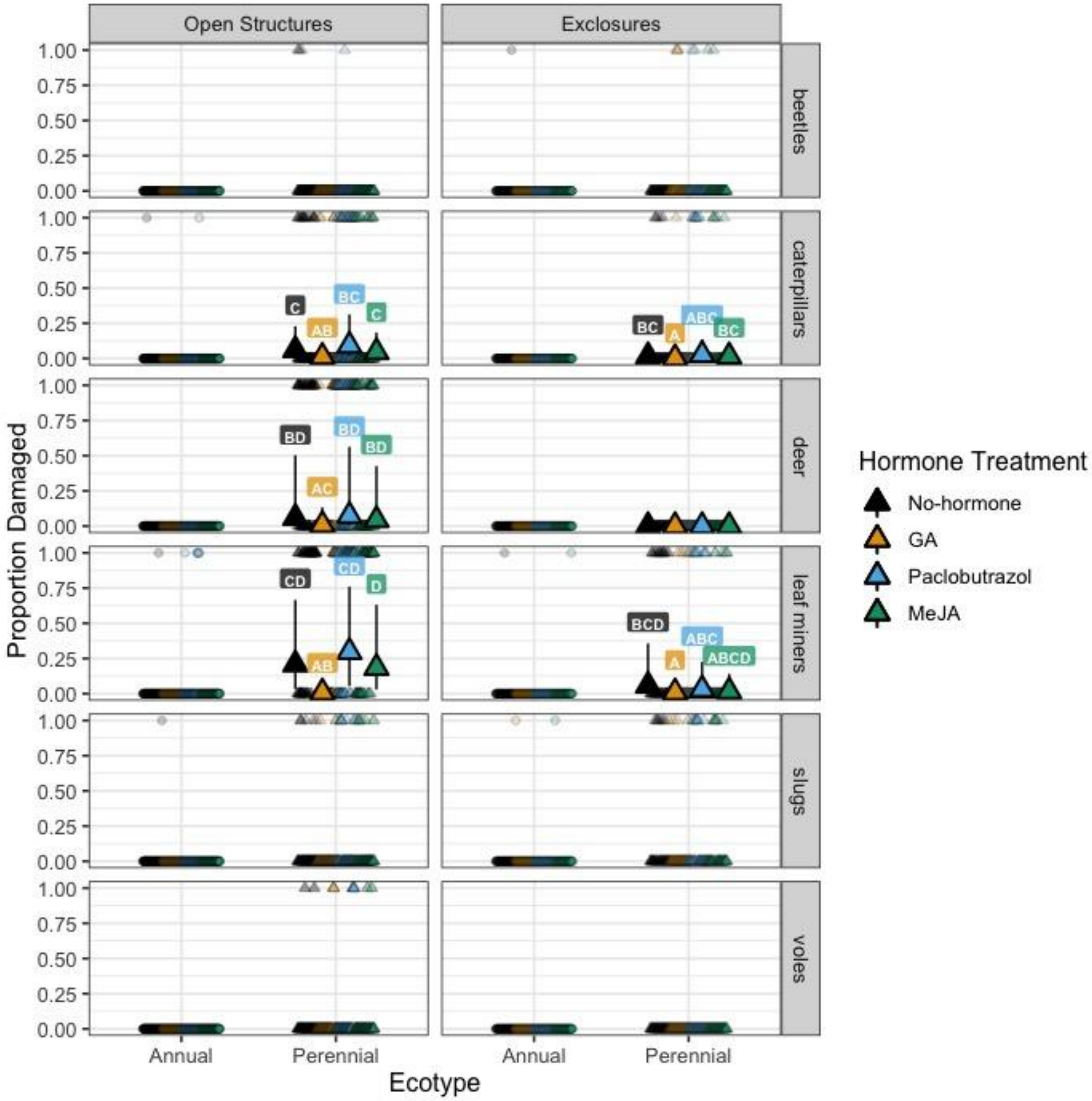


**Figure S2.** The proportion of annuals (circles) and perennials (triangles) damaged by beetles, caterpillars, deer, leaf miners, slugs, and voles treated with gibberellic acid (GA, yellow), paclobutrazol (blue), and methyl jasmonate (MeJA, green), and the no-hormone controls (black) in open structures and herbivore exclosures at the coastal site, Bodega Marine Reserve, in 2020. Larger symbols in the foreground are the mean predictions and 95% confidence intervals from mixed models, smaller and lighter symbols in the background are the raw data. Since no perennials were damaged by deer in exclosures, the binomial model did not accurately estimate that parameter and thus we did not plot those confidence intervals (0-100%). Results of Tukey post-hoc contrasts within each site are indicated above each prediction; shared letters indicate that groups do not significantly differ, while non-overlapping letters indicate that groups significantly differ within each site.

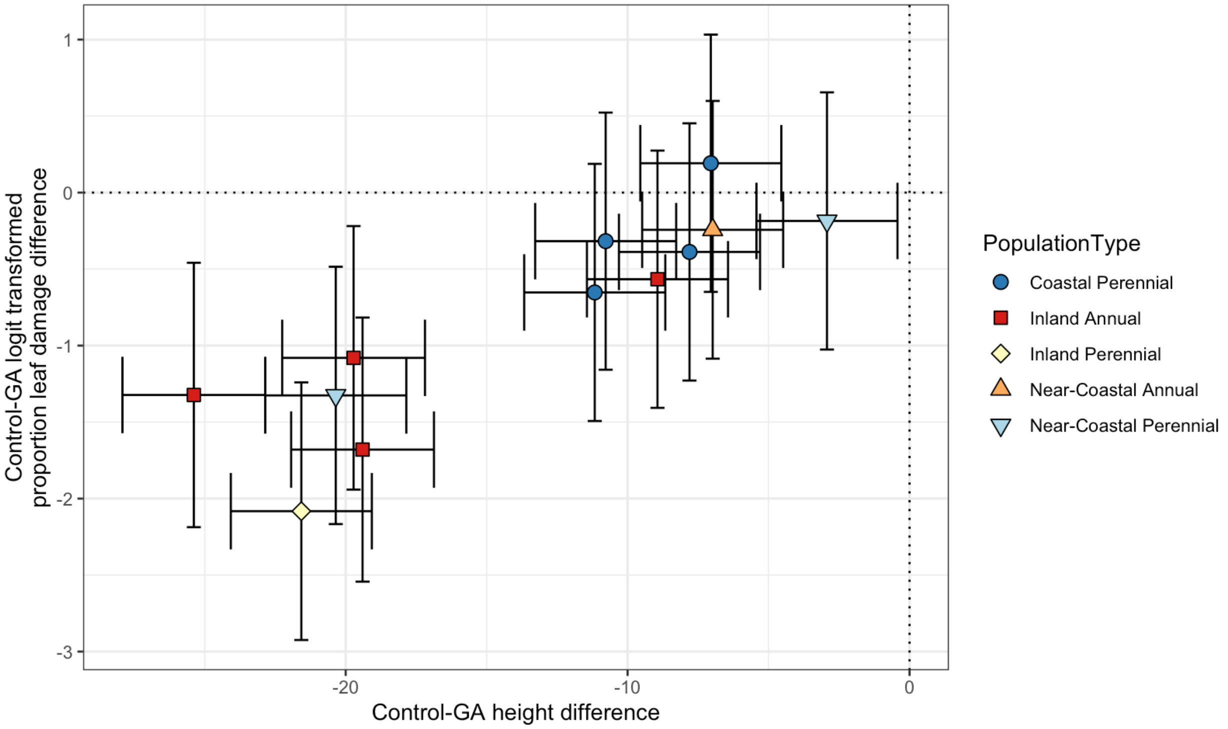


**Figure S3.** Accessions that grew tallest in response to GA treatment in the greenhouse had the most leaf damage after a week of ocean exposure in 2023. Points are Control-GA pairwise contrast estimates (i.e., the difference between the control and GA treatment for each population) for height on the x-axis and logit-transformed proportion leaf damage on the y-axis. The horizontal and vertical error bars are 95% confidence intervals for each estimate. If the confidence intervals overlap with the 0 dotted lines, that means there is no difference between the control and treatment. Negative values indicate that the controls were shorter and had less leaf damage.
